# YAP engages RIF1 to dampen replication stress-induced DNA damage in human squamous cell carcinoma

**DOI:** 10.1101/2024.12.20.629444

**Authors:** Jodie Bojko, Elodie Sins, Benjamin Flynn, James Scarth, Mikal Negasi, Bertram Aschenbrenner, Molly Guscott, Alexander Howard, Emily Lay, Emma Bailey, Natalia Krajic, Marco Franciosi, Sandra Catalán Jiménez, Kelli Gallacher, Ilaria Di Girolamo, Reem Bagabas, Caterina Missero, Jun Wang, Sarah McClelland, Beate M. Lichtenberger, Ute Jungwirth, Gernot Walko

## Abstract

Squamous cell carcinoma cells experience high levels of replication stress due to oncogene-induced cell cycle deregulation. How such cells sustain rapid proliferation despite replication stress is still not fully understood. Here, we discovered, using rapid immunoprecipitation mass spectrometry of endogenous protein (RIME) analysis, that the squamous cell carcinoma oncoprotein YAP engages with RIF1, a key regulator of DNA replication and DNA damage repair under replication stress. RIF1 is highly expressed in human squamous cell carcinoma cell lines and tissues and upregulated during tumour progression. Depletion of RIF1 in squamous cell carcinoma cells exacerbates their endogenous replication stress. Mechanistically, we show that YAP interacts with RIF1 specifically at broken replication forks, and that YAP depletion impairs DNA damage repair under replication stress. Our results thus demonstrate that YAP’s oncogenic functions in squamous cancers involve both transcriptional and non-transcriptional mechanisms, the latter through interaction with RIF1 to dampen replication stress.

## Introduction

Squamous cell carcinomas (SCCs) are among the most common solid tumours [1], with cutaneous SCC (cSCC) being the most prevalent type, imposing a major healthcare burden due to its high incidence [2].

Oncogenic signalling drives cancer cell proliferation by interfering with cell cycle control and upregulation of transcription, thereby causing replication stress, which occurs when DNA replication fork progression in S phase slows or stalls [3–5]. Ineffective resolution of replication stress leads to genomic instability or mitotic catastrophes [3, 4]. Cancer cells with high endogenous replication stress rely on DNA damage repair (DDR) for survival [3]. Of note, stem and progenitor cells in healthy human squamous epithelia have already high levels of endogenous replication stress due to their high proliferation rates, and couple activation of DDR pathways to polyploidisation and terminal differentiation as a homeostatic mechanism [6–8]. The same mechanism operates to eliminate pre-cancerous cells [7–9]. In SCC, this protective mechanism is not fully operational, allowing SCC cells to proliferate despite DNA damage [10].

RIF1 is a key regulator of DNA replication and DDR [11]. By organising higher order chromatin structures into zones of coordinated DNA replication, and through recruitment of protein phosphatase 1 (PP1) to prevent MCM (minichromosome maintenance complex) subunit phosphorylation, RIF1 suppresses late replicating origin firing [12–19]. Under replication stress, RIF1 has two known functions: it protects stalled replication forks from resection [20–22], and in DDR interacts with BRCA1 at broken replication forks [23]. While RIFs role in non-homologous end joining (NHEJ) is well documented [24–27], its role in DNA replication under replication stress is underexplored.

The transcriptional co-regulators Yes-associated protein (YAP) and transcriptional co-activator with PDZ-binding motif (TAZ, also called WW Domain Containing Transcription Regulator 1 (WWTR1)) are essential oncoproteins in SCC [28–30]. YAP/TAZ operate as transcriptional co-activators and co-repressors by indirectly binding to DNA via different transcription factors (TFs; predominantly those of the TEAD family [28]) [31]. While YAP is essential in squamous epithelial stem cell proliferation upon injury [32, 33], overexpression of oncogenic YAP induces rapid senescence [33]. It is likely that this oncogene-induced senescence mechanism is linked to increased YAP-driven transcription, causing transcription-replication conflicts and replication stress [34, 35].

How YAP/TAZ control oncogenic signalling in human SCC at the molecular level, and how human SCC cells deal with oncogene-induced replication stress remains poorly understood. Using a multi-omics approach we characterized transcriptional and non-transcriptional mechanisms of YAP’s oncogenic role in human SCC. We identified RIF1 as a binding partner of YAP in human SCC cells, expanding its known interaction in early amphibian development to human cancers. Importantly, our finding that YAP engages RIF1 under replication stress and facilitates DDR has translational implications, since replication stress is a crucial vulnerability of cancer cells that can be exploited therapeutically.

## Materials and Methods

Complete details of the materials and methods are provided in the supplemental materials and methods.

## Results

### YAP/TAZ-TEAD-mediated oncogenic signalling balances proliferation and differentiation and drives metastatic progression of human squamous cell carcinoma

In healthy epidermis, YAP/TAZ balance cell proliferation and terminal differentiation [32, 36]. To test if YAP/TAZ play similar roles in cSCC, we performed RNAseq analysis on SCC13 cells, a widely used human cSCC cell line [37], following acute siRNA-mediated knockdown of YAP/TAZ (siYAP/TAZ; Supplementary Fig. 1A, B). While we were unable to significantly deplete TAZ (WWTR1), the TAZ-targeting siRNA pool prevented compensatory upregulation of TAZ in response to YAP knockdown (Supplementary Fig. 1B) [32]. DESeq2 analysis revealed a large number of differentially expressed genes (DEGs) upon siYAP/TAZ (977 DEGs downregulated, 2,209 DEGs upregulated; Supplementary Table 1). Downregulated DEGs comprised both, genes positively regulated by YAP/TAZ in cooperation with TEADs [38], and genes regulated by YAP/TAZ in cooperation with the Myb-MuB (MMB) complex [39] (Supplementary Fig. 1C and Supplementary Table 1). Gene set enrichment analysis (GSEA) revealed significant enrichment of pathways associated with cell cycle regulation, DNA replication, and mitosis among the downregulated DEGs, and of pathways associated with keratinization, cornification, extracellular matrix (ECM) remodelling, and immunoregulation among the upregulated DEGs (Supplementary Figs. 1D, E, and Supplementary Table 1). YAP/TAZ knockdown also led to increased terminal differentiation which was evident from (i) enrichment of an epidermal terminal differentiation gene signature [40] in the upregulated DEGs and of an epidermal progenitor cell signature [40] in the downregulated DEGs (Supplementary Fig. 2A), and (ii) upregulation of transcriptional regulators of terminal differentiation and genes belonging to the epidermal differentiation complex (EDC) [41] (Supplementary Fig. 2B and Supplementary Table 1).

We next used the YAP/TAZ target gene expression signature derived from our RNA-seq data to perform gene set variation analysis (GSVA) on one of the largest bulk RNA-seq datasets from patients with cSCC (237 patients) [42]. This revealed that genes that are positively regulated by YAP/TAZ in SCC13 cells are expressed significantly higher in cSCC tumours (as well as their perilesional skin tissue) that have progressed to metastatic disease, compared to those cSCC tumours that have not (Fig. 1A). Genes that were positively regulated by YAP/TAZ in SCC13 cells were also highly expressed in cSCC metastases. Conversely, genes that were negatively regulated by YAP/TAZ in SCC13 cells, including those genes related to terminal differentiation, displayed significantly reduced expression in cSCCs that had progressed to metastatic disease, as well as in the respective metastases (Fig. 1B). This finding provides intriguing evidence in clinical samples for the involvement of YAP/TAZ in the metastatic progression of human cSCC.

**Figure 1.**
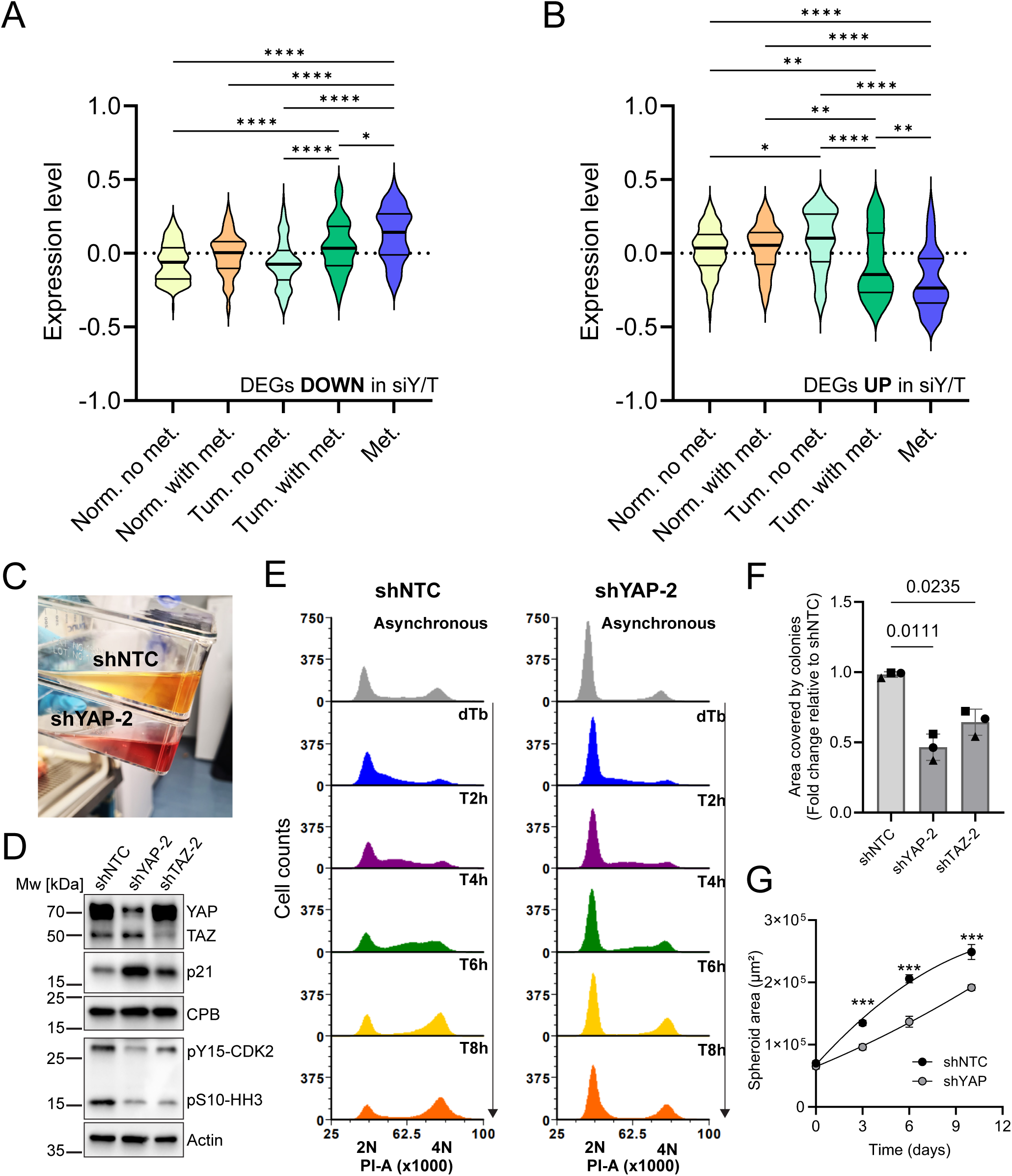
YAP/TAZ-TEAD-mediated oncogenic transcriptional programmes control the balance between proliferation and differentiation in squamous cell carcinoma progression. (A, B) Violin blots (thick line, median; thin lines, upper- and lower quartiles) show the distribution of Gene Set Variation Analysis (GSVA) scores for differentially expressed genes (DEGs) that were (A) downregulated or (B) upregulated in siYAP/TAZ SCC13 cells across different cSCC patient samples [42]. Norm_no_met / Tum_no_met: perilesional normal skin and primary cSCC tissue from patients with no metastasis over 3 years; Norm_with_met / Tum_with_met: perilesional normal skin and primary cSCC tissue from patients with metastasis during 3 years follow-up; Met: tissue from metastasis. Pairwise comparisons between sample groups were conducted using a t test. The p-values from these comparisons are indicated above the boxplots with brackets linking the compared groups. The significance levels are denoted as follows: *p=0.032; **p=0.0021; ***p=0.0002; ****p<0.0001. (C) Photograph shows lack of medium acidification by doxycycline-induced shYAP compared to shNTC SCC13 cells during a 4-day culture period. (D) Immunoblot analysis of SCC13 cells expressing doxycycline-induced shYAP, shTAZ, or shNTC for 72 h, using the indicated antibodies. Actin or cyclophilin B (CBP) were used as loading controls. (E) Cell cycle analysis by propidium iodide (PI) labelling of asynchronous populations of shNTC and shYAP SCC13 cells (grey profiles), and of cells synchronised at the G1/S boundary by double thymidine block (dTb) and released into S phase and analysed at indicated time points (T=hours post thymidine release, coloured profiles). (F) Colony formation assays were quantified by measuring the percentage of well area occupied by colonies. Bars show the means from n=3 independent experiments (performed with 3 technical replicates), individual data points (different shapes indicate different experiments) show the means from each experiment, error bars show SD. Exact p-values are shown, RM one way ANOVA with Geisser-Greenhouse correction and Dunnett’s multiple comparison test. (G) 3D spheroid growth of shNTC and shYAP SCC13 cells. p-values are shown for multiple comparisons at all timepoints, Ordinary two-way ANOVA and Šidák’s multiple comparison test. *p<0.033, **p<0.002, ***p<0.001. Sources of variation: shRNA, p<0.001; time, p<0.001; shRNA x time, p<0.001.

In accordance with the detected transcriptomic changes in siYAP/TAZ SCC13 cells, we observed that cell proliferation was strongly reduced upon expression of doxycycline-inducible YAP-targeting shRNAs (shYAP) (Fig. 1C). This was evident from reduced levels of Ser10-phosphorylated histone H3 (pHH3), a marker of mitotic cells, and strong upregulation of the CDK inhibitor p21 (Fig. 1D). Consistent with p21 upregulation (Fig. 1D, Supplementary Fig. 2C) and the reduced expression of several key regulators of DNA replication (Supplementary Fig. 1C and Supplementary Table 1), most shYAP SCC13 cells arrested in G1, and a much smaller proportion of cells progressed through S phase (Fig. 1E). Compromised cell cycle progression and increased terminal differentiation led to reduced 2D clonal growth (Fig. 1F) and 3D spheroid growth (Fig. 1G) of shYAP SCC13 cells. The effects of shTAZ on SCC13 proliferation were less strong (Figs. 1D, F). To assess if in cSCC cells YAP/TAZ were driving oncogenic transcription through TEAD TFs, we employed the potent allosteric pan-TEAD inhibitor GNE-7883 [43]. GNE-7883 reduced proliferation of SCC13 cells and other cSCC cell lines in dose-dependent manner (Figs. 2A, B). GNE-7883 treatment of SCC13 cells phenocopied shYAP by strongly inducing p21 and the terminal differentiation marker involucrin, and downregulating pHH3 (Figs. 2C, D). GNE-7883 also effectively blocked proliferation of SCC13 cells growing as 3D spheroids (Figs. 2E, F). Consistent with CDKN1A/p21 being a major effector of disrupted YAP/TAZ-TEAD signalling, we found that SCC13 cells expressing CDKN1A-targeting shRNAs were able to proliferate normally upon GNE-7883 treatment (Figs. 2G, H). We conclude that YAP/TAZ-TEAD regulate the balance between proliferation and terminal differentiation, thereby promoting a highly proliferative and undifferentiated cell state that is relevant for disease progression. This important function is conserved between normal epidermal keratinocytes [32, 36] and neoplastic keratinocytes of both BCC [44] and cSCC (this study).

**Figure 2.**
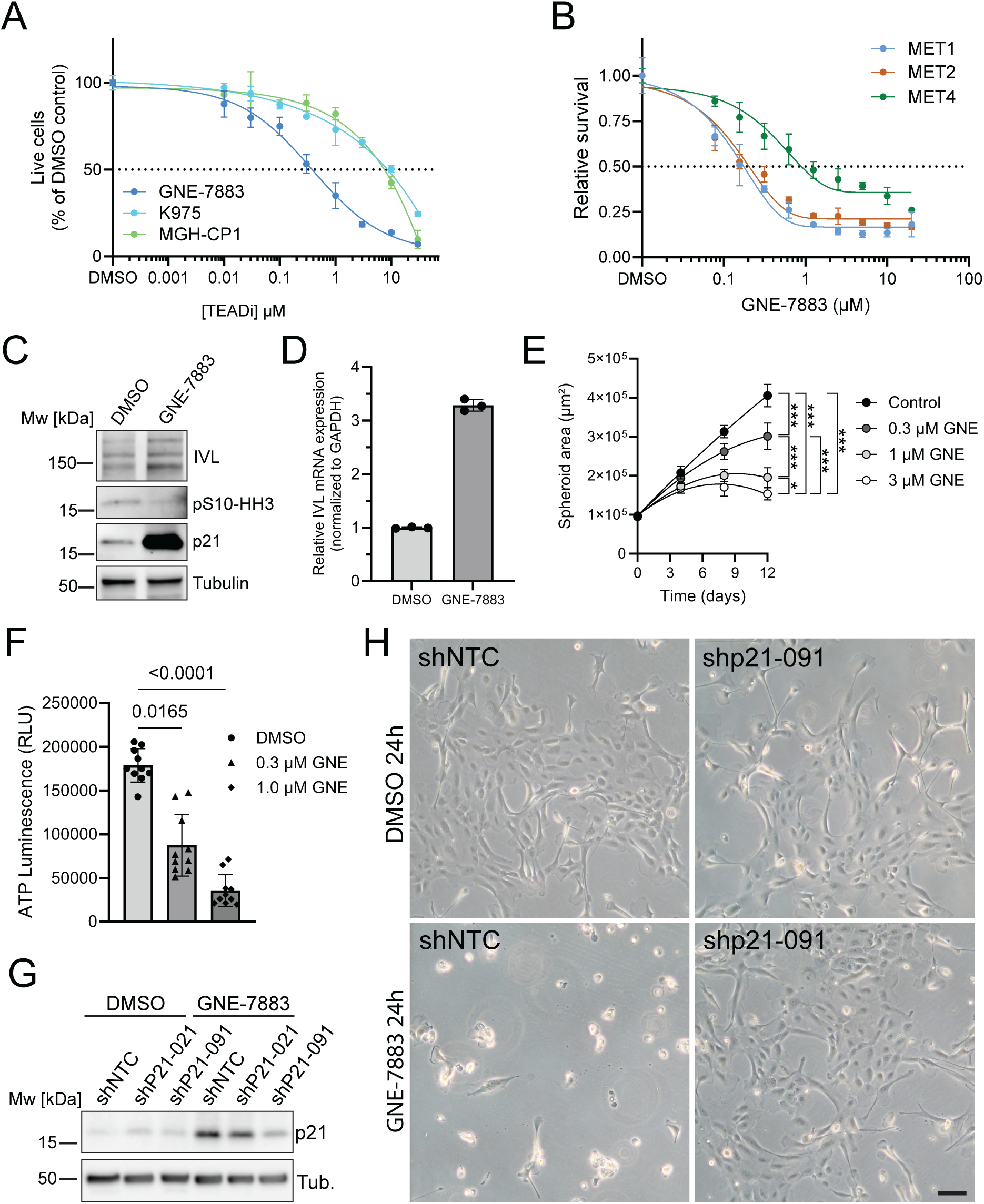
Pharmacological inhibition of YAP/TAZ-TEAD complex formation phenocopies YAP depletion. (A) Representative proliferation dose–response curves (means±SD) of SCC13 cells treated with different TEAD inhibitors. (B) Representative viability dose–response curves (means±SD) of different cell lines from a human cSCC progression series [60] treated with different GNE-7883 concentrations. (C) Immunoblot analysis of SCC13 cells treated with vehicle control (DMSO) or GNE-7883 (2 µM) for 48 h, using the indicated antibodies. Tubulin was used as loading control. (D) qRT-PCR analysis of IVL mRNA expression in SCC13 cells treated with GNE-7883 (2 µM). Data shown are from n=1 experiment performed with three technical replicates. Individual data points: mean fold change in mRNA abundance (normalized to GAPDH) compared to DMSO control. Bars represent the means from all replicates; error bars represent SD. (E) SCC13 3D spheroid growth in the presence of vehicle (DMSO) or different concentrations of GNE-7883 (GNE). p-values are shown for multiple comparisons at 12 days timepoint, Ordinary two-way ANOVA and Dunnett’s multiple comparison test. *p<0.033, **p<0.002, ***p<0.001. Sources of variation: GNE treatment, p<0.001; time, p<0.001; GNE treatment x time, p<0.001. (F) Cell viability measured at the endpoint of the experiment shown in (E). Each datapoint represents one spheroid. Bars represent the means from all spheroids; error bars represent SD. Exact p-values are shown, Kruskal-Wallis test with Dunn’s multiple comparison test. (G) Immunoblot analysis of SCC13 cells expressing shNTC or different shP21 treated with vehicle control (DMSO) or GNE-7883 (2 µM), using the indicated antibodies. Tubulin was used as loading control. (H) Phase contrast images show that p21 depletion rescues cell viability in GNE-7883 (2 µM)-treated SCC13 cells. Bar, 10 µm.

### YAP/TAZ-TEAD-mediated oncogenic signalling is major contributor to replication stress in squamous cell carcinoma cells

Reduced activation of the replication stress checkpoint (phosphorylation of CHK1 on Ser317; [45, 46]) was evident in SCC13 cells upon shYAP but not shTAZ knockdown (Supplementary Fig. 3A), while depletion of either YAP or TAZ decreased the levels of γH2AX. In the context of replication stress, γH2AX marks accumulation of DNA double strand breaks (DSBs) at collapsed replication forks [6, 23, 46] (Supplementary Fig. 3A). Furthermore, using mRNA expression data from the DepMap project [47], we discovered a strong correlation between the expression of a replication stress signature [48] and an SCC-specific YAP/TAZ activity signature comprising 10 highly-correlated YAP/TAZ target genes from our RNAseq dataset in HNSCC (Supplementary Table 1, Supplementary Figs. 4A–D). To provide additional evidence for a causal link between oncogenic YAP signalling and replication stress, we overexpressed wildtype YAP and hyperactive YAP5SA and YAPS127A mutants [33] in SCC cells. As shown in Supplementary Fig. 3B, all three YAP transgenes increased YAP’s transcriptional output, as evident from strongly elevated expression of the canonical YAP/TAZ-TEAD target gene CCN1, and notably reduced cell proliferation when compared to empty vector control (Supplementary Fig. 3C). Both wildtype YAP and YAPS127A enhanced replication stress, as evident from increased γH2AX expression and increased CHK1 Ser345 phosphorylation [45, 46], while YAP5SA overexpression caused increased p21 expression, probably indicating oncogene-induced senescence (Supplementary Fig. 3B).

To demonstrate that transcription can be a general contributor to replication stress in SCC cells, we transiently inhibited RNA synthesis in SCC13 cells using triptolide before assessing γH2AX levels [49]. Triptolide treatment reduced RNA synthesis (Supplementary Fig. 3D) and γH2AX expression to a similar extend as TEAD inhibition by GNE-7883 (Supplementary Figs. 3E, F). Together, these findings showed that YAP/TAZ-TEAD-driven oncogenic signalling is a major contributor to the endogenous replication stress of SCC cells.

### A fraction of chromatin-associated YAP interacts with RIF1

To understand how YAP executes its transcriptional roles, we performed rapid immunoprecipitation mass spectrometry of endogenous protein (RIME) analysis [50] in SCC13 cells and normal human epidermal keratinocytes (NHK) (Fig. 3), using a knockdown-validated YAP-specific antibody that does not recognise TAZ (Supplementary Figs. 5A, B, G). YAP immunoprecipitation (IP) efficiency was consistently high across all YAP-RIME replicates, with high YAP peptide coverage (63-65%) detected by mass spectrometry analysis (Supplementary Fig. 5C and Supplementary Table 2). Upon stringent filtering of the mass spectrometry data (see Supplementary Materials and Methods) YAP-RIME enabled identification of 47 and 14 high confidence YAP-interacting proteins in SCC13 cells (Fig. 3A, Supplementary Table 2) and NHK cells (Supplementary Fig. 5D and Supplementary Table 2), respectively. Significantly enriched gene ontology terms were all related to processes occurring in the nucleus in both interactomes (Fig. 3B, Supplementary Fig. 5E). Among the SCC13 YAP-interacting proteins were several known regulators of YAP, including Neurofibromatosis type (NF2) [51], Angiomotin-like 1 (AMOTL1) [52], PPP1CB [53], and Prolyl 4-Hydroxylase subunit alpha 2 (P4HA2) [54] (Fig. 3A). Our YAP-RIME also identified several subunits of the BAF-type SWI/SNF chromatin remodelling complex (ARID1A, SMARCA4, SMARCC1, SMARCE1, SMARCD2) [55], and key components of the NuRD chromatin remodelling complex, Chromodomain-helicase-DNA-binding protein 4 (CHD4) and Metastasis Associated 1 Family Member 2 (MTA2). While key components of the general transcriptional machinery (POLR2A, POLR2B, GTF21, TOP1) were present in the SCC13 YAP interactome, TEAD TFs were notably absent (Fig. 3A). A closer inspection of our mass spectrometry data revealed that TEADs, as well as other known YAP-interacting transcription factors, such as JunB and TP63 [56, 57], had been identified but did not pass our stringent bioinformatics filtering criteria (Supplementary Table 2). The overall poor identification of TFs by our YAP-RIME was likely due to their insufficient crosslinking to the DNA by formaldehyde alone, as previously observed for the highly dynamic transcription factor NFκB [58].

**Figure 3.**
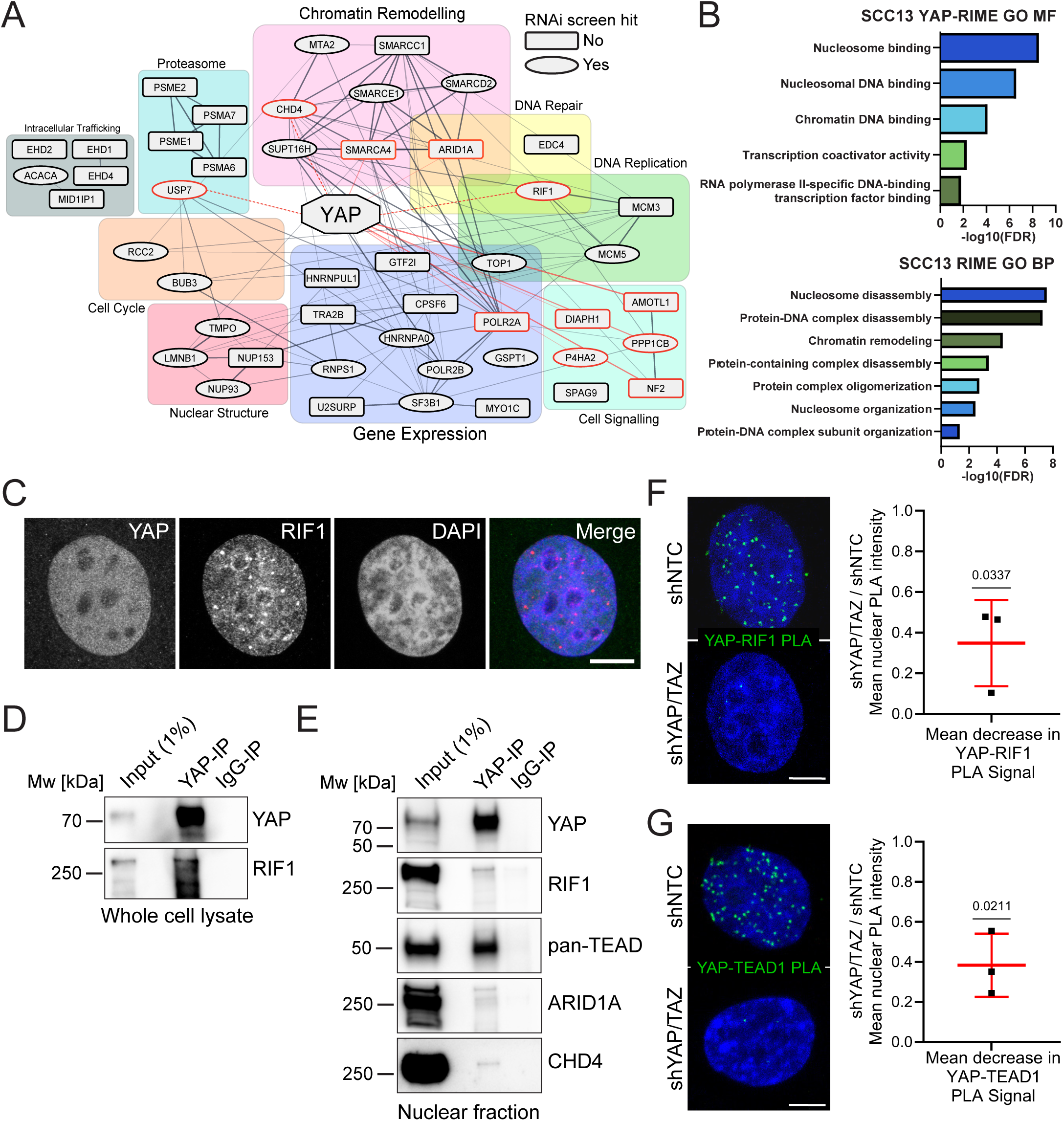
RIME identifies RIF1 as an interactor of chromatin-associated YAP in squamous cell carcinoma cells. (A) Interaction network of YAP protein interactors identified by RIME in SCC13 cells. Oval nodes: proteins that were previously identified as candidate regulators of SCC13 cell proliferation and/or survival in a genome wide RNA-interference (RNAi) screen (hit: ‘Yes’)[32]; rectangular shaped nodes: proteins that were not identified in the RNAi screen (hit: ‘No’). Red connecting lines (edges) show previously identified YAP interactions according to the STRING Homo sapiens database, and nodes with red borders indicate known YAP interactors (manually curated from STRING and literature search); dashed red lines show manually curated known YAP interactions that were not yet present in the STRING database. Proteins were manually grouped based on function (gene names displayed, generated in Cytoscape 3.9.1 using STRING app 1.7.1). (B) Gene ontology terms (‘molecular functions’ and ‘biological process’) enriched within the high-confidence YAP-interacting proteins identified by RIME. (C) Confocal fluorescence image (z-projection) of YAP (green) and RIF1 (red) nuclear localisation in SCC13 cells. DNA was counterstained with Hoechst dye (blue). Scale bar, 10 µm. (D, E) Immunoblot analysis of YAP and IgG-control immunoprecipitates and respective inputs from SCC13 (D) whole cell extracts and (E) and nuclei-enriched fractions using the indicated antibodies. (F, G) Representative confocal microscopy images (z-projections) and image quantifications of (F) YAP-RIF1 and (G) YAP-TEAD1 in-situ PLA in control (siNTC&shNTC) and YAP/TAZ knockdown (siYAP/TAZ&shYAP) cells (PLA signal, green; DNA (DAPI) blue; n >250 cells (YAP-RIF1), n >500 cells (YAP-TEAD1) per condition in n=3 independent experiments). Scale bars, 5 µm. Datapoints show fold decrease in mean nuclear PLA signal intensity in siYAP/TAZ&shYAP cells compared to siNTC&shNTC cells. Lines show means from all three experiments, error bars show SD. Exact p-values are shown, one sample two-tailed t test.

To identify physiologically relevant interactions, the YAP-interacting proteins were compared to candidate positive and negative regulators of cell proliferation and survival identified in a prior genome-wide RNA screens in SCC13 [32]. This revealed that 22 out of the 47 high confidence YAP-interacting proteins were likely relevant for cell proliferation and/or survival (Fig. 3A, Supplementary Table 2).

Our attention was drawn to RIF1, given its recent identification as a non-transcriptional YAP interaction partner in a developmental context in Xenopus laevis [59]. We detected RIF1 as a YAP-chromatin-interacting protein in both the SCC13 and NHK YAP-RIME experiments (Fig. 3A, Supplementary Fig. 5D) with a much stronger enrichment in SCC13 YAP (Supplementary Fig. 5F and Supplementary Table 2). Of note, PPP1CB, one of the three catalytic subunits of PP1―a key effector of RIF1 [13, 16, 17, 19], was also present in the SCC13 YAP-RIME (Fig. 3A, Supplementary Table 2). Like YAP, RIF1 featured among the positive regulators of cell proliferation and survival of SCC13 cells identified in the aforementioned genome-wide RNA interference screen [32] (Fig. 3A, Supplementary Table 2). As expected, YAP and RIF1 were both localised in the nucleus of SCC13 cells (Fig. 3C) and present in chromatin fractions (Supplementary Fig. 5G), confirming their chromatin-association. To biochemically validate the YAP-RIF1 interaction, co-IP experiments were performed under native non-cross-linking conditions using a different YAP-specific antibody (Supplementary Fig. 6A). Indeed, RIF1 could be immunoprecipitated together with YAP from both, whole-cell detergent extracts and nuclei-enriched fractions (Figs. 3D, E). Further validating our RIME results, CHD4 and ARID1A also co-immunoprecipitated with YAP under native conditions. Under these conditions, TEAD transcription factors could also be immunoprecipitated together with YAP (Fig. 3E). In-situ proximity ligation assays (PLA) confirmed very close YAP-RIF1 proximity in the nucleus of SCC13 cells (Fig. 3F). To achieve a sufficiently strong knockdown of both YAP and TAZ to control for PLA signal specificity, we combined our doxycycline-inducible YAP-targeting shRNAs with an siRNA pool targeting YAP and TAZ (Fig. 3F, Supplementary Fig. 6B). The YAP-RIF1 PLA signal was significantly decreased upon YAP/TAZ knockdown. Consistent with our co-IP results, YAP was also found in very close proximity to TEAD1 in the nucleus of SCC13 cells by PLA (Fig. 3G). The YAP-RIF1 PLA signal was completely absent when the two primary antibodies were omitted (Supplementary Figs. 6C, D) and also largely absent from mitotic chromosomes (Supplementary Fig. 6E). To test if the YAP-RIF1 interaction also occurred in other types of SCC, we used the lung SCC (LUSC) cell line HARA. Co-IP was performed with nuclei-enriched HARA cell fractions treated with Benzonase nuclease to enzymatically digest DNA and RNA (Supplementary Figs. 6F, G), demonstrating that YAP and RIF1 are not co-immunoprecipitated because of a DNA bridge between the two proteins. In summary, our results show that RIF1 is a binding partner of chromatin-associated YAP in both healthy human NHKs and cancerous cSCC and LUSC cells.

### YAP/TAZ-TEAD modulate RIF1 protein levels

Our RNAseq analysis revealed that RIF1 mRNA expression was moderately downregulated in response to siYAP/TAZ (Supplementary Fig. 1A and Supplementary Table 1). A similar, albeit statistically not significant trend could be confirmed in independent siYAP/TAZ samples by RT-qPCR analysis (Fig. 4A). These mild effects of YAP depletion on RIF1 mRNA expression are likely indirect and may be linked to cell cycle de-regulation. Depletion of RIF1 did not impact on expression of most of the DEGs seen upon siYAP/TAZ (Supplementary Figs. 1B, C, and 2B, C), ruling out a direct involvement in YAP/TAZ-TEAD-controlled transcription.

**Figure 4.**
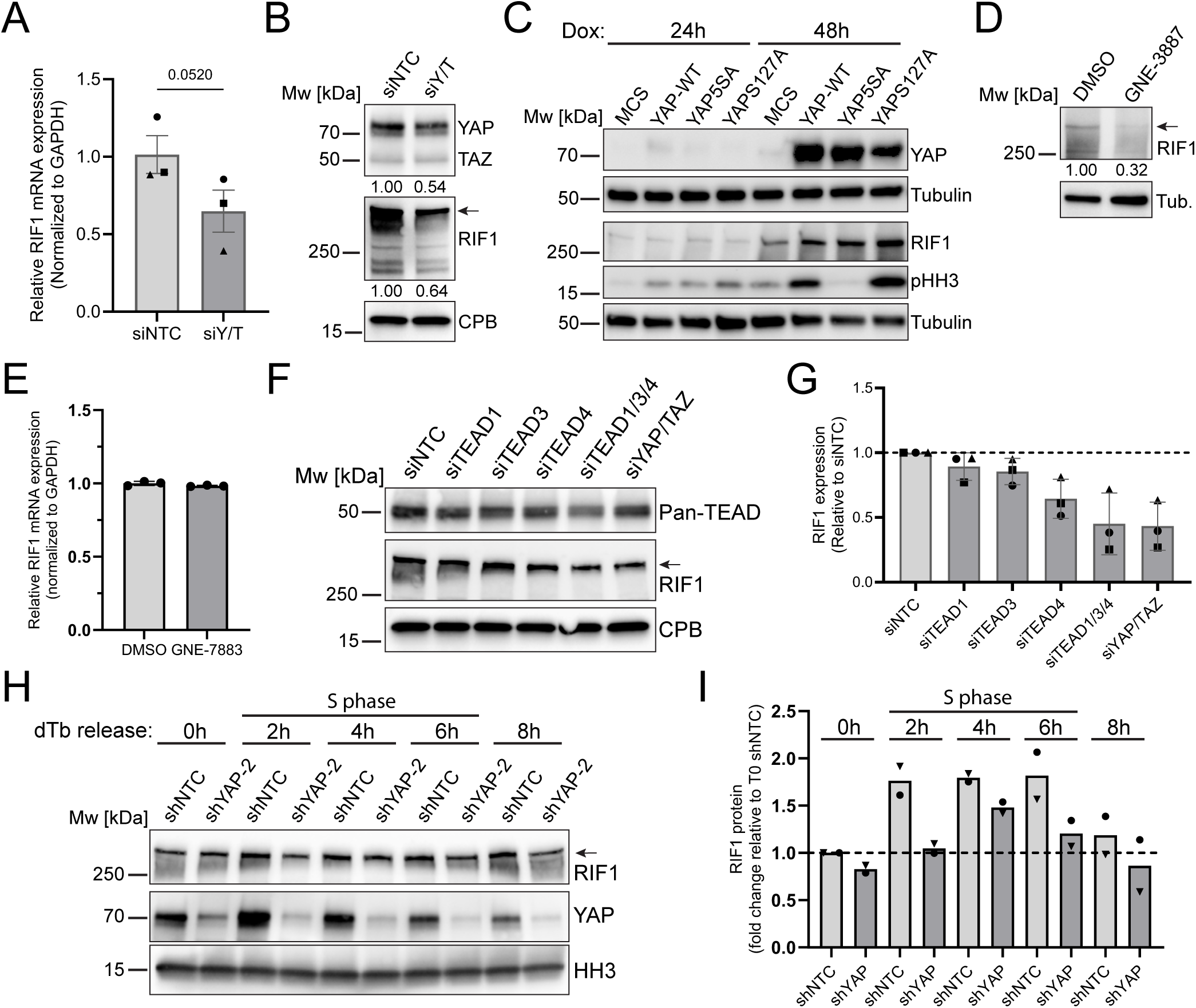
YAP/TAZ-TEAD modulate RIF1 protein stability. (A) qRT-PCR analysis of RIF1 mRNA expression in SCC13 cells transfected with an siRNA SMARTpool targeting YAP/TAZ (siY/T) or a non-targeting control SMARTpool (siNTC) for 48 h. Data shown are from n=3 independent experiments performed with three technical replicates. Individual data points: mean fold change in mRNA abundance (normalized to GAPDH) compared to siNTC in each experiment. Bars represent the means from all experiments, error bars represent SD. Exact p-values are shown, paired t test. (B–D) Immunoblot analysis of SCC13 cells (B) transfected with siNTC or siY/T for 48 h, (C) expressing empty vector control (MCS) or different YAP transgenes upon doxycycline (dox) induction, or (D) treated with GNE-7883 (2 µM) for 24 h, using the indicated antibodies. Tubulin or cyclophilin B (CBP) were used as loading controls. Arrows, position of full-length RIF1 bands. In (B, D) numbers below lanes represent protein ratios relative to an arbitrary level of 1.0 set for siNTC or DMSO control, respectively. (E) qRT-PCR analysis of RIF1 mRNA expression in SCC13 cells treated with GNE-7883 for 24 h. Data shown are from n=1 experiment performed with three technical replicates. Individual data points: mean fold change in mRNA abundance (normalized to GAPDH) compared to DMSO control. Bars represent the means from all replicates, error bars represent SD. (F) Immunoblot analysis of RIF1 protein levels in SCC13 cells transfected with siRNA SMARTpools targeting TEAD1, TEAD3, TEAD4, or YAP/TAZ (siY/T), or an siNTC SMARTpool for 24 h. Cyclophilin B (CBP) was used as loading control. (G) Quantification of RIF1 protein expression in experiment shown in (F). Individual data points show fold change in RIF1 protein levels (normalised to tubulin) in siTEAD1, siTEAD3, siTEAD4, siTEAD1/3/4, and siY/T compared to siNTC in n=3 independent experiments (different shapes of data points), bars show the mean, error bars show SD. (H) Immunoblot analysis of chromatin fractions from SCC13 cells expressing doxycycline-induced shYAP or shNTC for 48 h, using the indicated antibodies. Cells were synchronised at the G1/S boundary by double thymidine block (dTb) and released into the S phase. Histone H3 (HH3) was used as loading control. Arrow, position of full-length RIF1 band. (I) Quantification of RIF1 protein levels in experiment shown in (H). Individual data points show fold change in RIF1 protein levels (normalised to HH3) compared to shNTC at timepoint 0 h in n=2 independent experiments (different shapes of data points).

The stability of a protein within a complex often depends on other components of the complex, as proper folding of one protein may require interactions with its partners. RIF1 protein expression was lower in siYAP/TAZ compared to siNTC SCC13 cells (Fig. 4B). Conversely, overexpression of wildtype and oncogenic YAP variants increased RIF1 protein levels irrespective of their effects on the cell cycle (pHH3) (Fig. 4C). Interestingly, short-term treatment (24 h) with GNE-7883 to disrupt YAP/TAZ-TEAD binding reduced RIF1 protein expression without affecting RIF1 mRNA levels (Figs. 4D, E). This suggests that YAP might modulate RIF1 protein levels in a complex with TEADs. Indeed, combined siRNA-mediated knockdown of TEAD1, 3, and 4 (siTEAD1/3/4) in SCC13 cells reduced RIF1 protein to levels similar to those observed in siYAP/TAZ cells (Figs. 4F, G). Interestingly, depletion of YAP impacted chromatin-associated RIF1 levels specifically during S phase (Figs. 4H, I). Since the data presented in Figs. 3E–G showed that YAP is in complex with RIF1 and TEADs, these findings thus suggest that YAP-TEAD modulate protein levels of a fraction of chromatin-associated RIF1 in S phase.

### RIF1 is highly expressed in squamous cell carcinoma and upregulated during tumour progression

To understand if RIF1 contributes to squamous carcinogenesis, we assessed RIF1 protein expression in a panel of human cSCC cell lines, which also included an isogenic cSCC progression series (PM1, MET1–4), including the pre-malignant PM1 precursor lesion [60]. Increased RIF1 protein levels were notable in all cSCC cell lines when compared to NHKs (Fig. 5A). Consistent with our findings that YAP-TEAD modulate RIF1 protein levels (Fig. 4), RIF1 protein expression was highest in SCC13 cells featuring high nuclear YAP levels [32], as also indicated by a low ratio of S127-YAP to total YAP (Fig. 5A). All cSCC lines displayed higher levels of γH2AX compared to NHKs, indicating increased endogenous replication stress (Fig. 5A).

**Figure 5.**
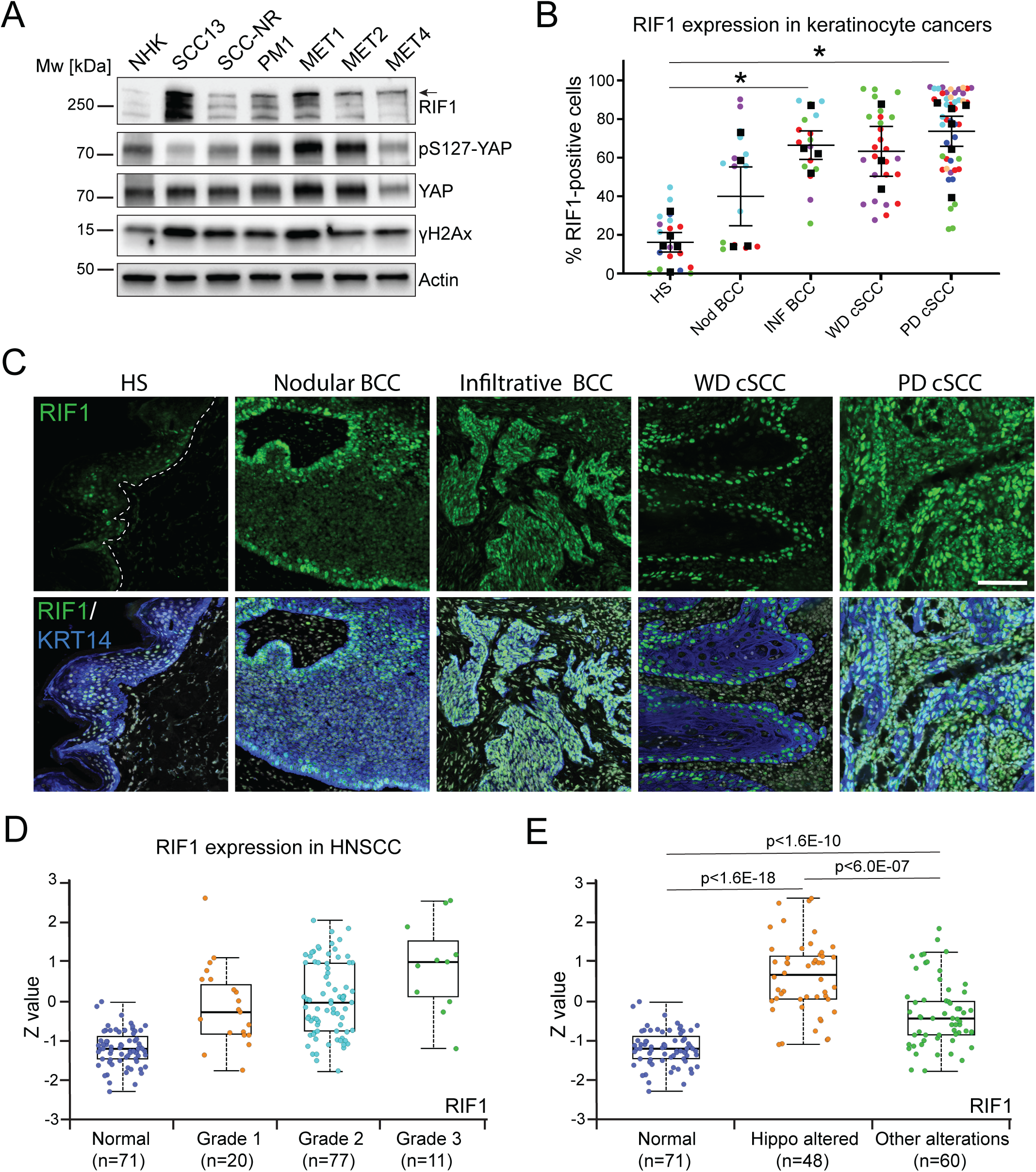
Increased expression of RIF1 in squamous cell carcinoma. (A) Immunoblot analysis of normal human epidermal keratinocytes (NHKs), keratinocytes isolated from a pre-malignant dysplastic skin lesion (PM1), various cell lines isolated from primary cutaneous SCCs (cSCC; SCC13, SCC-NR, MET1), a recurrent cSCC (MET2), and a lymph node metastasis (MET4), using the indicated antibodies. Actin was used as loading control. Arrow, position of full-length RIF1 band. (B, C) Quantification (B) and representative images (C) of RIF1 expression in healthy adult human skin (HS), nodular (Nod) and (Inf) infiltrative basal cell carcinoma (BCC), and well-differentiated (WD) and poorly differentiated (PD) cSCC. Tissue sections were immunolabelled with antibodies against RIF1 (green), and keratin 14 (Krt14, blue) to identify normal and neoplastic epidermal keratinocytes. White dotted line demarcates dermal–epidermal boundary in HS. Scale bar, 50 μm. Superplots in show percentage of RIF1-positive keratinocytes in different areas of the same tumour (coloured symbols), mean percentage of RIF1-positive keratinocytes in 3–5 different patient samples (black datapoints), and mean percentage of RIF1-positive keratinocytes in each sample group (lines). Error bars show SD from each sample group. *p<0.05, Brown-Forsythe and Welsh ANOVA test with Dunett’s multiple comparison test. (D, E) RIF1 protein expression in HNSCC samples from the Clinical Proteomic Tumour Analysis Consortium (CPTAC), stratified according to (D) tumour stage, or (E) presence of alterations in the Hippo pathway or other signalling pathways. Jitter plots (overlayed onto box plots) show Z-values (standard deviations from the median across samples for the given sample type) from individual samples. Box plots indicate the median (middle line), 25^th^ percentile (bottom line), 75^th^ percentile (top line), and minimum and maximum (whiskers). Log2 spectral count ratio values from CPTAC were first normalized within each sample profile, then normalized across samples. Graphs were downloaded from the UALCAN database (https://ualcan.path.uab.edu/). (D) p<0.05 for all pairwise comparisons. In (E) exact p-values are shown.

To investigate if high RIF1 expression was a general hallmark of cancers originating from epidermal keratinocytes, we quantified the levels of RIF1 protein in a tissue array comprising cSCCs of different differentiation grades, as well as different types of BCCs (Figs. 5B, C). We detected increased numbers of RIF1-expressing cells within the keratin-positive tumour cell masses of both BCC and cSCC compared to normal skin epidermis, especially within the more aggressive tumour types of infiltrative BCC and poorly differentiated cSCC (Figs. 5B, C). These data are corroborated by immunohistochemistry data from the Human Protein Atlas, depicting increased numbers of RIF1-expressing tumour cells in cSCC, HNSCC and cervical SCC (Supplementary Figs. 7A–C). To obtain a pan-cancer overview of RIF1 expression, we used the UALCAN portal [61] to access protein expression data from the Clinical Proteomic Tumour Analysis Consortium (CPTAC) [62]. In HNSCC, RIF1 protein expression increased with tumour grade (Fig. 5D) and RIF1 expression was higher across several cancer types compared to normal tissue (Supplementary Fig. 7D). The UALCAN portal also allows assessment of the impact of somatic alterations (small mutations or copy number alterations) in signalling pathways on the cancer proteome [61, 62]. We found that in both HNSCC (Fig. 5E) and LUSC (Supplementary Fig. 7E) increased RIF1 protein expression predominantly correlated with somatic Hippo pathway alterations, in contrast to alterations in other signalling pathways, implying altered Hippo pathway activity in the regulation of RIF1 expression. In contrast, in other cancer types, alterations in pathways other than Hippo signalling had a strong impact on increased tumour expression of RIF1 (Supplementary Figs. 7F, G). Collectively, our multi-omics analysis confirm a close relationship between Hippo signalling and RIF1 protein expression and point towards a role of RIF1 in squamous cell carcinogenesis.

### Depletion of RIF1 increases replication fork speed in squamous cell carcinoma cells

Consistent with a role of RIF1 in controlling S phase progression of SCC cells, we observed that on release from a double thymidine block, shRIF1 SCC13 cells progressed more rapidly through S phase (Fig. 6A). Phosphorylation levels of MCM2 and MCM4 proteins, targets of the Dbf4-dependent kinase (DDK; composed of CDC7 kinase and its activator, DBF4) [16, 18], were increased in RIF1-depleted SCC13 cells in a RIF1 dosage-dependent manner (Figs. 6B, C), particularly in early and mid S phase (2- and 4-h time points post-dTb, Fig. 6C), indicating alterations in the dynamics of origin firing [16–19]. Accordingly, chromatin loading of PCNA, which is required for DNA replication after the action of CDC7, was also increased during early and mid S phase of shRIF1 SCC13 cells (Fig. 6C). Therefore, the faster progression of SCC13 cells through S phase after RIF1 depletion likely reflects an increase in DNA replication initiation events in early and mid S phase [16, 18]. We also examined DNA replication dynamics in shRIF1 cells using a novel long-read sequencing method, DNAscent [63]. We discovered that RIF knockdown increased fork speed along with led to longer replication tracks (Figs. 6D, E, and Supplementary Figs. 8A, B).

**Figure 6.**
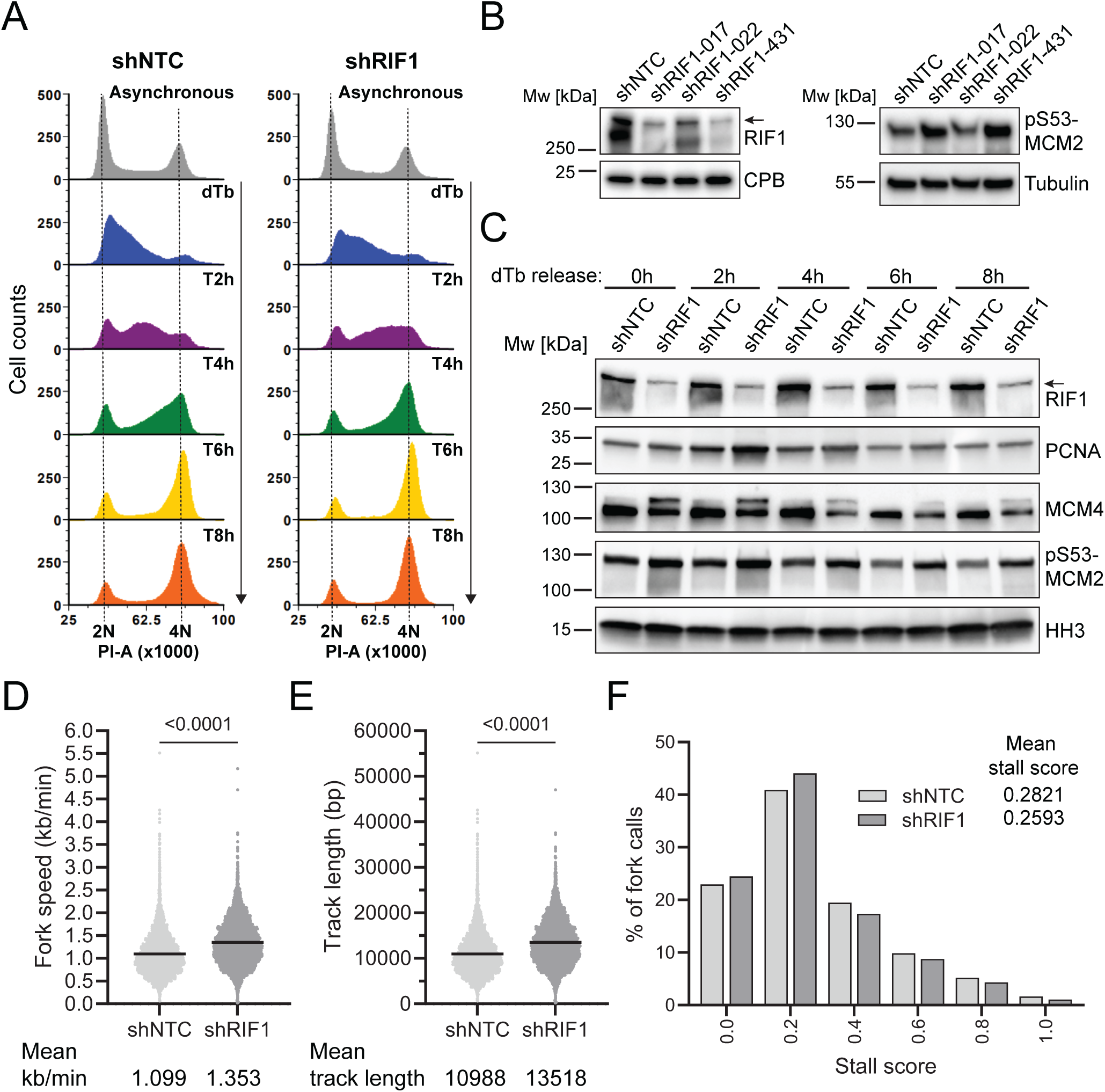
Depletion of RIF1 in squamous cell carcinoma cells affects DNA replication dynamics. (A) Representative cell cycle analysis by PI labelling of asynchronous populations of shNTC and shRIF1 SCC13 cells (grey profiles) and of cells synchronised at the G1/S boundary by double thymidine block (dTb) and released into S phase (T=hours post thymidine release, coloured profiles). (B, C) Immunoblot analysis of (B) cell extracts and (C) chromatin fractions from shNTC and shRIF1 SCC13 cells, using the indicated antibodies. In (C) cells were synchronised at the G1/S boundary by double thymidine block (dTB) and released into the S phase. Note that DDK-induced phosphorylation of MCM4 results in a band shift. Cyclophilin B (CPB), tubulin, and histone H3 (HH3) were used as loading controls. shRIF1-017 was used in experiments shown in (A, C). Arrows in (B, C), position of full-length RIF1 band. (D, E) Replication fork speed (D) and replication track length (E) measured by DNAscent. Mean fork speed (kb/min) and track length (bp) are indicated. Black bars show medians. Total number of forks scored in n=2 independent experiments: shNTC=5632; shRIF1=6216. (F) Combined distribution of stall scores from n=2 independent experiments. Mean stall cores are indicated.

### RIF1 protects against endogenous replication stress in squamous cell carcinoma cells

Under replication stress, RIF1 protects stalled replication forks [16, 20–22], but also facilitates homology-mediated DDR at broken replication forks [23]. The elevated expression of RIF1 in SCC cells could therefore indicate a role for RIF1 in enabling the cells to deal with high levels of endogenous replication stress that are, at least in part, controlled by YAP-TEAD-driven oncogenic mechanisms (Fig. 2). Indeed, we found a strong correlation between RIF1 expression and a replication stress signature [48] in HNSCC cell lines (Supplementary Figs. 4A, E). In contrast to a slowing down or stalling of replication fork progression, which is frequently observed when cells experience replication stress [3–5], our DNAscent analysis revealed increased replication fork speed upon RIF knockdown (Figs. 6D, E). Moreover, replication fork stalling or pausing, as defined by a shift of DNAscent ‘stall score’ distribution closer to 1.0, was not detected (Fig. 6F). Previous studies have found that increased replication speed can lead to DNA damage, whereby faster replication forks have insufficient time to recognize damaged DNA in need of repair [64, 65]. The only moderate activation of CHK1 (phosphorylation on Ser317 and S345; [45, 46]) and very mild increase of ATR-mediated S33-phosphorylation of RPA32 [46] in shRIF1 SCC13 cells, in contrast to the strongly elevated levels of γH2AX and hyperphosphorylated pS4/8-RPA32 (indicating phosphorylation by ATM and DNA-PK in response to generation of DNA DSBs [46, 66, 67] (Figs. 7A, B) are corroborating such a scenario. Indeed, replication fork collapse and concomitant activation of ATM/DNA-PK signalling were previously observed in response to ATR inhibition in cells experiencing hydroxyurea (HU)-induced replication stress [67]. It is worth noting that DNAScent only detects short (<20 min) replication fork stalls/pauses and currently cannot measure longer fork stalls and fork breakage. Given that RIF1 facilitates repair of broken replication forks [23] it is likely that loss of this additional function also contributes to the increased accumulation of DNA damage in shRIF1 cells. Because the replication checkpoint was not robustly activated in shRIF1 SCC13 cells (Figs. 7A, B), they were able to progress through the cell cycle despite persistent DNA damage (Figs. 7C, D). A small increase in the G2/M fraction in shRIF1 SCC13 cells (Figs. 7C, D) was noticeable as well as a slightly elevated pHH3 expression in siRIF1 cells (Fig. 7E). The accumulation of DNA damage in cultures of shRIF1 SCC13 cells became more accentuated after a few passages, when cells had undergone multiple rounds of cell division (Fig. 7F), ultimately resulting in reduced clonal growth (Figs. 7G, H). To confirm that the replication stress-protecting effect of RIF1 is conserved in other types of SCC, we depleted RIF1 in the HNSCC cell line SAS, which displays a strong correlation between RIF1 expression and replication stress (Supplementary Fig. 4E) and carries a YAP-MAML2 gene fusion resulting in a hyperactive YAP variant [68]. Indeed, we observed a dosage-dependent effect of shRIF1 on cell viability (Supplementary Figs. 9A–C) and DNA damage (Supplementary Figs. 9D, E). Decreased cell viability (Supplementary Fig. 9F) and increased γH2AX levels (Supplementary Fig. 9G) were also detected in the LUSC cell line HARA upon shRIF1. Together, our results point to a scenario where SCC cells depend on elevated RIF1 expression to dampen their high levels of endogenous replication stress.

**Figure 7.**
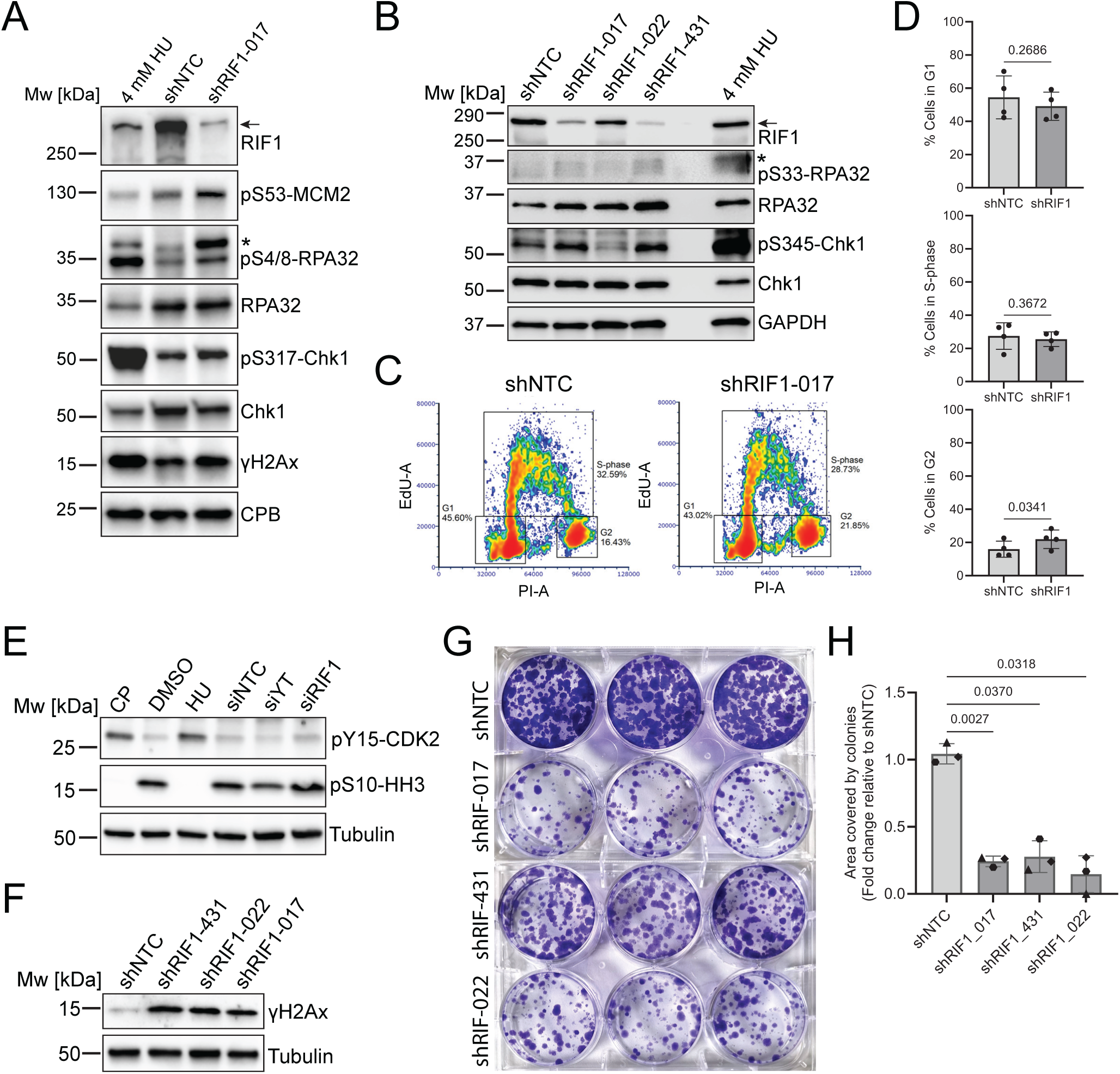
RIF1 protects against endogenous replication stress. (A, B) Immunoblot analysis of cell extracts from SCC13 cells expressing shNTC or different shRIF1, using the indicated antibodies. Cyclophilin B (CPB), tubulin, or GAPDH were used as loading controls. Cells treated for 24 h with 4 mM HU served as positive controls for severe replication stress. Positions of strongly hyperphosphorylated RPA32 bands are indicated with asterisks. Arrows, positions of full-length RIF1 bands. (C) Representative cell cycle profile (PI labelling following a 30-min EdU pulse) of unsynchronised shNTC and shRIF1 SCC13 cells. Gates used for quantification of percentage of cells in the different cell cycle phases are shown. (D) Quantification of percentage of cells in different cell cycle phases. Bars show means from n=4 independent experiments, individual data points show cell percentages in each experiment, error bars show SD. Exact p-values are shown, paired t test. (E, F) Immunoblot analysis of cell extracts from SCC13 cells transfected with (E) siNTC, siYAP/TAZ (siY/T), or siRIF1 for 48 h, or treated with cisplatin (CP, 4 µM) or hydroxyurea (HU, 4 mM) for 24 h to induce S phase arrest, or (F) expressing shNTC or different shRIF1, using the indicated antibodies. Tubulin was used as loading control. (G) Representative images of colony formation efficiency of shNTC and shRIF1 SCC13 cells. Cells in triplicate wells were stained with crystal violet. (H) Percentage of well area occupied by colonies. Bars show the means from n=3 independent experiments (performed with 3 technical replicates), individual data points (different shapes indicate different experiments) show the means from each experiment, error bars show SD. Exact p-values are shown, RM one way ANOVA with Geisser-Greenhouse correction and Dunnett’s multiple comparison test.

### YAP is involved in DNA damage repair at collapsed replication forks

Our discovery of RIF1 as a YAP interaction partner in human SCC cells raised the exciting possibility that in this cancer context, YAP could play dual roles in both transcription and DNA replication under replication stress. To test, if YAP, like RIF1, might function at collapsed replication forks [23], we treated SCC13 cells with 4 mM HU [23, 69] for 24 h. This triggered strong activation of the replication stress checkpoint (phosphorylation of CHK1 on Ser317; [45, 46]) and accumulation of DNA DSBs, marked by strongly elevated γH2AX expression (Fig. 8A). Persisting γH2AX expression following HU washout indicated accumulation of DNA damage at stalled and subsequently inactivated replication forks [69] (Fig. 8A). This was further confirmed by the presence of RAD51 foci [23] (Fig. 8B) and appearance of pan-nuclear γH2AX staining, indicating severe DNA replication stress [70], upon 24 h HU treatment (Supplementary Fig. 10A). Indeed, some cell death (appearance of sub-G1 fraction in EdU/PI cell cycle profiles, Supplementary Fig. 10A) was noticeable from 15 h HU treatment onwards. Interestingly, YAP protein levels were increased following washout of HU after 24 h of treatment, suggesting a potential role of YAP in DDR upon replication fork collapse (Fig. 8C). Indeed, while the YAP-RIF1 PLA signal remained unchanged upon short-term (3 h) HU treatment, it increased significantly after long-term (24 h) HU treatment (Fig. 8D). Moreover, upon long-term HU treatment, more YAP molecules were found in close vicinity of γH2AX by in-situ PLA (Fig. 8E, Supplementary Fig. 10C), suggesting that YAP associates with RIF1 preferentially near DNA DSBs at broken replication forks. To test if YAP is directly involved in DDR at broken replication forks, we optimized conditions for RNA interference experiments to only minimally perturb the cell cycle in response to siYAP/TAZ (Fig. 8F, Supplementary Fig. 10D). For these experiments, HU concentration was reduced to 2 mM to avoid potential confounding effects of HU-induced cell death. As shown in Fig. 8F, by 48 h post-transfection with a low concentration of siYAP/TAZ (1 nM), SCC13 cells displayed reduced levels of YAP and only modest upregulation of p21 and downregulation of pHH3, while γH2AX levels (as a read-out of YAP-TEAD-driven replication stress) were strongly reduced. Treatment with 2 mM HU during the last 24 h post-siYAP/TAZ transfection led to prominent S phase arrest, as evident from strongly reduced pHH3 levels and replication stress checkpoint activation (phosphorylation of CHK1 on Ser345) (Fig. 8F). Under these conditions, YAP depletion did not cause a decrease in γH2AX levels but instead led to a significant increase in comparison to untreated siYAP/TAZ cells (Fig. 8F). Since transcription is globally dampened under high concentrations of HU [71] and YAP was depleted specifically during the HU treatment period, these findings suggest a novel non-transcriptional function of YAP where it is involved in DDR at broken replication forks.

**Figure 8.**
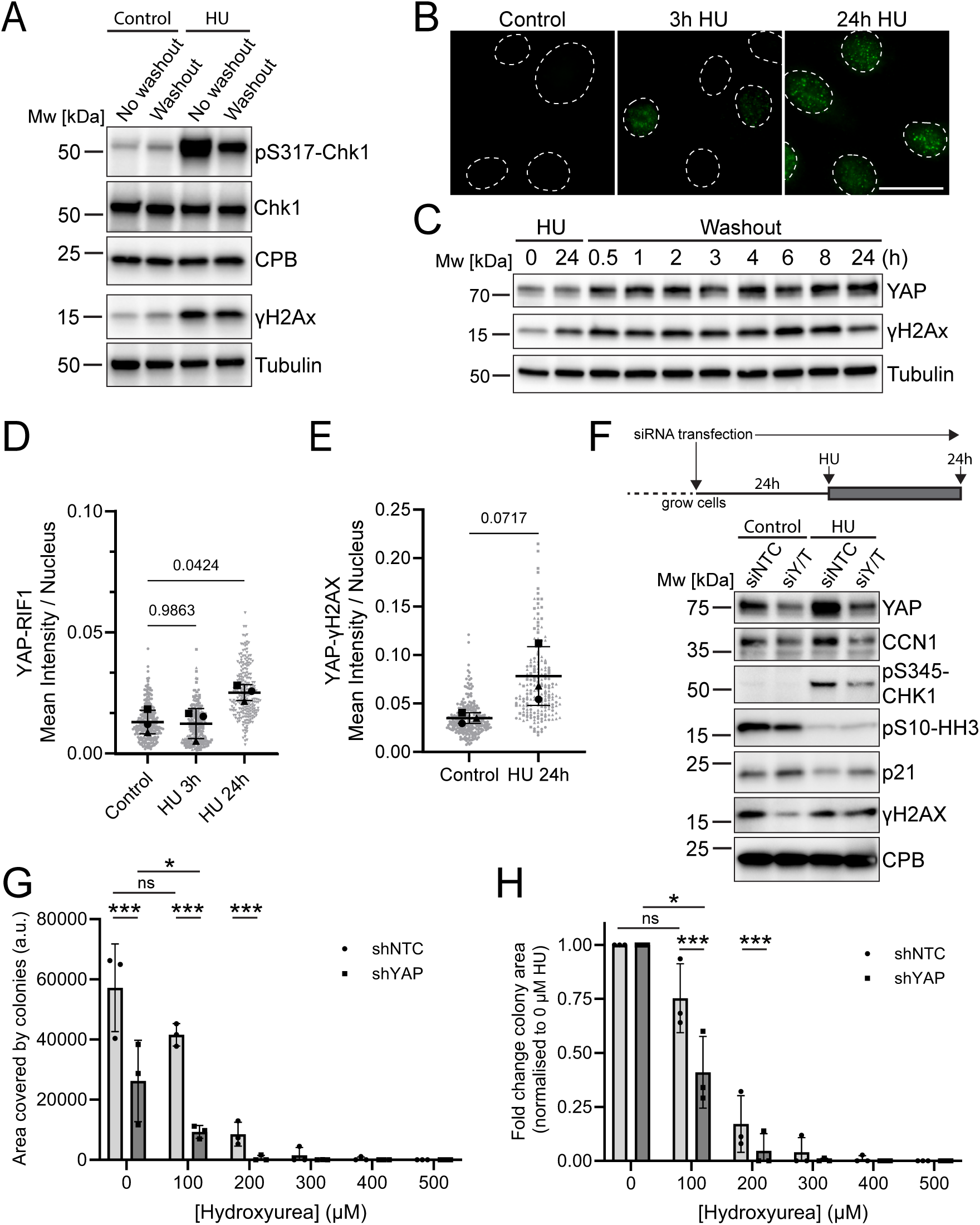
YAP is involved in DNA damage repair at collapsed replication forks. (A) Immunoblot analysis of SCC13 control cells and cells exposed to 4 mM hydroxyurea (HU) for 24 h, using the indicated antibodies. Washout, medium removal and replacement with fresh medium. Tubulin or cyclophilin B (CBP) were used as loading controls. (B) Representative microscopy images of RAD51 (green) foci in SCC13 cells treated with 4 mM HU for the indicated times. Dashed outlines indicate nuclear areas from DNA counterstain with DAPI. Scale bar, 25 µm. (C) Immunoblot analysis of SCC13 cells exposed to 4 mM HU for 24 h, followed by HU washout, using the indicated antibodies. Tubulin was used as loading control. (D, E) Image quantifications of (D) YAP-RIF1 and (E) YAP-γH2AX in-situ PLA in SCC13 cells treated with 4 mM HU for the indicated times. >200 cells were measured per experimental condition in n=3 independent experiments. Superplots show nuclear PLA signal intensity per cell (grey datapoints), mean nuclear PLA signal intensity of the respective cell populations in each of n=3 experiments (black datapoints, different shapes indicate different experiments), and mean nuclear PLA signal intensity from all three experiments (lines). Error bars show SD from all three experiments. Exact p-values are shown. (D) Brown-Forsythe and Welsh ANOVA test with Dunett’s multiple comparison test, (E) unpaired t test. (F) Immunoblot analysis of SCC13 cells treated with 2 mM HU for 24 h or left untreated (control), 24 h after transfection with siNTC or siYAP/TAZ (siY/T), using the indicated antibodies. Tubulin was used as loading control. (G, H) Synergism of YAP depletion (shYAP) and HU treatment on clonal colony growth was quantified by measuring the percentage of well area occupied by colonies after 10 days in culture. In (H) values have been normalized to controls (no HU treatment) for shYAP and shNTC. Bars show the means from n=3 independent experiments (performed with 3 technical replicates), individual data points show the means from each experiment, error bars show SD. p-values are shown for multiple comparisons, RM two-way ANOVA (matched values are spread across a row) and Šidak’s multiple comparison test. *p<0.033, **p<0.002, ***p<0.001. Sources of variation in (G, H): HU concentration, p<0.001; shRNA, p<0.001; HU concentration x shRNA, p<0.001.

Given the dual function of YAP in promoting TEAD-mediated transcription and facilitating DDR under severe replication stress, we examined if pharmacologically induced replication stress could act synergistically with YAP depletion to impair SCC proliferation. Of note, HU is sometimes used in chemoradiotherapy for HNSCC patients [72]. Indeed, we observed a synergistic effect of shYAP and low doses of HU on SCC13 clonal colony growth (Figs. 8G, H). This finding highlights the possibility of using pharmacological replication stress inducers in combination with YAP-TEAD inhibitors [73] to treat SCC.

## Discussion

Our findings provide novel insight into the oncogenic mechanisms of hyperactive YAP/TAZ in SCC. While YAP/TAZ promote proliferation by regulating transcriptional programmes that drive cell cycle progression, they also suppress commitment to terminal differentiation. Thus, the functions of YAP/TAZ are highly conserved within the epidermal lineage, as both normal epidermal stem cells [32, 33, 36] and neoplastic keratinocytes in BCC [44] and cSCC (this study) show strong dependency on YAP/TAZ-TEAD signalling for cell proliferation. This suggests that cancers originating from epidermal stem/progenitor cells ‘hijack’ the tissue regenerating functions of YAP/TAZ to fuel their high rates of proliferation while remaining in an undifferentiated cell state. The increased expression of genes that are positively regulated by YAP/TAZ in tumour tissue of human cSCCs that underwent metastatic progression (Fig. 1) provides strong support for this idea. Our RIME analysis revealed that the transcriptional co-regulator functions of YAP/TAZ are likely mediated by the BAF-type SWI/SNF and the NuRD chromatin remodelling complexes. Indeed, in HNSCC cells, BAF-type SWI/SNF and YAP-TEAD complexes were shown to be co-dependent for chromatin binding, whereby they create a positive-feedback mechanism to maintain open chromatin for transcription [74]. In contrast, recruitment of the NuRD complex [75] could be involved in the observed co-repressor functions of YAP/TAZ.

In a previous study which employed a transgenic mouse model of cSCC development, simultaneous activation of oncogenic Kras^G12D^ and knock-down of YAP/TAZ in epidermal stem cells was found to induce apoptosis [76]. In contrast, we found that YAP/TAZ depletion in human SCC has cytostatic- and differentiation-promoting, rather than cytotoxic effects, which appear to be mediated by induction of p21. It is worth noting that activating mutations in the RAS proto-oncogenes are not very frequent in human cSCC [2], with some tumours (∼10%) showing copy number loss of HRAS [77]. Induction of apoptosis in response to YAP/TAZ depletion might thus be specific to SCCs expressing oncogenic RAS [76].

We show that a key function of RIF1 in human squamous cancers is to regulate DNA replication dynamics. Our findings in SCC13 cells show that in the absence of RIF1, origin firing and replication fork speed are increased (Fig. 6). However, our DNAscent analysis revealed that the rate of replication fork stalling/pausing in SCC13 cells was not increased by RIF1 depletion. A previous study found that replication stress can be caused by increased fork speed and without fork stalling when the average fork velocity increases above a ∼40% gain [64]. Although the average increase in fork speed in shRIF1 SCC13 cells was only ∼25%, we believe that this increased fork speed can generate increasing levels of replication stress and DNA damage over a few cell division cycles, as was previously shown for cells with elevated E2F-driven transcription [78].

Since origin firing occurs preferentially within open chromatin regions that contain transcriptional-regulatory elements, some of which were shown to be enriched for TEAD TF binding motifs [79], modulation of RIF1 chromatin association by YAP-TEAD (Fig. 4) could serve to coordinate YAP-TEAD-driven transcription with DNA replication. Unfortunately, because of the confounding effects of YAP depletion on the cell cycle (Fig. 1), it was not possible for us to assess if YAP was involved in DNA replication under unperturbed conditions.

An important discovery of our study was a specific role of YAP in facilitating DDR at deactivated replication forks under severe replication stress. How YAP might work together with RIF1 to promote DDR at broken replication forks remains elusive at this point, particularly since the mechanism by which RIF1 enables BRCA1/RAD51-dependent repair of broken replication forks it yet not well understood [23]. Notably, this DDR function is attributed specifically to the long isoform of RIF1 (RIF1-long), while control of replication dynamics is mediated by both RIF1-long and RIF1-short isoforms [80].

Our findings thus provide evidence for a non-canonical, non-transcriptional role of YAP in SCC, whereby it engages RIF1 to dampen the high levels of endogenous replication stress in these cancers, which are—at least in part—caused by YAP’s oncogenic functions itself. Induction of replication stress together with pharmacological YAP/TAZ-TEAD inhibition [73] could thus potentially be used as a combination therapy strategy to treat SCC.

## Acknowledgements

We would like to thank all members of the Walko and Jungwirth labs for their inputs. We wish to thank our colleagues from the Barts Centre for Squamous Cancer at Queen Mary University of London for critical feedback on our draft manuscript. We thank Genomics Birmingham Genomics Service at the University of Birmingham, for the generation of the DNAscent data. We also acknowledge support from the bioimaging facility of the Barts Cancer Institute (Queen Mary University of London).

## Author contributions

Jodie Bojko: Data Curation, Formal Analysis, Investigation, Methodology, Visualization, Writing: Original Draft Preparation; Writing: Review & Editing; Elodie Sins: Data Curation, Investigation, Methodology; Benjamin Flynn: Data Curation, Formal Analysis, Investigation, Methodology, Software, Validation, Visualization; James Scarth: Data Curation, Formal Analysis, Investigation, Methodology, Visualization; Mikal Nigasi: Data Curation, Formal Analysis, Investigation, Methodology, Visualization; Bertram Aschenbrenner: Data Curation, Formal Analysis, Investigation, Methodology, Software, Visualization; Molly Guscott: Formal Analysis, Investigation, Methodology, Software; Alexander Howard: Formal Analysis, Investigation, Methodology; Emily Lay: Formal Analysis, Investigation, Methodology; Emma Bailey: Formal Analysis, Investigation, Methodology, Software, Visualization; Natalia Krajic: Investigation, Methodology; Marco Franciosi: Investigation, Methodology; Sandra Catalan: Investigation, Methodology; Kelli Gallacher: Investigation, Methodology; Ilaria di Girolamo: Data Curation, Investigation; Reem Bagabas: Data Curation, Investigation; Caterina Missero: Funding acquisition, Resources, Supervision, Validation; Jun Wang: Funding acquisition, Methodology, Resources, Supervision; Sarah McClelland: Funding acquisition, Resources, Supervision, Validation; Beate Lichtenberger: Funding acquisition, Resources, Supervision, Validation, Writing: Original Draft Preparation; Ute Jungwirth: Formal Analysis, Funding acquisition, Resources, Supervision, Validation, Visualization, Writing: Original Draft Preparation, Writing: Review & Editing; Gernot Walko: Conceptualization, Data Curation, Formal Analysis, Funding acquisition, Methodology, Project Administration, Resources, Supervision, Validation, Visualization, Writing: Original Draft Preparation, Writing: Review &Editing.

## Data availability

The authors declare that all data supporting the findings of this study are available within the paper and its supplementary information files. Raw Illumina sequencing data have been deposited in the Gene Expression Omnibus (GEO) following the MIAME guidelines, under the accession number GSE284725. The mass spectrometry proteomics data have been deposited to the ProteomeXchange Consortium via the PRIDE partner repository with the dataset identifier PXD061022 and 10.6019/PXD061022. DNAscent data have been deposited on European Nucleotide Archive (ENA), accession ID PRJEB103830.

## References

1 Dotto GP, Rustgi AK. Squamous Cell Cancers: A Unified Perspective on Biology and Genetics. Cancer Cell 2016; 29: 622–637.

2 Winge MCG, Kellman LN, Guo K, Tang JY, Swetter SM, Aasi SZ et al. Advances in cutaneous squamous cell carcinoma. Nat Rev Cancer 2023; 23: 430–449.

3 da Costa A, Chowdhury D, Shapiro GI, D’Andrea AD, Konstantinopoulos PA. Targeting replication stress in cancer therapy. Nat Rev Drug Discov 2023; 22: 38–58.

4 Gaillard H, Garcia-Muse T, Aguilera A. Replication stress and cancer. Nat Rev Cancer 2015; 15: 276–289.

5 Bowry A, Kelly RDW, Petermann E. Hypertranscription and replication stress in cancer. Trends Cancer 2021; 7: 863–877.

6 Molinuevo R, Freije A, Contreras L, Sanz JR, Gandarillas A. The DNA damage response links human squamous proliferation with differentiation. J Cell Biol 2020; 219.

7 Freije A, Ceballos L, Coisy M, Barnes L, Rosa M, De Diego E et al. Cyclin E drives human keratinocyte growth into differentiation. Oncogene 2012; 31: 5180–5192.

8 Freije A, Molinuevo R, Ceballos L, Cagigas M, Alonso-Lecue P, Rodriguez R et al. Inactivation of p53 in Human Keratinocytes Leads to Squamous Differentiation and Shedding via Replication Stress and Mitotic Slippage. Cell Rep 2014; 9: 1349–1360.

9 Gandarillas A, Davies D, Blanchard JM. Normal and c-Myc-promoted human keratinocyte differentiation both occur via a novel cell cycle involving cellular growth and endoreplication. Oncogene 2000; 19: 3278–3289.

10 Alonso-Lecue P, de Pedro I, Coulon V, Molinuevo R, Lorz C, Segrelles C et al. Inefficient differentiation response to cell cycle stress leads to genomic instability and malignant progression of squamous carcinoma cells. Cell Death Dis 2017; 8: e2901.

11 Blasiak J, Szczepanska J, Sobczuk A, Fila M, Pawlowska E. RIF1 Links Replication Timing with Fork Reactivation and DNA Double-Strand Break Repair. Int J Mol Sci 2021; 22.

12 Foti R, Gnan S, Cornacchia D, Dileep V, Bulut-Karslioglu A, Diehl S et al. Nuclear Architecture Organized by Rif1 Underpins the Replication-Timing Program. Mol Cell 2016; 61: 260–273.

13 Gnan S, Flyamer IM, Klein KN, Castelli E, Rapp A, Maiser A et al. Nuclear organisation and replication timing are coupled through RIF1-PP1 interaction. Nat Commun 2021; 12: 2910.

14 Moriyama K, Yoshizawa-Sugata N, Masai H. Oligomer formation and G-quadruplex binding by purified murine Rif1 protein, a key organizer of higher-order chromatin architecture. J Biol Chem 2018; 293: 3607–3624.

15 Cornacchia D, Dileep V, Quivy JP, Foti R, Tili F, Santarella-Mellwig R et al. Mouse Rif1 is a key regulator of the replication-timing programme in mammalian cells. EMBO J 2012; 31: 3678–3690.

16 Alver RC, Chadha GS, Gillespie PJ, Blow JJ. Reversal of DDK-Mediated MCM Phosphorylation by Rif1-PP1 Regulates Replication Initiation and Replisome Stability Independently of ATR/Chk1. Cell Rep 2017; 18: 2508–2520.

17 Moiseeva TN, Yin Y, Calderon MJ, Qian C, Schamus-Haynes S, Sugitani N et al. An ATR and CHK1 kinase signaling mechanism that limits origin firing during unperturbed DNA replication. Proc Natl Acad Sci U S A 2019; 116: 13374–13383.

18 Yamazaki S, Ishii A, Kanoh Y, Oda M, Nishito Y, Masai H. Rif1 regulates the replication timing domains on the human genome. EMBO J 2012; 31: 3667–3677.

19 Hiraga SI, Ly T, Garzon J, Horejsi Z, Ohkubo YN, Endo A et al. Human RIF1 and protein phosphatase 1 stimulate DNA replication origin licensing but suppress origin activation. EMBO Rep 2017; 18: 403–419.

20 Buonomo SB, Wu Y, Ferguson D, de Lange T. Mammalian Rif1 contributes to replication stress survival and homology-directed repair. J Cell Biol 2009; 187: 385–398.

21 Mukherjee C, Tripathi V, Manolika EM, Heijink AM, Ricci G, Merzouk S et al. RIF1 promotes replication fork protection and efficient restart to maintain genome stability. Nat Commun 2019; 10: 3287.

22 Garzon J, Ursich S, Lopes M, Hiraga SI, Donaldson AD. Human RIF1-Protein Phosphatase 1 Prevents Degradation and Breakage of Nascent DNA on Replication Stalling. Cell Rep 2019; 27: 2558–2566 e2554.

23 Dong Q, Day M, Saito Y, Parker E, Watts LP, Kanemaki MT et al. The human RIF1-Long isoform interacts with BRCA1 to promote recombinational fork repair under DNA replication stress. Nat Commun 2025; 16: 5820.

24 Wang H, Zhao A, Chen L, Zhong X, Liao J, Gao M et al. Human RIF1 encodes an anti-apoptotic factor required for DNA repair. Carcinogenesis 2009; 30: 1314–1319.

25 Eke I, Zong D, Aryankalayil MJ, Sandfort V, Bylicky MA, Rath BH et al. 53BP1/RIF1 signaling promotes cell survival after multifractionated radiotherapy. Nucleic Acids Res 2020; 48: 1314–1326.

26 Mei Y, Peng C, Liu YB, Wang J, Zhou HH. Silencing RIF1 decreases cell growth, migration and increases cisplatin sensitivity of human cervical cancer cells. Oncotarget 2017; 8: 107044–107051.

27 Noordermeer SM, Adam S, Setiaputra D, Barazas M, Pettitt SJ, Ling AK et al. The shieldin complex mediates 53BP1-dependent DNA repair. Nature 2018; 560: 117–121.

28 Franklin JM, Wu Z, Guan KL. Insights into recent findings and clinical application of YAP and TAZ in cancer. Nat Rev Cancer 2023; 23: 512–525.

29 Howard A, Bojko J, Flynn B, Bowen S, Jungwirth U, Walko G. Targeting the Hippo/YAP/TAZ signalling pathway: Novel opportunities for therapeutic interventions into skin cancers. Exp Dermatol 2022; 31: 1477–1499.

30 Maehama T, Nishio M, Otani J, Mak TW, Suzuki A. The role of Hippo-YAP signaling in squamous cell carcinomas. Cancer Sci 2021; 112: 51–60.

31 Hillmer RE, Link BA. The Roles of Hippo Signaling Transducers Yap and Taz in Chromatin Remodeling. Cells 2019; 8.

32 Walko G, Woodhouse S, Pisco AO, Rognoni E, Liakath-Ali K, Lichtenberger BM et al. A genome-wide screen identifies YAP/WBP2 interplay conferring growth advantage on human epidermal stem cells. Nat Commun 2017; 8: 14744.

33 De Rosa L, Secone Seconetti A, De Santis G, Pellacani G, Hirsch T, Rothoeft T et al. Laminin 332-Dependent YAP Dysregulation Depletes Epidermal Stem Cells in Junctional Epidermolysis Bullosa. Cell Rep 2019; 27: 2036–2049 e2036.

34 Lavado A, Park JY, Pare J, Finkelstein D, Pan H, Xu B et al. The Hippo Pathway Prevents YAP/TAZ-Driven Hypertranscription and Controls Neural Progenitor Number. Dev Cell 2018; 47: 576–591 e578.

35 Gertzmann D, Presek C, Mattes AL, Sanger M, Zoller M, Schulein-Volk C et al. Oncogenic YAP sensitizes cells to CHK1 inhibition via CDK4/6 driven G1 acceleration. EMBO Rep 2025.

36 Yuan Y, Park J, Feng A, Awasthi P, Wang Z, Chen Q et al. YAP1/TAZ-TEAD transcriptional networks maintain skin homeostasis by regulating cell proliferation and limiting KLF4 activity. Nat Commun 2020; 11: 1472.

37 Rheinwald JG, Beckett MA. Tumorigenic keratinocyte lines requiring anchorage and fibroblast support cultured from human squamous cell carcinomas. Cancer Res 1981; 41: 1657–1663.

38 Zanconato F, Forcato M, Battilana G, Azzolin L, Quaranta E, Bodega B et al. Genome-wide association between YAP/TAZ/TEAD and AP-1 at enhancers drives oncogenic growth. Nat Cell Biol 2015; 17: 1218–1227.

39 Pattschull G, Walz S, Grundl M, Schwab M, Ruhl E, Baluapuri A et al. The Myb-MuvB Complex Is Required for YAP-Dependent Transcription of Mitotic Genes. Cell Rep 2019; 27: 3533–3546 e3537.

40 Lopez-Pajares V, Qu K, Zhang J, Webster DE, Barajas BC, Siprashvili Z et al. A LncRNA-MAF:MAFB transcription factor network regulates epidermal differentiation. Dev Cell 2015; 32: 693–706.

41 Kypriotou M, Huber M, Hohl D. The human epidermal differentiation complex: cornified envelope precursors, S100 proteins and the ’fused genes’ family. Exp Dermatol 2012; 21: 643–649.

42 Wang J, Harwood CA, Bailey E, Bewicke-Copley F, Anene CA, Thomson J et al. Transcriptomic analysis of cutaneous squamous cell carcinoma reveals a multigene prognostic signature associated with metastasis. J Am Acad Dermatol 2023; 89: 1159–1166.

43 Hagenbeek TJ, Zbieg JR, Hafner M, Mroue R, Lacap JA, Sodir NM et al. An allosteric pan-TEAD inhibitor blocks oncogenic YAP/TAZ signaling and overcomes KRAS G12C inhibitor resistance. Nat Cancer 2023; 4: 812–828.

44 Yuan Y, Salinas Parra N, Chen Q, Iglesias-Bartolome R. Oncogenic Hedgehog-smoothened signaling depends on YAP1-TAZ/TEAD transcription to restrain differentiation in basal cell carcinoma. J Invest Dermatol 2021.

45 Ashley AK, Shrivastav M, Nie J, Amerin C, Troksa K, Glanzer JG et al. DNA-PK phosphorylation of RPA32 Ser4/Ser8 regulates replication stress checkpoint activation, fork restart, homologous recombination and mitotic catastrophe. DNA Repair (Amst) 2014; 21: 131–139.

46 Liu S, Opiyo SO, Manthey K, Glanzer JG, Ashley AK, Amerin C et al. Distinct roles for DNA-PK, ATM and ATR in RPA phosphorylation and checkpoint activation in response to replication stress. Nucleic Acids Res 2012; 40: 10780–10794.

47 Dempster JM, Pacini C, Pantel S, Behan FM, Green T, Krill-Burger J et al. Agreement between two large pan-cancer CRISPR-Cas9 gene dependency data sets. Nat Commun 2019; 10: 5817.

48 Takahashi N, Kim S, Schultz CW, Rajapakse VN, Zhang Y, Redon CE et al. Replication stress defines distinct molecular subtypes across cancers. Cancer Res Commun 2022; 2: 503–517.

49 Kotsantis P, Silva LM, Irmscher S, Jones RM, Folkes L, Gromak N et al. Increased global transcription activity as a mechanism of replication stress in cancer. Nat Commun 2016; 7: 13087.

50 Mohammed H, Taylor C, Brown GD, Papachristou EK, Carroll JS, D’Santos CS. Rapid immunoprecipitation mass spectrometry of endogenous proteins (RIME) for analysis of chromatin complexes. Nat Protoc 2016; 11: 316–326.

51 Zhang N, Bai H, David KK, Dong J, Zheng Y, Cai J et al. The Merlin/NF2 tumor suppressor functions through the YAP oncoprotein to regulate tissue homeostasis in mammals. Dev Cell 2010; 19: 27–38.

52 Zhou Y, Zhang J, Li H, Huang T, Wong CC, Wu F et al. AMOTL1 enhances YAP1 stability and promotes YAP1-driven gastric oncogenesis. Oncogene 2020; 39: 4375–4389.

53 Wang P, Bai Y, Song B, Wang Y, Liu D, Lai Y et al. PP1A-mediated dephosphorylation positively regulates YAP2 activity. PLoS One 2011; 6: e24288.

54 Zhu M, Peng R, Liang X, Lan Z, Tang M, Hou P et al. P4HA2-induced prolyl hydroxylation suppresses YAP1-mediated prostate cancer cell migration, invasion, and metastasis. Oncogene 2021; 40: 6049–6056.

55 Mittal P, Roberts CWM. The SWI/SNF complex in cancer - biology, biomarkers and therapy. Nat Rev Clin Oncol 2020; 17: 435–448.

56 He L, Pratt H, Gao M, Wei F, Weng Z, Struhl K. YAP and TAZ are transcriptional co-activators of AP-1 proteins and STAT3 during breast cellular transformation. Elife 2021; 10.

57 Ning B, Tilston-Lunel AM, Simonetti J, Hicks-Berthet J, Matschulat A, Pfefferkorn R et al. Convergence of YAP/TAZ, TEAD and TP63 activity is associated with bronchial premalignant severity and progression. J Exp Clin Cancer Res 2023; 42: 116.

58 Nowak DE, Tian B, Brasier AR. Two-step cross-linking method for identification of NF-kappaB gene network by chromatin immunoprecipitation. Biotechniques 2005; 39: 715–725.

59 Melendez Garcia R, Haccard O, Chesneau A, Narassimprakash H, Roger J, Perron M et al. A non-transcriptional function of Yap regulates the DNA replication program in Xenopus laevis. Elife 2022; 11.

60 Proby CM, Purdie KJ, Sexton CJ, Purkis P, Navsaria HA, Stables JN et al. Spontaneous keratinocyte cell lines representing early and advanced stages of malignant transformation of the epidermis. Exp Dermatol 2000; 9: 104–117.

61 Chandrashekar DS, Karthikeyan SK, Korla PK, Patel H, Shovon AR, Athar M et al. UALCAN: An update to the integrated cancer data analysis platform. Neoplasia 2022; 25: 18–27.

62 Zhang Y, Chen F, Chandrashekar DS, Varambally S, Creighton CJ. Proteogenomic characterization of 2002 human cancers reveals pan-cancer molecular subtypes and associated pathways. Nat Commun 2022; 13: 2669.

63 Jones MJK, Rai SK, Pfuderer PL, Bonfim-Melo A, Pagan JK, Clarke PR et al. A high-resolution, nanopore-based artificial intelligence assay for DNA replication stress in human cancer cells. Nat Commun 2025; 16: 7732.

64 Maya-Mendoza A, Moudry P, Merchut-Maya JM, Lee M, Strauss R, Bartek J. High speed of fork progression induces DNA replication stress and genomic instability. Nature 2018; 559: 279–284.

65 Raso MC, Djoric N, Walser F, Hess S, Schmid FM, Burger S et al. Interferon-stimulated gene 15 accelerates replication fork progression inducing chromosomal breakage. J Cell Biol 2020; 219.

66 Fousek-Schuller VJ, Borgstahl GEO. The Intriguing Mystery of RPA Phosphorylation in DNA Double-Strand Break Repair. Genes (Basel) 2024; 15.

67 Toledo LI, Altmeyer M, Rask MB, Lukas C, Larsen DH, Povlsen LK et al. ATR prohibits replication catastrophe by preventing global exhaustion of RPA. Cell 2013; 155: 1088–1103.

68 Picco G, Chen ED, Alonso LG, Behan FM, Goncalves E, Bignell G et al. Functional linkage of gene fusions to cancer cell fitness assessed by pharmacological and CRISPR-Cas9 screening. Nat Commun 2019; 10: 2198.

69 Petermann E, Orta ML, Issaeva N, Schultz N, Helleday T. Hydroxyurea-stalled replication forks become progressively inactivated and require two different RAD51-mediated pathways for restart and repair. Mol Cell 2010; 37: 492–502.

70 Moeglin E, Desplancq D, Conic S, Oulad-Abdelghani M, Stoessel A, Chiper M et al. Uniform Widespread Nuclear Phosphorylation of Histone H2AX Is an Indicator of Lethal DNA Replication Stress. Cancers (Basel) 2019; 11.

71 Cui P, Lin Q, Xin C, Han L, An L, Wang Y et al. Hydroxyurea-induced global transcriptional suppression in mouse ES cells. Carcinogenesis 2010; 31: 1661–1668.

72 Li Q, Tie Y, Alu A, Ma X, Shi H. Targeted therapy for head and neck cancer: signaling pathways and clinical studies. Signal Transduct Target Ther 2023; 8: 31.

73 Sato K, Faraji F, Cervantes-Villagrana RD, Wu X, Koshizuka K, Ishikawa T et al. Targeting YAP/TAZ-TEAD signaling as a therapeutic approach in head and neck squamous cell carcinoma. Cancer Lett 2025: 217467.

74 Chang CY, Shipony Z, Lin SG, Kuo A, Xiong X, Loh KM et al. Increased ACTL6A occupancy within mSWI/SNF chromatin remodelers drives human squamous cell carcinoma. Mol Cell 2021; 81: 4964–4978 e4968.

75 Kim M, Kim T, Johnson RL, Lim DS. Transcriptional co-repressor function of the hippo pathway transducers YAP and TAZ. Cell Rep 2015; 11: 270–282.

76 Debaugnies M, Sanchez-Danes A, Rorive S, Raphael M, Liagre M, Parent MA et al. YAP and TAZ are essential for basal and squamous cell carcinoma initiation. EMBO Rep 2018.

77 Inman GJ, Wang J, Nagano A, Alexandrov LB, Purdie KJ, Taylor RG et al. The genomic landscape of cutaneous SCC reveals drivers and a novel azathioprine associated mutational signature. Nat Commun 2018; 9: 3667.

78 Pennycook BR, Vesela E, Peripolli S, Singh T, Barr AR, Bertoli C et al. E2F-dependent transcription determines replication capacity and S phase length. Nat Commun 2020; 11: 3503.

79 Kumagai A, Dunphy WG. Binding of the Treslin-MTBP Complex to Specific Regions of the Human Genome Promotes the Initiation of DNA Replication. Cell Rep 2020; 32: 108178.

80 Watts LP, Natsume T, Saito Y, Garzon J, Dong Q, Boteva L et al. The RIF1-long splice variant promotes G1 phase 53BP1 nuclear bodies to protect against replication stress. Elife 2020; 9.

