## Supplementary Figures for "YAP engages RIF1 to dampen replication stress-induced DNA damage in human squamous cell carcinoma"

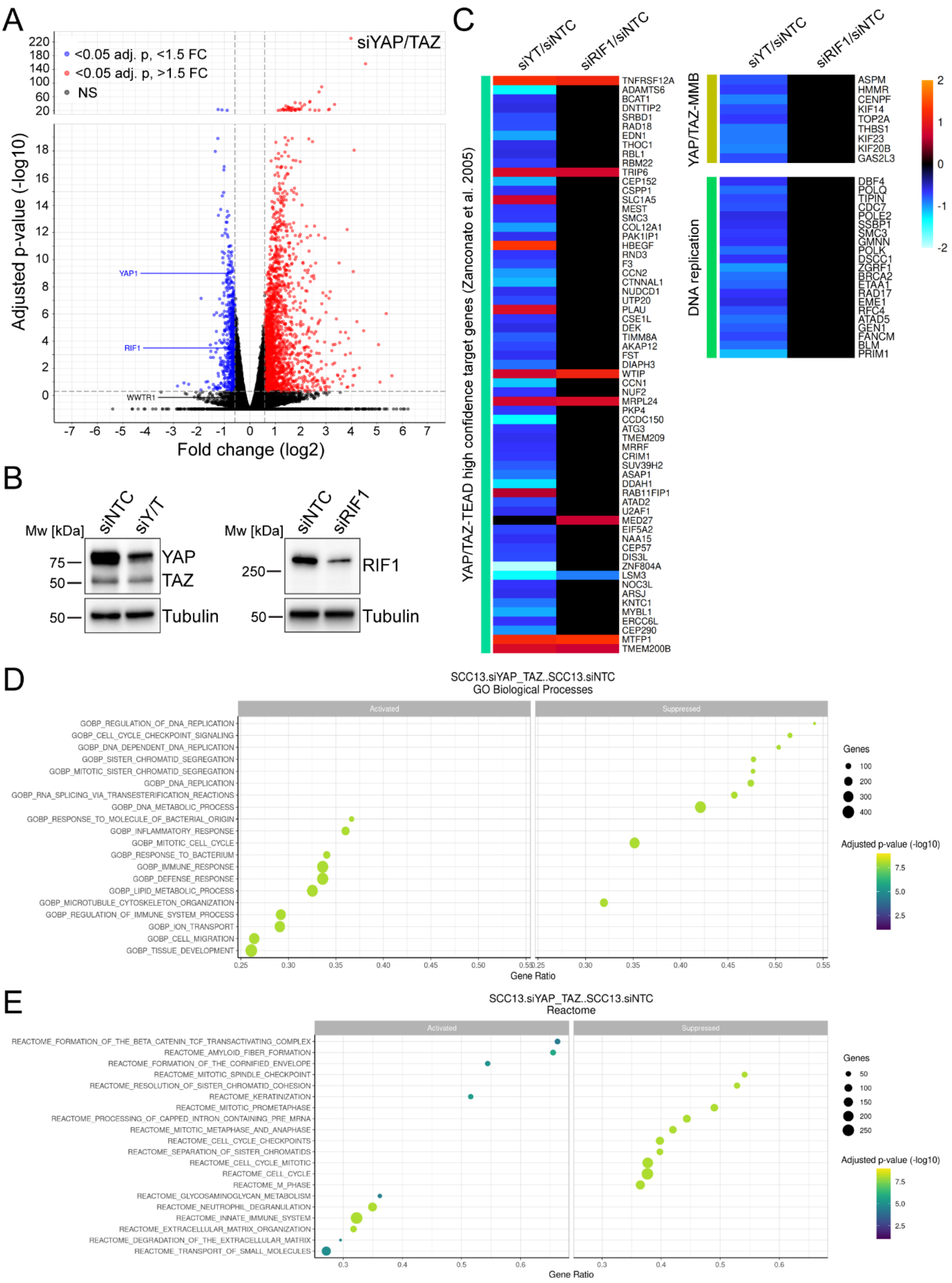

***Supplementary Figure 1. The YAP/TAZ-regulated transcriptome in SCC13 cells.***

(**A**) Significantly enriched or depleted RNAs in siYAP/TAZ SCC13 cells identified by RNA-Seq analysis using DESeq2. Volcano plot shows statistical significance (-log 10 (adj. p-value)) versus relative RNA abundance (log2-fold change) in siYAP/TAZ compared to siNTC cells. Vertical dotted grey lines: log2-fold change cut-off (0.58, 1.5-fold change); horizontal grey dotted lines: cut-off for statistical significance ($p<0.05$). (**B**) Immunoblot analysis of SCC13 cells expressing siNTC, siYAP/TAZ (siY/T), or siRIF1 for 48 h, using the indicated antibodies. Tubulin was used as loading control. (**C**) Heatmaps show the log2-fold change of selected genes representing high confidence YAP/TAZ-TEAD target genes, YAP-MMB target genes, and genes directly involved in DNA replication, in the indicated pairwise comparisons between siNTC and siYAP/TAZ or siNTC and siRIF1 SCC13 cells. Black colour indicates that the gene is not differentially regulated in that pairwise comparison. (**D**, **E**) Bubble plots show results of gene set enrichment analysis (GSEA) on RNA-seq data from siYAP/TAZ SCC13 cells. The top 10 downregulated (suppressed) or upregulated (activated) pathways in siYAP/TAZ SCC13 cells are shown.

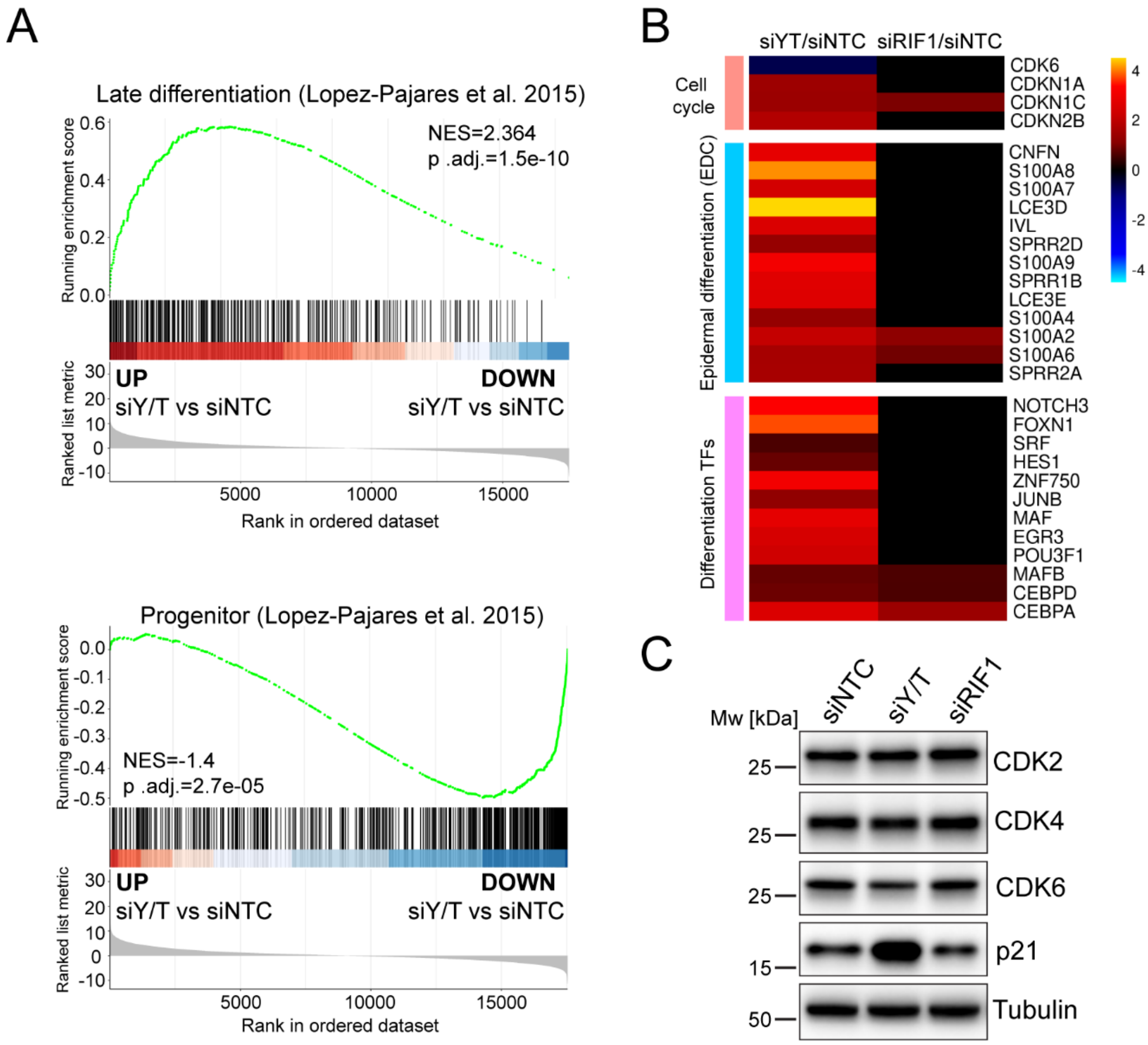


***Supplementary Figure 2. YAP/TAZ suppress terminal differentiation commitment.***

(**A**) GSEA showing enrichment of epidermal late differentiation (upper panel) and progenitor cell (lower panel) signatures [40] among the differentially expressed genes in siYAP/TAZ (siY/T) versus siNTC SCC13 cells. (**B**) Heatmaps show the log2-fold change of selected genes representing key cell cycle regulatory genes, transcription factors (TFs) regulating terminal differentiation, and genes belonging to epidermal differentiation complex, in the indicated pairwise comparisons between siNTC and siYAP/TAZ or siNTC and siRIF1 SCC13 cells. Black colour indicates that the gene is not differentially regulated in that pairwise comparison. (**C**) Immunoblot analysis of SCC13 cells expressing siNTC, siYAP/TAZ (siY/T), or siRIF1 for 48 h, using the indicated antibodies. Tubulin was used as loading control.

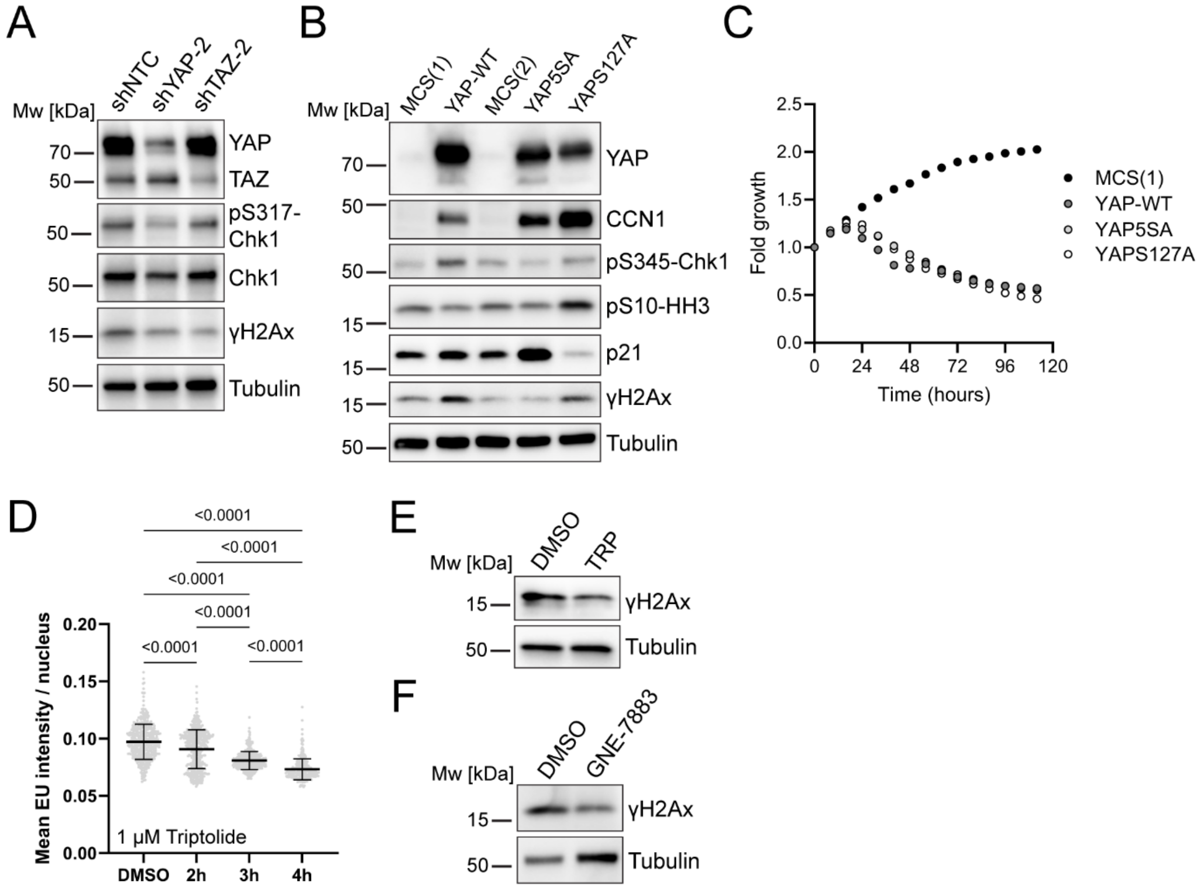


***Supplementary Figure 3. YAP/TAZ-TEAD-mediated oncogenic signalling is major contributor to replication stress in squamous cell carcinoma cells.***

(**A**) Immunoblot analysis of SCC13 cells expressing doxycycline-induced shNTC, shYAP, or shTAZ using the indicated antibodies. Tubulin was used as loading control. (**B**) Immunoblot analysis of SCC cells expressing empty vector controls (MCS) or the different YAP transgenes, using the indicated antibodies. YAP transgene expression was induced for 48 h prior to cell lysis. Tubulin was used as loading control. MCS(1) and MCS(2) refer to two different empty vector control lines. (**C**) Proliferation kinetics of cells in (B). (**D**) Quantification of nascent RNA synthesis by EU incorporation in cells treated with transcription inhibitor triptolide. n=406 cells (2h), n=452 cells (3h), n=357 cells (3h), n=283 cells (4h), in n=1 experiment. Exact p-values are shown, Kruskal-Wallis test with Dunn's multiple comparison test. (**E, F**)

Immunoblot analysis of SCC13 cells treated with DMSO, triptolide (TRP, 1 μM), or GNE-7883 (2 μM), using the indicated antibodies. Tubulin was used as loading control.

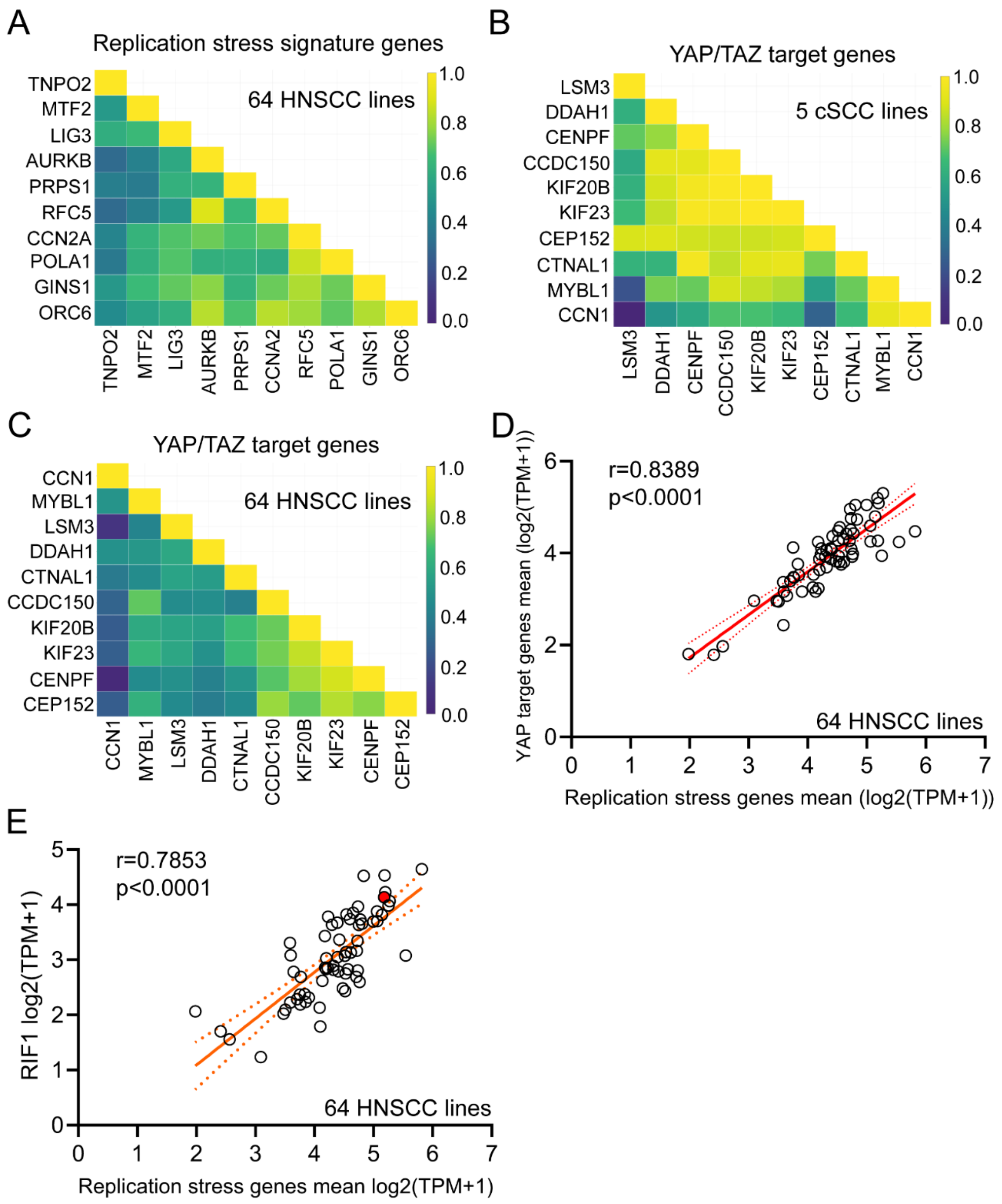

***Supplementary Figure 4. Correlation analysis of YAP/TAZ target genes, replication stress-associated genes, and RIF1 expression in HNSCC cell lines.***

(**A**–**C**) Correlation heatmaps of the indicated genes (mean mRNA expression) distinguished by (A, C) 64 HNSCC or (B) 5 cSCC cell lines. Graphs were exported from the DepMap portal. (**D, E**) Correlation analysis between the expression of replication stress-associated genes and (D) YAP/TAZ target genes or (E) *RIF1* mRNA expression levels (expression public 24Q2 release), in a panel of 64 HNSCC cell lines. Spearman correlation coefficient and statistical significance are indicated. Data were accessed using the DepMap portal (https://depmap.org/portal/). The red-coloured circle in (E) indicates the SAS cell line used in this study.

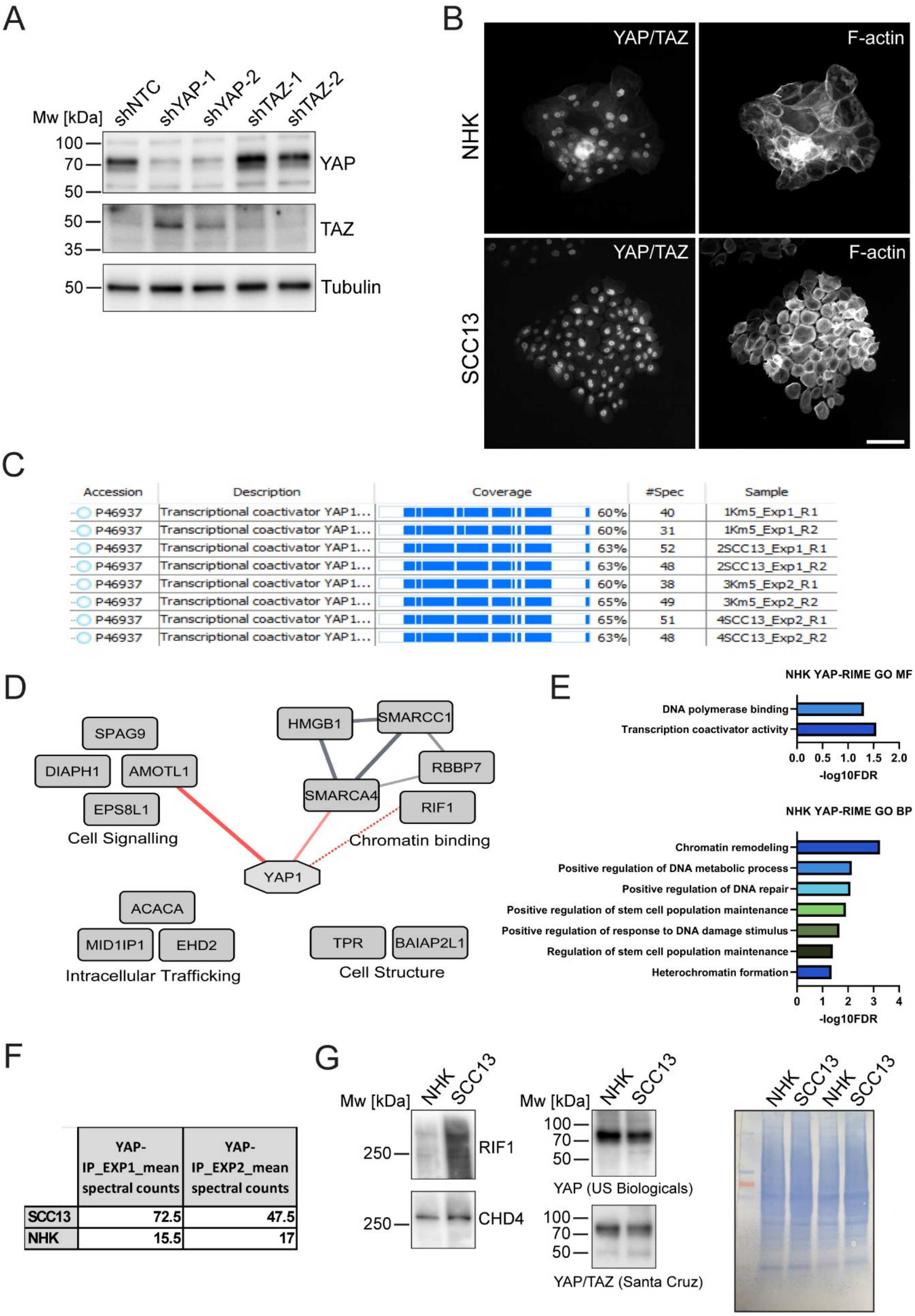
A
Mw [kDa]
shNTC
shYAP-1
shYAP-2
shTAZ-1
shTAZ-2
100
70
50
YAP
50
35
TAZ
50
Tubulin
B
YAP/TAZ
F-actin
NHK
SCC13
C
Accession | Description | Coverage | #Spec | Sample
P46937 | Transcriptional coactivator YAP1... | 60% | 40 | 1Km5_Exp1_R1
P46937 | Transcriptional coactivator YAP1... | 60% | 31 | 1Km5_Exp1_R2
P46937 | Transcriptional coactivator YAP1... | 63% | 52 | 2SCC13_Exp1_R1
P46937 | Transcriptional coactivator YAP1... | 63% | 48 | 2SCC13_Exp1_R2
P46937 | Transcriptional coactivator YAP1... | 60% | 38 | 3Km5_Exp2_R1
P46937 | Transcriptional coactivator YAP1... | 65% | 49 | 3Km5_Exp2_R2
P46937 | Transcriptional coactivator YAP1... | 65% | 51 | 4SCC13_Exp2_R1
P46937 | Transcriptional coactivator YAP1... | 63% | 48 | 4SCC13_Exp2_R2
D
SPAG9
DIAPH1
AMOTL1
EPS8L1
Cell Signalling
HMGB1
SMARCC1
RBBP7
SMARCA4
RIF1
Chromatin binding
YAP1
ACACA
MID1IP1
EHD2
Intracellular Trafficking
TPR
BAIAP2L1
Cell Structure
E
NHK YAP-RIME GO MF
DNA polymerase binding
Transcription coactivator activity
0.0 0.5 1.0 1.5 2.0
-log10FDR
NHK YAP-RIME GO BP
Chromatin remodeling
Positive regulation of DNA metabolic process
Positive regulation of DNA repair
Positive regulation of stem cell population maintenance
Positive regulation of response to DNA damage stimulus
Regulation of stem cell population maintenance
Heterochromatin formation
0 1 2 3 4
-log10FDR
F
| YAP-IP_EXP1_mean spectral counts | YAP-IP_EXP2_mean spectral counts
SCC13 | 72.5 | 47.5
NHK | 15.5 | 17
G
Mw [kDa]
NHK
SCC13
250
RIF1
250
CHD4
100
70
50
YAP (US Biologicals)
100
70
50
YAP/TAZ (Santa Cruz)
NHK
SCC13
NHK
SCC13

***Supplementary Figure 5. RIME identifies RIF1 as an interactor of chromatin-associated YAP in normal human keratinocytes.***

(**A**) Immunoblot analysis of cell extracts from SCC13 cells expressing a non-targeting control shRNA (shNTC), two different shRNAs targeting YAP (shYAP-1, shYAP-2), or two different shRNAs targeting TAZ (shTAZ-1, shTAZ-2) using the indicated antibodies. Note that the rabbit anti-human YAP antibody used for RIME (Y1200-01D, US Biologicals) did not recognize TAZ (Mw YAP ~78 kDa, Mw TAZ ~50 kDa). Tubulin was used as loading control. (**B**) Representative wide-field microscopy images of YAP nuclear localisation in NHK and SCC13 cells prior to RIME experiments. F-actin was counterstained with phalloidin. Scale bar, 100 µm. (**C**) Coverage of bait protein YAP across both experiments and all technical replicates. Regions shown in blue represent peptides detected by mass spectrometry. Km5 = NHK cells, strain km, passage 5. (**D**) Interaction network of YAP protein interactors identified by RIME in NHKs. Red connecting lines (edges) show previously identified YAP interactions according to the STRING Homo sapiens database, and nodes with red borders indicate known YAP interactors (manually curated from STRING and literature search); dashed red lines show manually curated known YAP interactions that were not yet present in the STRING database. Proteins were manually grouped based on function (gene names displayed, generated in Cytoscape 3.9.1 using STRING app 1.7.1). (**E**) Gene ontology terms ('molecular functions' and 'biological process') enriched within the high-confidence YAP-interacting proteins identified by RIME in NHKs. (**F**) Mean spectral counts (total number of identified peptide spectra per protein) for YAP and RIF1 in the YAP-RIME from the two independently performed experiments in SCC13 and NHK cells. (**G**) Immunoblot analysis of chromatin fractions from NHK and SCC13 cells, using the indicated antibodies. Similar amounts of total chromatin-associated proteins were loaded on SDS-PAGE, as validated by Coomassie staining.

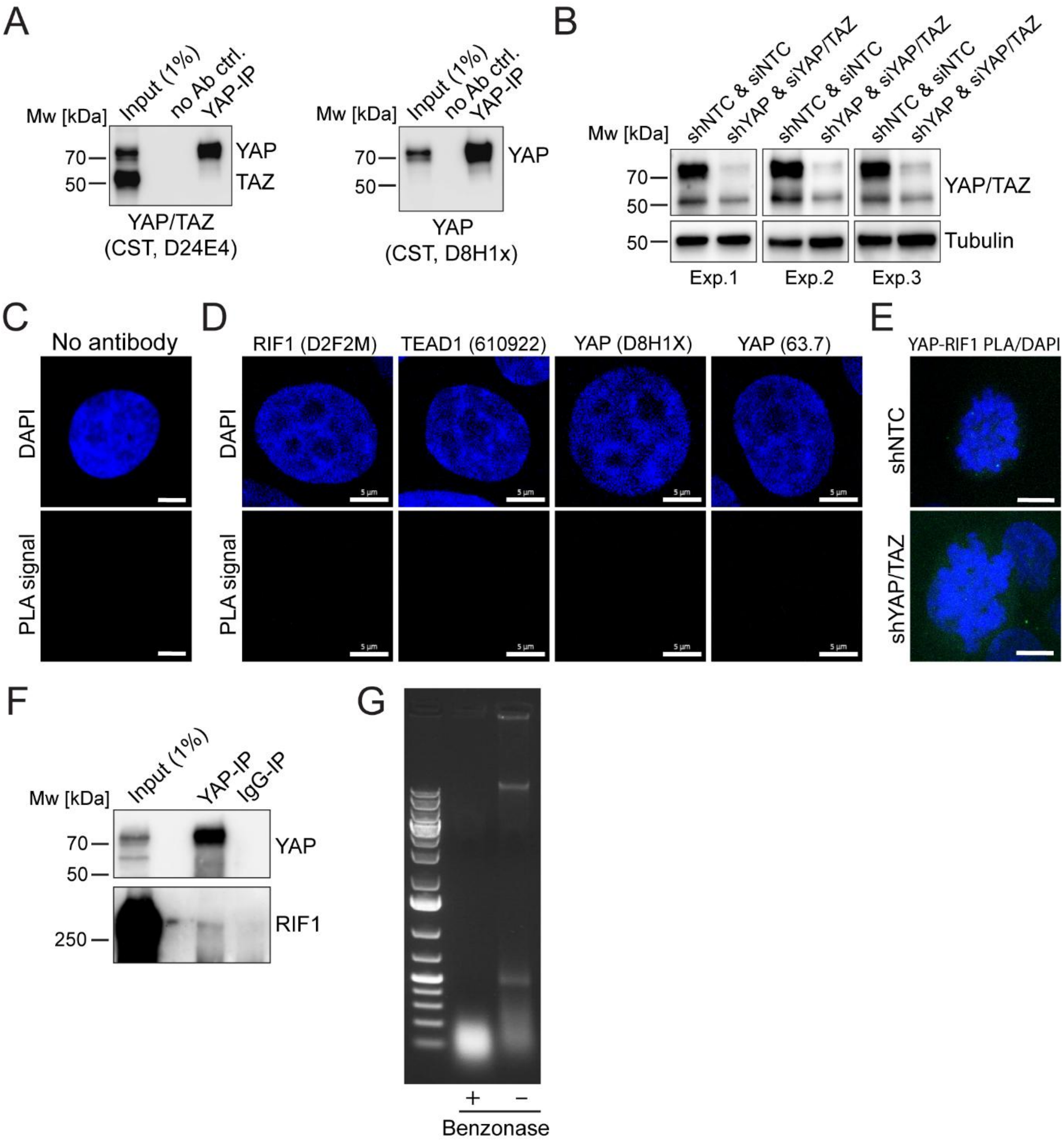


***Supplementary Figure 6. Validation of YAP/TAZ knockdown efficiency and antibody specificity for in-situ PLA experiments, and confirmation of YAP-RIF1 interaction in HARA cells.***

(**A**) Immunoblot analysis of YAP (rabbit monoclonal anti-YAP antibody; D8H1X, Cell Signalling)) and IgG-control immunoprecipitates and respective inputs from SCC13 cell extracts using the indicated antibodies. (**B**) Immunoblot analysis of lysates from cells used in in-situ PLA experiments shown in Figures 3F, G. To achieve the strongest YAP/TAZ depletion possible, SCC13 cells expressing inducible shNTC or shYAP were treated with doxycycline for 72 h and were in parallel also transfected with non-targeting control or YAP/TAZ-targeting siRNA SMARTpools (10 nM), respectively. Tubulin was used as loading control. (**C**, **D**) Representative confocal microscopy images (z-projections) of in-situ PLA (PLA signal, green; DNA (DAPI) blue) in SCC13 cells labelled with (C) no primary antibodies or (D) only one of the two primary antibodies. Scale bars, 5 µm. (**E**) Representative confocal microscopy images (z-projections) of YAP-RIF1 in-situ PLA (PLA signal, green; DNA (DAPI) blue) in miotic cells. Scale bars, 5 µm. (**F**) Immunoblot analysis of YAP and IgG-control immunoprecipitates and respective inputs from HARA cell nuclei-enriched fractions (treated with Benzonase® nuclease) using the indicated antibodies. (**G**) Agarose gel electrophoresis of DNA extracts from Benzonase®-treated and untreated HARA cells shows efficient DNA degradation upon Benzonase® treatment. Left lane: Thermo Scientific™ GeneRuler 1 kb Plus DNA Ladder (Fisher, UK).

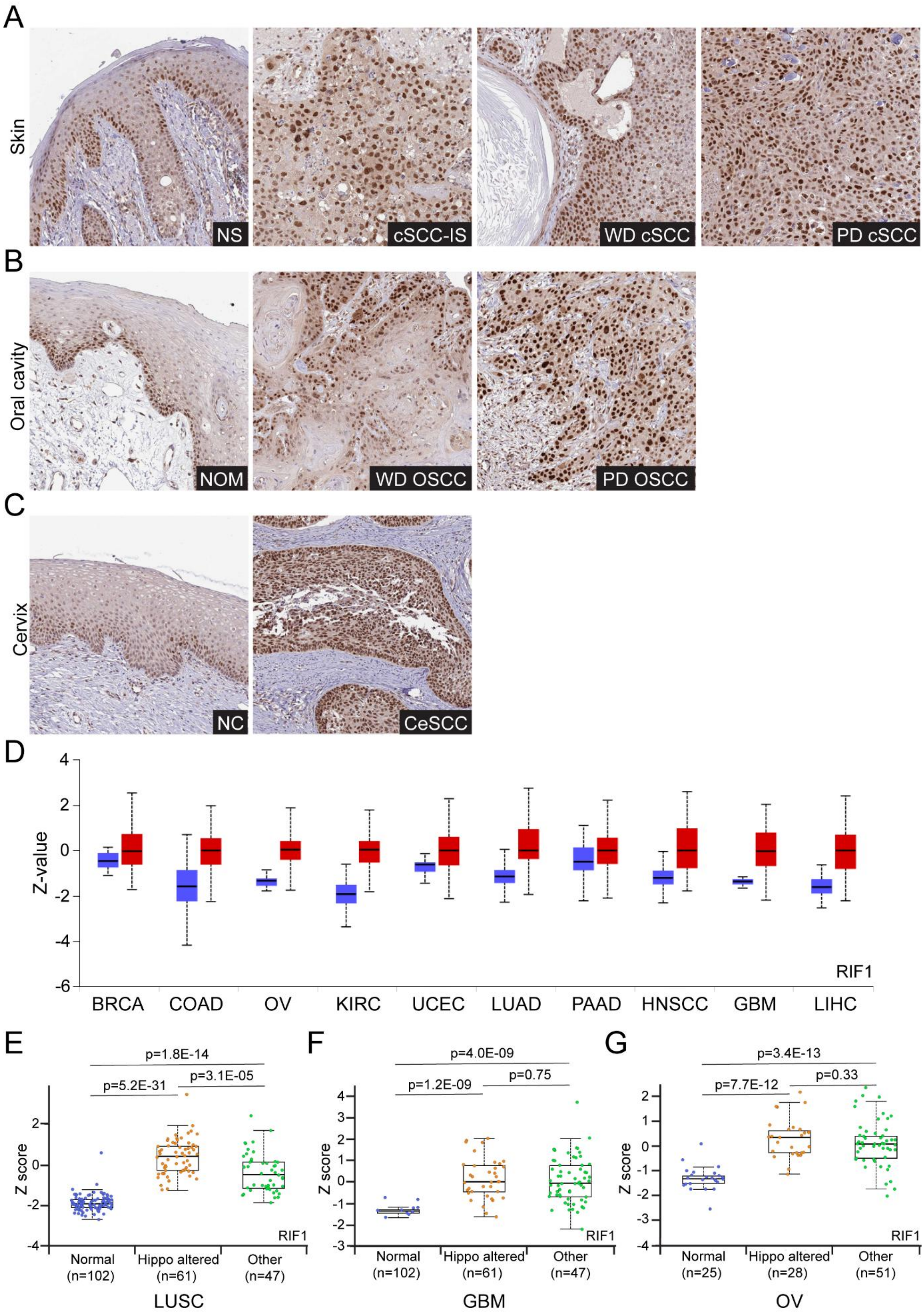
A
Skin
NS
cSCC-IS
WD cSCC
PD cSCC
B
Oral cavity
NOM
WD OSCC
PD OSCC
C
Cervix
NC
CeSCC
D
Z-value
RIF1
BRCA
COAD
OV
KIRC
UCEC
LUAD
PAAD
HNSCC
GBM
LIHC
E
p=1.8E-14
p=5.2E-31
p=3.1E-05
Z score
RIF1
Normal (n=102)
Hippo altered (n=61)
Other (n=47)
LUSC
F
p=4.0E-09
p=1.2E-09
p=0.75
Z score
RIF1
Normal (n=102)
Hippo altered (n=61)
Other (n=47)
GBM
G
p=3.4E-13
p=7.7E-12
p=0.33
Z score
RIF1
Normal (n=25)
Hippo altered (n=28)
Other (n=51)
OV

***Supplementary Figure 7. RIF1 expression in different types of squamous cancers.***

(**A**–**C**) Representative RIF1 immunohistochemistry (anti-RIF1 rabbit polyclonal antibody HPA036887, Atlas Antibodies) of normal tissue and squamous cell carcinoma tissue from (A) skin, (B) oral cavity and (C) cervix, exported from the Human Protein Atlas database (https://www.proteinatlas.org/). NS, normal skin; cSCC-IS, cutaneous squamous cell carcinoma in-situ; WD cSCC, well differentiated cSCC; PD cSCC, poorly differentiated cSCC; NOM, normal oral mucosa; WD OSCC, well differentiated oral SCC; PD, poorly differentiated oral SCC; NC, normal cervix; CeSCC, cervical SCC. (**D**) RIF1 protein expression across different cancers, with tumour (red) and normal (blue) samples, from the Clinical Proteomic Tumour Analysis Consortium (CPTAC). Box plots indicate median Z-values (middle line), 25$^{th}$ percentile (bottom line), 75$^{th}$ percentile (top line), and minimum and maximum (whiskers). Log2 spectral count ratio values from CPTAC were first normalized within each sample profile, then normalized across samples. Graphs were downloaded from the UALCAN database (https://ualcan.path.uab.edu/). BRCA, breast cancer (18 normal and 125 tumour samples); COAD, colon adenocarcinoma (100 normal and 97 tumour samples); OV, ovarian serous cystadenocarcinoma (25 normal and 100 tumour samples); KIRC, kidney renal clear cell carcinoma (84 normal and 110 tumour samples); UCEC, uterine corpus endometrial carcinoma (31 normal and 100 tumour samples); LUAD, lung adenocarcinoma (111 normal and 111 tumour samples); PAAD, pancreatic adenocarcinoma (74 normal and 137 tumour samples); HNSCC, head and neck squamous cell carcinoma (71 normal and 108 tumour samples); GBM, glioblastoma multiforme (10 normal and 99 tumour samples); LIHC, liver hepatocellular carcinoma (165 normal and 165 tumour samples). (**E**–**G**) RIF1 protein expression in (E) LUSC, (F) GMB, and (G) OV samples from the Clinical Proteomic Tumour Analysis Consortium (CPTAC), stratified according to presence of alterations in the Hippo pathway or other signalling pathways. Jitter plots (overlayed onto box plots) show Z-values (standard deviations from the median across samples for the given sample type) from individual samples. Box plots indicate the median (middle line), 25th percentile (bottom line), 75th percentile (top line), and minimum and maximum (whiskers). Log2 spectral count ratio values from CPTAC were first normalized within each sample profile, then normalized across samples. Graphs were downloaded from the UALCAN database (https://ualcan.path.uab.edu/). Exact p-values are shown.

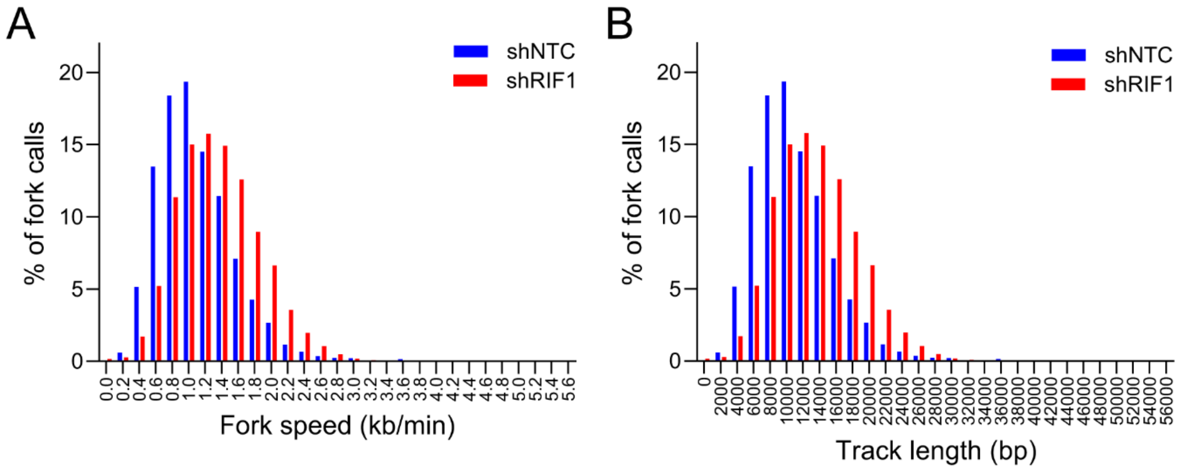


***Supplementary Figure 8. RIF1 depletion in SCC13 cells increases fork speed.***

(**A, B**) Distribution of replication fork speed (A) and replication track length (B) measured by DNAscent. Total number of forks scored in n=2 independent experiments: shNTC = 5632; shRIF1 = 6216.

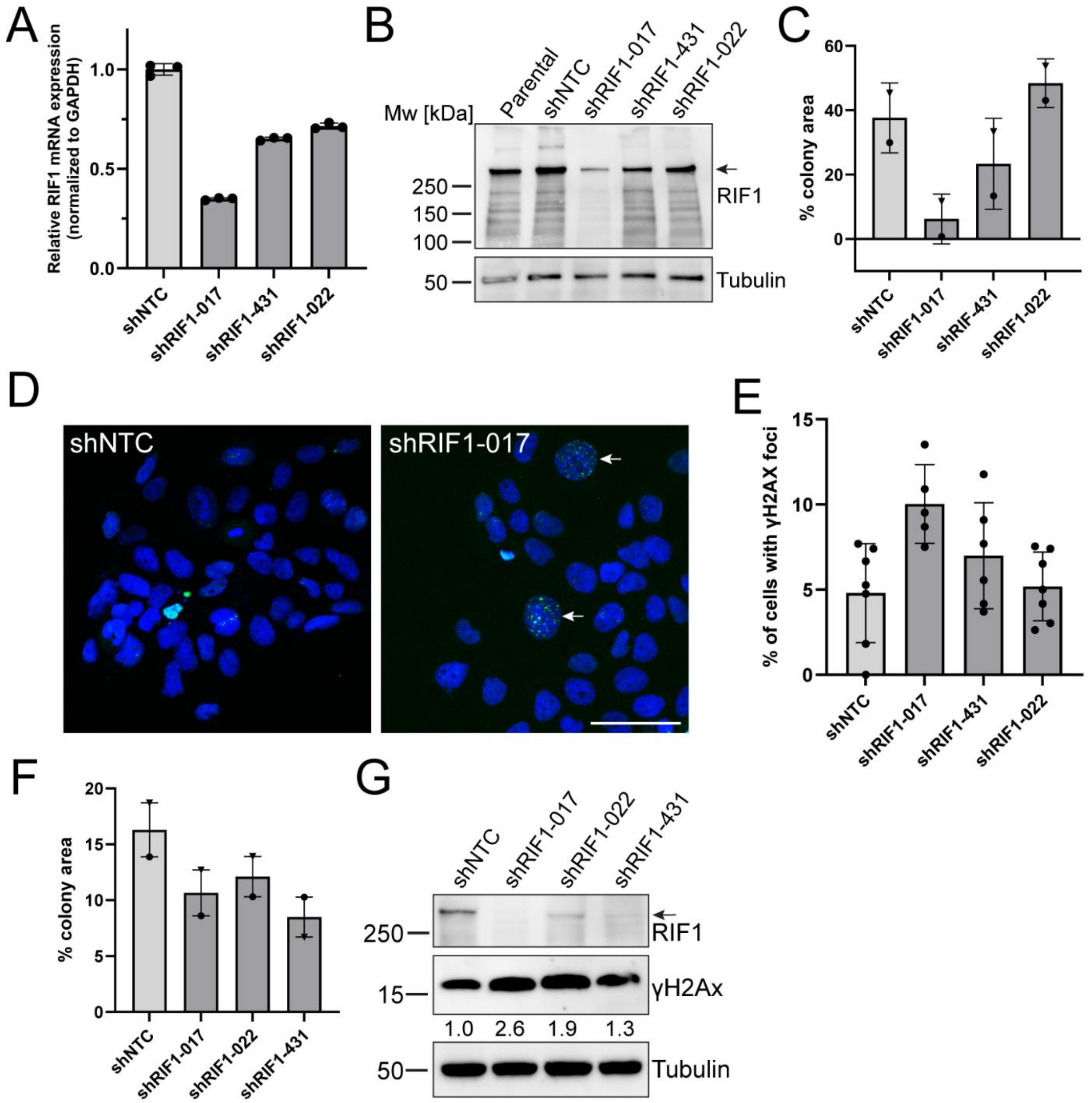


***Supplementary Figure 9. RIF1 depletion affects cell viability in HNSCC and LUSC cell lines.***

(**A**) qRT-PCR analysis of *RIF1* mRNA expression in SAS HNSCC cells expressing different shRIF1 or shNTC. Data shown are from n=1 experiment performed with three technical replicates. Individual data points: mean fold change in mRNA abundance (normalized to GAPDH) compared to siNTC in each experiment. Bars represent the means. (**B**) Immunoblot analysis of SAS cells expressing shNTC or different shRIF1, using the indicated antibodies. Tubulin was used as loading control. Arrow, position of full-length RIF1 band. (**C**) Colony formation assays were quantified by measuring the percentage of well area occupied by colonies. Bars show the means from n=2 independent experiments (performed with 3 technical replicates), individual data points (different shapes indicate different experiments) show the means from each experiment, error bars show SD. (**D**) Representative confocal fluorescence images (z-projections) of γH2AX (green) in shNTC and shRIF1 SAS cells. DNA was counterstained with Hoechst dye (blue). Arrows indicate nuclei with multiple γH2AX foci. Scale bar, 50 µm. (**E**) Quantification of percentage of cells with γH2AX foci in SAS cells expressing different shRIF1 or shNTC. Bars show the means from n=5–7 confocal images from n=1 experiment, individual data points show the cell percentages from each image, error bars show SD. (**F**) Colony formation assays were quantified by measuring the percentage of well area occupied by colonies. Bars show the means from n=2 independent experiments (performed with 3 technical replicates), individual data points (different shapes indicate different experiments) show the means from each experiment, error bars show SD. (**G**) Immunoblot analysis of HARA cells expressing shNTC or different shRIF1, using the indicated antibodies. Tubulin was used as loading control. Arrow, position of full-length RIF1 band. Numbers below lanes represent protein ratios relative to an arbitrary level of 1.0 set for shNTC.

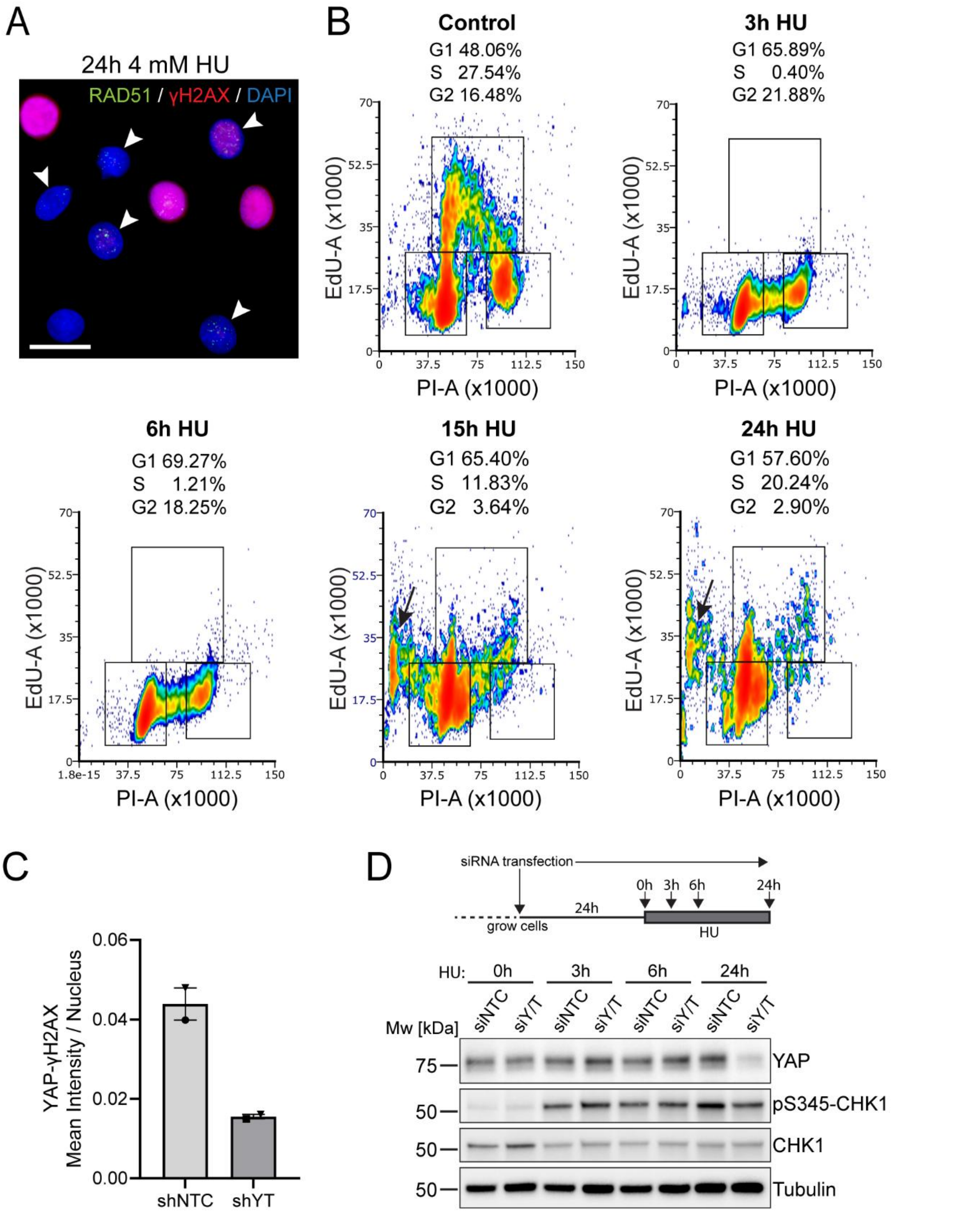


***Supplementary Figure 10. Prolonged hydroxyurea treatment induces replication fork collapse and DNA damage in SCC13 cells.***

(**A**) Representative microscopy image of RAD51 (green) and γH2AX (red) expression in SCC13 cells treated with 4 mM HU for 24 h. Nuclear DNA was counterstained with DAPI.

Arrowheads indicate cells with RAD51 foci. Scale bar, 25 µm. (**B**) Cell cycle profiles (PI labelling following a 30-min EdU pulse) of unsynchronised SCC13 cells at the indicated timepoints of 4 mM HU treatment. The percentages of cells in the different cell cycle phases are shown for each timepoint. Some cells appear to manage re-entering S phase during HU treatment, probably through firing of new origins. Arrows indicate sub-G1 fraction (apoptotic) cells. (**C**) PLA signal specificity control for experiment shown in Figure 8E. Datapoints (different shapes indicate different experiments) show mean nuclear PLA signal intensity in siNTC&shNTC cells and siYAP/TAZ&shYAP cells. Bars show means from n=2 independent experiments, error bars show SD. (**D**) Optimization of YAP/TAZ-targeted RNAi for the experiment shown in Figure 8F. 24 h after transfection with siNTC or siYAP/TAZ (siY/T) (both at 1 nM), SCC13 cells were treated with 2 mM HU for the indicated times. Immunoblot analysis confirmed activation of CHK1 upon HU treatment and YAP depletion by the experimental endpoint (48 h post-siRNA transfection). Tubulin was used as loading control.
