## Supplementary Materials and Methods for "YAP engages RIF1 to dampen replication stress-induced DNA damage in human squamous cell carcinoma"

### *Patient samples*

All human tissue work was performed in the lab of Dr Beate Lichtenberger at the Medical University of Vienna, Austria. Written informed patient consent was obtained before tissue collection in accordance with the Declaration of Helsinki. The study was approved by the Institutional Review Board under the ethical permits EK#1695/2021 and EK#1783/2020.

### *Cell culture*

All cell lines were routinely (quarterly test intervals) confirmed to be free of mycoplasma infection using PCR LookOut® Mycoplasma PCR Detection Kit (Merck).

Mouse 3T3 embryonic fibroblasts (subclone J2), originally established from random-bred Swiss mice [1], were used to provide optimal clonal growth support to normal and neoplastic human epidermal keratinocytes grown in co-culture [2]. Stocks were kindly provided by Prof Fiona Watt (Centre for Gene Therapy and Regenerative Medicine, King's College London, UK; EMBL, Heidelberg, Germany; not authenticated) or were purchased from Kerafast (Catalogue #EF3003). 3T3-J2 fibroblasts were routinely cultured in high-glucose DMEM (Merck) supplemented with 10% (v/v) bovine serum (Fisher Scientific), 100 IU/ml penicillin and 100 µg/ml streptomycin (Merck), and 2 mM L-glutamine (Merck) (complete fibroblast medium) at 37°C, 5% CO2. Fibroblast cultures were not allowed to grow to confluence and were not grown beyond passage 12 to prevent senescence or spontaneous transformation. To prepare feeder cell layers, 3T3-J2 fibroblasts were treated for 3 hours with 4 µg/ml mitomycin C (Merck) at 37°C, 5% CO2, to irreversibly stall the cell cycle [3].

Stock cultures of primary normal human keratinocytes (NHKs, strain km, male) were obtained from surgically discarded foreskin [4]. SCC13 cells are a cell line established from a recurring cutaneous SCC (cSCC) in facial epidermis of a female patient and were obtained from Dr James Rheinwald (Department of Dermatology, Harvard Skin Research Centre, USA), not authenticated [5]. To study the progression of human cSCC, we used four isogenic human cell lines that were isolated from skin biopsies from a immune-suppressed male patient at progressive disease stages: PM1 cells were derived from a pre-malignant dysplastic lesion on the patient’s head, MET1 was derived from a primary invasive cSCC growing on the patient’s right hand, MET2 was established from a rapid local recurrence after surgical removal of the primary tumour, and MET4 was established from an axillary lymph node metastasis that developed from the primary cSCC [6]. The progression series cell lines were purchased from Ximbio (PM1 #153571, MET1 #153539, MET2 #153569, MET4 #153570) and were last re-authenticated in Autumn of 2023. SCC-NR cells were isolated by Dr Nicholas Rabey at Addenbrooke’s Hospital (Cambridge, UK) from a portion of a surgically excised cutaneous SCC from the back of the hand of an elderly male patient and were used in all experiments at passages 3 or 4 [4]. All other cell lines were used at passages 2–8. NHKs and cSCC cells were grown in FAD medium (1 part high glucose DMEM GlutaMAX™ supplemented with pyruvate (Fisher Scientific) plus 3 parts Ham’s F12 GlutaMAX™ (Fisher Scientific), supplemented with $1.8 \times 10^{-4}$ M adenine (Merck), 10% (v/v) foetal bovine serum (Fisher Scientific, batch-tested for optimal clonal growth support), 5 µg/ml insulin (Merck), 0.5 µg/ml hydrocortisone (Merck), 10 ng/ml epidermal growth factor (Peprotech), 8.5 ng/ml cholera toxin (Merck), 2 nM T3 supplement (3,3’,5-triiodo-l-thyronine sodium salt) (Merck), and 100 IU/ml penicillin and 100 µg/ml streptomycin (Merck) on a feeder layer of mitotically inactivated 3T3-J2 cells, and grown at 37°C at 5% $CO_2$, as previously described [4].

HARA lung SCC cells (male) [7] and SAS HNSCC cells (female) were purchased from the National Institutes of Biomedical Innovation, Health, and Nutrition (NIBIOHN) JCRB Cell Bank, Japan (catalogue #JCRB1080.0 and #JCRB0260, respectively), and were re-authenticated in August 2024. HARA cells were cultured in complete RPMI medium (10% (v/v) fetal bovine serum, 100 IU/ml penicillin and 100 µg/ml streptomycin (Merck), and 2 mM L-glutamine (Merck). SAS cells were cultured in 45% (v/v) DMEM (Fisher Scientifc) and 45% (v/v) F-12 medium (Fisher Scientifc), 10% (v/v) fetal bovine serum, 100 IU/ml penicillin and 100 µg/ml streptomycin (Merck), and 2 mM L-glutamine (Merck).

***Generation of stable shRNA-expressing cell lines***

SCC13 cells (cultured on feeder cells), SAS cells, and HARA cells, cultured to ~70% confluence, were disaggregated in trypsin/EDTA and cells ($2.5 \times 10^5$) were seeded into 6-well cell culture plates (Corning™ Costar™, UK) and cultured for 24 h in complete growth medium. Medium was then changed to Opti-MEM (Fisher Scientific). To generate stable SCC13, SAS, or HARA cell lines expressing RIF1- or *CDKN1A*/p21-targeting shRNAs, SCC13, SAS, or HARA cells were infected with MISSION® lentiviral particles (Merck, the sequences of the shRNAs can be found in (Supplementary Table 3A) at a multiplicity of infection (MOI) of 2–3 in the presence of 5 µg/ml polybrene (Hexadimethrine bromide, Merck). For creating stable SCC13 cell lines expressing YAP- or TAZ-targeting shRNAs, the SMARTvector™ inducible lentiviral platform was used, which implements a microRNA-scaffolded Tet-On 3G bipartite induction system (Horizon Discovery-Dharmacon™, the sequences of the shRNAs can be found in Supplementary Table 3B). The lentiviral vector included a puromycin resistance element (PuroR) for selection post-infection, as well as a 2A self-cleaving peptide which allowed for PuroR and Tet-On 3G expression from a single mCMV promoter that was selected for negligible basal expression and strong activation when induced (data not shown). A doxycycline-inducible TRE3G promoter was driving expression of the shRNA as well as a fluorescent turbo-RFP reporter. After 24 h incubation with viral particles, medium was changed to FAD medium, and shRNA-expressing cells were selected for 7 days using puromycin (2.5 µg/ml, Merck) in co-culture with puromycin-resistant 3T3-J2 fibroblasts (kindly provided by Prof Fiona Watt, Centre for Gene Therapy and Regenerative Medicine, King's College London, UK; EMBL, Heidelberg, Germany). For experiments using the stable SCC13 cell lines with doxycycline inducible shRNAs, cells were incubated with FAD medium supplemented with 5 µg/ml doxycycline (Tocris) for 48 hours before trypsinisation and seeding for downstream experiments.

***Generation of transgenic cell lines overexpressing inducible YAP constructs***

Transgenic SCC cell lines were generated in the lab of Caterina Missero (University of Naples, Italy). To express wild-type YAP and hyperactive YAP mutants (S127A and 5SA), a doxycycline-inducible system was used. To prepare the lentiviral particles, HEK293T cells were co-transfected with second-generation viral packaging plasmids (psPAX2 and pMD2.G), as well as FUW-tetO-wtYAP (Addgene #84009), FUW-tetO-YAP1-S127A [8], FUW-tetO-YAP1-5SA [8], FUdeltaGW-rtTA (Addgene #19780, encoding the repressor), or the FUW-tetO-MCS control plasmid (Addgene #84008), using Lipofectamine 2000 (Invitrogen). The plasmids carrying the cassettes for expressing the YAP mutants were kindly provided by Prof Stefano Piccolo (University of Padua, Italy). Briefly, $3 \times 10^6$ HEK293T cells were seeded in a 6 cm cell culture dish and, the next day, transfected with psPAX2 and pMD2.G vectors together with the plasmid of interest at a ratio of 3 µg: 1 µg: 2 µg and 15 µl of Lipofectamine 2000 (Fisher Scientific). 24 hours later, the medium was replaced and virus supernatant collection was performed after 48 hours and agaon after 72 hours. Viral supernatants were pooled togheter and filtered through 0.45 µm filters. 1 x $10^6$ SCC13 cells were seeded in a 6 cm tissue culture plate and co-infected the following day for 3 hours with 500 µl of rtTA and 500 µl of either wt- or mutant YAP, or control (MCS) viral supernatant aliquots. Transgene expression was induced by 5 µg/ml doxycycline (Tocris). Transgenic cell lines were cultured in Keratinocyte SFM (Thermo Fisher Scientific).

***siRNA transfection***

24 hours after seeding, cells were transfected with 1-20 nM of ON-TARGET plus SMARTpool siRNAs (Horizon Discovery, Supplementary Table 3C) using INTERFERin® transfection reagent (Polyplus, VWR) in Opti-MEM (Fisher Scientific) for 5 hours, then washed with PBS and grown in FAD medium for up to 72 hours.

***Pharmacological inhibitors***

GNE-7883, MGH-CP1, and K-975 were obtained from MedChemExpress. Hydroxyurea was purchased from Fisher Scientific and was dissolved in sterile Milli-Q water; all other inhibitors were dissolved in sterile DMSO with sonication.

***Colony formation assay***

1 x $10^3$ cells per well were seeded in triplicate technical replicates into 6-well cell culture plates (Corning™ Costar™) without feeder cells and were cultured for 10-14 days with

frequent (every 2–3 days) media changes, then fixed in 4% PFA for 30 minutes at room temperature. In case of treatment with hydroxyurea, different drug concentrations were added 24 hours after seeding and fresh drug was added with each medium change. Colonies were stained using crystal violet solution (2% (w/v) crystal violet (Merck) dissolved in 20% (v/v) methanol) for one hour at room temperature and then washed extensively in deionised water and air-dried. Colonies were either imaged using a transilluminator and cell phone camera or were scanned using a flat-bed scanner. Quantification of culture area covered by colonies were performed using ImageJ.

***Cell viability assay***

MET1, MET2, and MET4 cells were seeded at a density of 1 x $10^3$ cells per well into 96 well plates (Corning™ Costar™), while SCC13 were seeded at a density of 3 x $10^3$ cells per well. After 24 hours, cells were treated with different TEAD inhibitors at different concentrations in triplicate wells for 96 hours. DMSO diluted in complete FAD medium was used as vehicle control. Cells were incubated at 37°C in 5% $CO_2$ for 96 hours, and their viabilities were then measured using the EZMTT cell viability assay (Merck). Briefly, the 50× EZMTT benzenesulfonate sodium salt was diluted 1:100 in complete FAD medium, the medium was aspirated from the wells, and the diluted EZMTT reagent was added. Cells were then incubated at 37°C for 1.5–2 hours. Absorbance at 450 nm was measured using a FLUOstar Omega plate reader (BMG LABTECH) and Omega software (version 1.20).

***Cell proliferation assays***

For CyQUANT™ assay, cells were seeded at a density of 5 x $10^3$ cells per well in black, clear-bottom 96-well plates (Greiner). After 24 hours, cells were treated with different concentrations of pharmacological inhibitors for 120 hours. Cell proliferation was assessed using the CyQUANT™ Direct Cell Proliferation Assay (Thermo Fisher Scientific). The Detection Reagent was prepared in PBS by diluting CyQUANT™ Direct Nucleic Acid Stain 1:250 and CyQUANT™ Direct Background Suppressor 1:50. Cells were incubated with 80 µl of the Detection Reagent (per well) for 60 minutes, at 37°C. Fluorescence intensity was measured using excitation/emission of 508/527 nm using the PHERAstar® FMX plate reader (BMG LABTECH).

Cell proliferation kinetics were also measured using the IncuCyte® SX5 Live-Cell Analysis System (Sartorius) within an incubator at 37°C with 5% $CO_2$. SCC13 cells stably expressing different doxycycline-inducible YAP transgenes or empty vector were seeded at a density of $3 \times 10^3$ into 6-well plates (Sarsted) 24 hours before imaging. Transgene expression was induced with 5 ug/ml doxycycline and cells were imaged using the 'Standard' module with a 10X objective (Nikon CFI Plan Fluor 10X 0.3 NA). Confluency was calculated using the AI Confluency model, with a minimum cell area set to 600 $um^2$. Confluency was normalised to the first scan for all conditions.

***3D spheroid growth assay***

SCC13 cells were seeded at a density of $2 \times 10^3$ cells per well in ultralow-adherence 96-well U-bottom plates (Corning Costar, 7007). Drug treatments were started after 48 hours, when cells had formed single spheroids (10 replicates per condition). Spheroid growth was observed for up to 15 days using the IncuCyte® SX5 Live-Cell Analysis System (Sartorius) within an incubator at 37°C with 5% $CO_2$. Individual spheroids were imaged at 10x magnification using phase + brightfield image on the day of treatment (day 0) and subsequently every 3-4 days. IncuCyte 2024A software was used analyse single spheroids with eccentricity < 0.7 and minimum area of 2 x $10^4$ $\mu m^2$. The largest brightfield object area ($\mu m^2$) for each spheroid was calculated, using brightfield channel masks surrounding the spheroid. CellTiter-Glo® 3D cell viability assay (Promega, G9681) was used to measure cell viability at the experimental end point. 100 µl of the CellTiter-Glo reagent was added to each well, and plates were covered in foil and incubated for 30 minutes at room temperature to allow for spheroid lysis. Luminescence was recorded using the PHERAstar® FMX plate reader (BMG LABTECH).

***Cell synchronisation by double thymidine block***

Cells were seeded in FAD medium and treated with 2.5 mM thymidine (Merck, UK) for 24 hours at 37°C, 5% $CO_2$. Cells were then washed with FAD medium (without thymidine), fresh FAD medium (without thymidine) was added, and cells were incubated for 15 hours before the medium was supplemented again with 2.5 mM thymidine for a further 24 hours. Following thymidine washout, cells were grown in FAD medium.

***Cell cycle analysis by flow cytometry***

Prior to trypsinization, Click-iT™ EdU (Fisher Scientific, UK) was added to the cell culture medium to a final concentration of 10 µM, and cells were incubated for 30 minutes at 37°C. Trypsinized cells were then centrifuged at 500 x g for 3 minutes (RT), the supernatant was removed, and the pellet resuspended in 3 ml PBS, followed by another centrifugation at 500 x g (RT) for 5 minutes. Pelleted cells were next resuspended in 100 µl 0.9% (w/v) NaCl and added dropwise to 1.8 ml ice cold 70% (v/v) ethanol for fixation and stored at -20°C for a minimum of 2 hours before being centrifuged again for 5 minutes at 400 x g (4°C). The cell pellet was then resuspended in 0.5 ml PBS and transferred to a clean tube, and cells were centrifuged for 5 minutes at 400 x g (4°C). This cell pellet was then resuspended in 100 µl Click-iT™ cell reaction buffer (Fisher Scientific) and incubated at RT in the dark for 30 minutes. EdU-labelled cells were then centrifuged for 5 minutes at 400 x g (4°C), cell pellets were resuspended in 500 µl 1% (w/v) BSA in PBS and centrifuged for 5 minutes at 400 x g (4°C), then resuspended in 500 µl PBS. RNase (Merck) was added to a final concentration of 0.5 µg/ml and the sample was incubated on a heat block at 37°C for 15 minutes. Propidium iodide (Merck) was added to a final concentration of 20 µg/ml and the sample was incubated on ice for 20 minutes in the dark before a minimum of 20,000 cells were analysed on a FACS Canto™ cell sorter (BBD Biosciences). Data were analysed using FCS Express™, De Novo Software (version 7.14.0020).

***Rapid immunoprecipitation mass spectrometry of endogenous proteins (RIME)***

Low passage cells were cultured on a mitotically inactivated fibroblast feeder layer to maintain the stem cell state of NHKs characterised by predominantly nuclear localisation of YAP [4, 9]. To this end, NHK and SCC13 cells were seeded at clonal density (5 x $10^5$ cells) into 15 cm dishes containing a mitotically inhibited 3T3-J2 fibroblast feeder layer and cultured for 4 days. This enabled the formation of small colonies with predominantly nuclear YAP/TAZ expression (Supplementary Fig. 5B). Two independent cultures (=independent experiments, starting from separate cell stocks) were grown for each cell type. Following removal of the feeder layer on day 4 of culture, cells were grown in complete FAD medium (that had been conditioned by mitotically inhibited 3T3-J2 fibroblast cells) for a further 24 hours. Cells were fixed with 1% (v/v) methanol-free formaldehyde for 8 min at RT and the fixation reaction was subsequently rapidly quenched with 0.125 M glycine for 5 min. Fixed cells were gently scraped into ice-cold washing buffer (0.5% (v/v) IGEPAL CA-630, 100 µM PMSF, in PBS, pH

7.4) and were washed three times by centrifugation (800 x g, 10 minutes, 4°C) and resuspension. The final cell pellets (containing between 2 x $10^7$ and 7 x $10^7$ cells) were stored at -80°C and shipped to Active Motif (USA) on dry ice.

RIME was performed by Active Motif (USA). Chromatin was isolated from formaldehyde-fixed cell pellets by the addition of lysis buffer (custom, proprietary to Active Motif), followed by cell disruption with a Dounce homogenizer (15 ml volume with tight-fitting pestle) to release nuclei. Nuclei were sonicated on ice in ChIP buffer (custom, proprietary to Active Motif) using an EpiShear Probe Sonicator (68% pulsed sonication: 30sec on/30sec off, 10 minutes total sonication time) to lyse the cells and shear the DNA to an average fragment length of 300-500 bp. Genomic DNA for chromatin quantification was prepared by treating aliquots of chromatin with RNase (Merck), proteinase K (Merck) and heat treatment (65°C overnight) for de-crosslinking, followed by ethanol precipitation. Pellets were resuspended in 1/5 volume Tris-EDTA buffer, and the DNA concentration was quantified on a NanoDrop 2000 (Fisher Scientific, USA) spectrophotometer. An aliquot of sheared chromatin (150 µg) was precleared with protein G agarose beads (Invitrogen). For each chromatin preparation, YAP RIME was performed in technical replicates. Control RIME was performed using isotype-matched IgG, and for each independent experiment, the two replicate IgG control immunoprecipitations were pooled for mass spectrometry analysis. YAP chromatin immunoprecipitation was performed using 15 µg of purified rabbit polyclonal YAP-specific antibody (US Biologicals, Y1200-01D) or isotype-matched rabbit IgG (control) and protein G magnetic beads (Active Motif). Immunoprecipitated chromatin-protein complexes were washed extensively with RIPA buffer (1x, Active Motif) and then with 100 mM ammonium bicarbonate (Merck). On-bead proteolytic digestion was performed using trypsin (Worthington Biochemical, USA; 10 ng/µl). Protein digests were separated from the beads using a magnetic rack, and digested peptide mixtures were desalted by solid-phase separation using a C18 spin column (Harvard Apparatus, USA). The eluted peptides were vacuum dried using a speedvac.

Digested peptides were reconstituted in 5% (v/v) acetonitrile and 0.1% (v/v) trifluoroacetic acid and analysed by LC-MS/MS on a Thermo Scientific Q Exactive Orbitrap Mass spectrometer linked to Dionex Ultimate 3000 HPLC (Thermo Scientific) and a nanospray Flex$^{TM}$ ion source. The digested and desalted peptides were loaded directly onto the separation column (Waters BEH C18, 75 µm x 100 mm, 130Å 1.7u particle size). Peptides were eluted using a 120-minute gradient with a flow rate of 323 nl/min. The gradient was formed using increasing/decreasing concentrations of buffer B (0.1% formic acid in acetonitrile); (1) increase to 5% (v/v) B over 12 min, (1) gradual increase to 25% (v/v) B until the 90 min mark, (3) gradual increase to 95% (v/v) B for 5 min, (4) 95% (v/v) B for another 4 min, (5) gradual decrease to 5% (v/v) B over 20 min. An MS survey scan was obtained for the m/z range 340-1600; MS/MS spectra were acquired using a top 15 method, where the top 15 ions in the MS spectra were subjected to HCD (High Energy Collisional Dissociation). An isolation mass window of 1.6 m/z was set for precursor ion selection, and normalized collision energy of 27% was used for fragmentation. A 20-second duration was used for the dynamic exclusion.

Tandem mass spectra were extracted and analysed by PEAKS Studio (version X+, Bioinformatics Solutions Inc., Canada). Charge state deconvolution and de-isotoping were not performed. Reference databases consisted of the UniProtKB/Swiss-Prot database (version 180508; 71,771 curated entries) and the cRAP database of common laboratory contaminants (www.thegpm.org/crap; 114 entries). Databases were searched with a mass tolerance of 0.02 Da for fragment ions in MS/MS mode, and a parent ion tolerance of 10 PPM in MS mode. Post-translational protein modifications allowed were methionine oxidation, asparagine and glutamine deamidation, which were searched as variable modifications.

PEAKS Studio software built-in decoy sequencing and FDR determination was used to validate MS/MS-based peptides. PEAKS Studio uses the "Decoy Fused Method" when calculating FDR where the default cut-off is set to -10logP > 20 to ensure high-quality peptide spectrum matches and to increase confidence in the proteins list obtained. Protein identifications were accepted if they could pass the -10logP of > 20 and contained at least 1 identified unique peptide. FDR for peptide spectrum matches was set at <1%. Proteins that contained similar peptides and could not be differentiated based on MS/MS analysis alone were grouped to satisfy the principles of parsimony. Proteins sharing significant peptide evidence were grouped into protein groups.

***Semiquantitative analysis of RIME proteomics data by spectral counting***

The following filtering criteria were applied to the protein data obtained in PEAKS studio for each cell type (Supplementary Table 2). (1) Identified proteins with <2 unique peptides were removed. (2) We then filtered out proteins with a spectral count average >2 in the IgG immunoprecipitation controls for experiments 1 and 2 combined, and proteins with a spectral count average of <4 in any of the two YAP RIME experiments. (3) Proteins matched to commonly identified IgG contaminant proteins (data downloaded from the contaminant repository for affinity purification (CRAPome, Crapome.org, accessed November 2019)) [10] and to mitochondrial proteins were excluded from further analysis. (4) Proteins displaying a fold change <4 of mean spectral counts in YAP immunoprecipitations vs spectral counts in the respective IgG control immunoprecipitation in any of the two independent experiments were removed to eliminate low confidence hits that were not considerably more represented in the YAP immunoprecipitation than the IgG immunoprecipitation. Using the 'Compartments' database (https://compartments.jensenlab.org/Search), most identified proteins had a high confidence score (≥4) for nuclear localization (Supplementary Table 2). Moreover, significantly enriched gene ontology terms for molecular functions (GO MF) and biological processes (GO BP) in both protein interactomes were all related to processes occurring in the nucleus (Fig. 3B, Supplementary Table 2). High confidence YAP-interacting proteins were further screened for potential physiological significance by filtering against lists of candidate genes positively and negatively regulating clonal growth of NHK and SCC-13 cells (Walko et al., 2017). Interaction networks were created using Cytoscape version 3.9.1 and using the Cytoscape STRING app version 1.7.1 and manually curated into functional groups using input from Gene Ontology Enrichment Analysis performed using gprofiler (https://biit.cs.ut.ee/gprofiler/gost) functional profiling tool with electronic GO annotations unselected (Reimand et al., 2019) and visualised using EnrichmentMap (Cytoscape app version: 3.3.4) in Cytoscape (version:3.9.1).

***Immunofluorescence microscopy***

Cells were seeded onto acid-washed and UV-sterilized glass coverslips coated with rat tail collagen I (10 µg/ml; Fisher Scientific) and fixed using 4% (v/v) paraformaldehyde (PFA) for 30 minutes at RT. Immunostaining was performed in a PBS-humidified chamber. Cells were permeabilised with 0.1% (v/v) Triton-X100 (Merck) for 25 minutes then blocked in 10% (v/v) BSA (Merck) + 0.1% (v/v) Triton-X100. Primary antibodies (Supplementary Table 4) were diluted in antibody diluent buffer (2% (v/v) BSA + 0.1% (v/v) Triton-X100) and incubated overnight at 4°C. Coverslips were washed thoroughly with PBS, secondary antibodies (Supplementary Table 4), diluted in antibody diluent, were added, and the samples incubated for 1 hour at RT in the dark, then washed multiple times with PBS and once with MilliQ water. For RAD51/γH2AX immunostainings, cells growing on glass coverslips were fixed with ice cold methanol for at least 1 hour at -20°C. 2% (w/v) BSA, 10% (w/v) milk powder, and 10% (v/v) goat's serum in PBS + 0.2% (v/v) Triton-X-100 was used for blocking in this case. Coverslips were mounted onto glass microscope slides using ProLong Gold™ anti-fade mounting medium with DAPI (Fisher Scientific) or alternatively, incubated with 2 µg/ml Hoechst 33342 (Fisher Scientific) in PBS for 5 minutes, washed once with PBS, and mounted with ProLong Gold™ anti-fade glass mounting medium without DAPI (Fisher Scientific).

For confocal microscopy, images were acquired using a ZEISS LSM 880 laser scanning confocal microscope using 40x/1.3 Oil DIC UV-IR M27 or 63x/1.4 Oil DIC UV-IR M27 objectives at 16 Bit depth, 0.5 µm optical Sections for Z-stacks, and 2048x2048 pixel resolution. Images were processed using Fiji/ImageJ (ImageJ; https://fiji.sc/ ) [11] or ZEN blue (Zeiss) with maximal orthogonal projection of Z-stacks. Image quantification was performed using CellProfiler (version 4.0.7) [12]. Minimal processing (changes in brightness and contrast) was applied equally across entire images and applied equally to controls.

Images of RAD51/γH2AX double immunostainings were acquired on a Leica DM6 microscope using Leica HC PL APO 40×/0.85 CORR CS and Leica HC PL APO 100×/1.40 OIL CS objectives, and at 24-bit depth and 2048x2048 pixel resolution. Acquisition settings remained consistent across all conditions. Minimal processing (changes in brightness and contrast) was applied equally across entire images and applied equally to controls.

Immunofluorescence images in Supplementary Figure 1B were obtained on a ZOE® Fluorescent Cell Imager (Bio Rad) using standard setup (20x objective) and digital zoom.

Minimal processing (changes in brightness and contrast) was applied equally across entire images and applied equally to controls.

***Quantification of RIF1 expression in human tissues***

Immunofluorescence staining was performed on 4 µm human FFPE sections according to standard protocols. Antigen retrieval was conducted in Dako Target Retrieval Solution, pH 9.0 (Agilent Technologies), and 3% (w/v) BSA in PBS containing 0.05% (v/v) Tween 20 (PBS-T) was used for blocking. Primary antibodies against Keratin 14 (chicken pAb (Poly9060), BioLegend Cat# 906004, 1:250) and RIF1 (rabbit mAb (D2F2M), Cell Signalling Technology Cat# 95558, 1:50) were diluted in 1% (w/v) BSA in PBS-T and incubated overnight. Fluorescently labelled donkey anti-chicken (IgY (H+L) highly cross adsorbed, Alexa Fluor™ 594, Fisher Scientific, Cat# A78951, 1:500,) and donkey anti-rabbit (IgG (H+L) highly cross-adsorbed secondary antibody, Alexa Fluor™ 488, Fisher Scientific, Cat# A-21206, 1:500) secondary antibodies were incubated for 90 minutes at ambient temperature. DAPI was used for nuclear counter-staining. Images were captured using the Vectra Polaris™ imaging system, and image analysis was conducted with the HALO® image analysis platform v3.6.4134. The HighPlex FL v4.2.14 plugin was employed for image analysis, with signal intensity thresholds set for the Opal620 (AF594) channel to differentiate positive from negative cells. The DenseNet V2 (Halo AI) classifier was trained and applied to differentiate between tumour and stroma regions. Depending on the size of the tissue, between 2-13 regions of interest with an area of 1 $mm^2$, representing the tumour were analyzed.

Additional images of human squamous cell carcinoma samples were obtained by datamining the www.proteinatlas.org database [13, 14].

***In-situ proximity ligation assay***

Cells were fixed and permeabilised as described above. Coverslips were blocked in Proximity Ligation Assay (PLA) (Duolink®, Sigma) blocking buffer, and primary antibodies (Table S4), diluted in Duolink® antibody diluent, were added overnight. Proximity ligation assay reactions were performed as per the manufacturer's instructions. Coverslips were mounted onto glass slides using Duolink® PLA mounting medium with DAPI and sealed with clear nail polish. Images were acquired on a Zeiss LSM-880 confocal microscope using a 40x/1.3 Oil DIC UV-IR M27 objective. All images belonging to the same experimental repeat were acquired in one imaging session at 16-Bit depth using a Z-stack (0.5 µm slice interval) and a 2048x2048 pixels dimension. Images were analysed on ZEN blue (version 3.3.89.0000) and processed using a maximal frontal orthogonal projection including all Z-stack slices. Background fluorescence was removed using the same parameters for each image belonging to the same image set (= experimental repeat). Image quantifications were performed using CellProfiler version 4.0.7. Nuclei were defined as parent objects using the 'IdentifyPrimaryObjects' module based on DAPI fluorescence signal and intensity of PLA signal per nucleus was quantified within the defined nuclear area using the 'MeasureIntensity' module. To account for differences in YAP-RIF1 and YAP-TEAD PLA fluorescence signal intensities between different experimental repeats, mean nuclear PLA signal fluorescence intensity in shYAP/TAZ cells was normalized to that in shNTC cells [15]. A one sample two-tailed t test was then used to test for statistical significance [15]. Statistical significance between the mean PLA signal intensities in the shNTC and shY/T cell populations (assessed by Mann-Whitney test) was high ($p<0.0001$) for each individual experiment (not shown).

***EU incorporation assay***

EU incorporation assay was performed using the Click-iT RNA Alexa Fluor 488 Imaging Kit (Invitrogen) according to the manufacturer's instructions. Cells were incubated with 1 mM EU for 1 h, fixed with 4% (v/v) PFA for 15 min at room temperature, permeabilized with 0.5% (v/v) Triton X-100 for 15 min and Click-iT reaction was performed. DNA was counterstained with DAPI, and images were acquired were obtained on a ZOE® Fluorescent Cell Imager (Bio Rad) using standard setup (20x objective) and digital zoom. Fiji/ImageJ (ImageJ; https://fiji.sc/ ) [11] was used to generate nuclear masks based on DAPI staining and mean AlexaFluor 488 fluorescence intensities were quantified per nucleus.

***Preparation of protein extracts for immunoblotting***

RIPA buffer (1x) (Cell Signalling Technologies) or Cell Lysis buffer (1x; Cell Signalling Technologies, #9803; for co-immunoprecipitation experiments), supplemented with protease inhibitors (cOmplete™ ULTRA, EDTA-free, Merck) and phosphatase inhibitors (PhosStop, Merck, UK), was added to cell culture plates and cells were lysed on ice for 15 minutes. Cell remnants were then scraped into the lysis buffer; the lysate was transferred into a tube and sonicated in an ice bath using an MSE Soniprep 150 plus sonicator at 12 Amp for 15 minutes with shaking. Lysates were centrifuged at 16,000 x g for 10 minutes (4°C). For co-immunoprecipitation experiments, sample sonication was not performed. The supernatant was collected, and BCA assay was performed to measure protein concentration using Pierce BCA Protein Assay Kit (Thermo Fisher).

***Preparation of chromatin-associated protein fractions***

Chromatin fractions were prepared as described [16]. Briefly, cytoskeleton buffer (300 mM sucrose, 3 mM $MgCl_2$, 100 mM NaCl, 0.5% (v/v) Triton X-100, 10 mM HEPES (pH 7.4)) supplemented with protease inhibitors (cOmplete™ ULTRA Tablets, EDTA-free, Merck) and phosphatase inhibitors (PhosStop, Merck) was added to cells growing in cell culture dishes, and these were incubated on ice for 5 minutes. Cells were then scraped into the buffer, and the extract was transferred to a microcentrifuge tube and incubated on ice for further 10 minutes. Samples were then centrifuged at 5,000 x g for 5 minutes at 4°C, and the supernatant was removed and discarded. Chromatin pellets were washed twice in cytoskeleton buffer without Triton-X100 and resuspended in 50 µl RIPA buffer (Cell Signalling Technologies) containing protease and phosphatase inhibitors and incubated on ice for 5 minutes. 50 units Benzonase® Nuclease (Merck) and $MgCl_2$ (Merck, final concentration: 3 mM) were added, and the chromatin fraction was incubated on ice for 20 minutes until it had visibly reduced viscosity. The chromatin fraction was then centrifuged at 16,000 x g for 10 minutes at 4°C, and the supernatant was recovered for immunoblotting analysis.

***Co-immunoprecipitation from nuclei-enriched fractions***

Cells were grown to ~70% confluency and were washed twice with ice-cold PBS before incubation on ice for 15 minutes with nuclear extraction buffer (50 mM KCl, 2 mM $MgCl_2$, 1 mM EDTA, 1 mM DTT, 0.1% (v/v) IGEPAL CA-630, 10 mM HEPES (pH 7.4)), supplemented with protease inhibitors (cOmplete™ ULTRA, EDTA-free, Merck) and phosphatase inhibitors (PhosStop, Merck). Cell remnants were then scraped into the buffer, the suspension was collected into pre-cooled microcentrifuge tubes, vortexed for 10 seconds, and centrifuged at 750 x g for 5 minutes (4°C) to pellet nuclei. The cytoplasmic fraction was removed and the nuclear pellet was washed once in ice-cold PBS, resuspended in ice-cold nuclear lysis buffer

(450 mM NaCl, 0.1 mM EDTA, 3 mM $MgCl_2$, 0.1% (v/v) Triton X-100, 1% (v/v) IGEPAL CA-630, 1 mM DTT, 20 mM HEPES (pH 7.4)) supplemented with protease inhibitors (cOmplete™ ULTRA, EDTA-free, Merck) and phosphatase inhibitors (PhosStop, Merck), transferred to a clean pre-cooled micro-centrifuge tube, and incubated on ice for 15 minutes. Chromatin was then sheared either by passaging 10 times through a 27G needle attached to a 1 ml syringe or by the addition of Benzonase® Nuclease (Merck). Needle-sheared nuclear fractions were centrifuged at 13,000 x g for 10 minutes (4°C) and the supernatant was diluted (1:3) to a final concentration of 150 mM NaCl using nuclear lysis buffer without NaCl. For Benzonase®-treatment of nuclear fractions, chromatin suspension in nuclear lysis buffer was similarly adjusted to 150 mM NaCl using nuclear lysis buffer. Benzonase® was then added, and the chromatin suspension was incubated on ice for 15 minutes with intermittent vortexing. EDTA was added to a 1 mM final concentration, and samples were centrifuged at 13,000 x g for 10 minutes (4°C).

BCA assay was performed to measure protein concentration, and agarose gel electrophoresis of DNA was used to validate the Benzonase digest. To this end, a 1% (w/v) agarose (Merck) gel was prepared using 1x Tris-acetate-EDTA buffer (Fisher Scientific) and GelRed® Nucleic Acid Stain (Merck). 10 µl of sample was incubated with 1 µl proteinase K for 1 hour at 65C and then diluted with TriTrack DNA loading dye (Fisher Scientific) and run for 1-2 hours at 100V in parallel with GeneRuler1 kb Plus DNA Ladder (Thermo Fisher). Gels were imaged using a Syngene Gel Doc system.

Nuclear protein samples (containing 2 mg/ml protein) were incubated with 0.07 ug anti-YAP (D8H1X, Lot 4, Cell Signalling Technology) antibody or 0.07 ug of isotype-matched IgG control antibody (Table S4) and DTT (1 mM) in micro-centrifuge tubes overnight at 4°C with constant rotation. Protein G Dynabeads™ (Thermo Fisher) were washed four times in PBS supplemented with 0.05% (v/v) Tween-20 (PBS-T, IP wash buffer) using a DynaMag™ magnetic rack (Thermo Fisher), transferred to a new tube and washed once in IP wash buffer. Protein G Dynabeads were then resuspended in 50 µl IP-buffer and added to IP reactions, which were then incubated with the beads for 4 hours at 4°C with constant rotation. To pellet the beads, samples were then placed on the DynaMag™ magnetic rack, the supernatant was removed, bead-bound protein complexes were washed 3 times in PBS-T, transferred to a fresh micro-centrifuge tube, resuspended in 20 µl 50 mM glycine (pH 2.8) and 10 µl 3x Laemmli buffer (BioRad, UK) + β-mercaptoethanol (elution buffer), and incubated for 10 minutes at 95°C. Beads were removed using the DynaMag™ magnetic rack and the eluted proteins were analysed by SDS-PAGE (alongside 10 ug (1%) of input lysate) and immunoblotting.

***Immunoblotting analysis***

Protein samples were denatured and reduced in Laemmli buffer (BioRad) containing 10% (v/v) β-mercaptoethanol for 5 minutes at 95°C and loaded onto precast 4-20% or 4-15% polyacrylamide mini-PROTEAN Tris-Glycine eXtended (TGX) gels (BioRad). Proteins were transferred onto either 0.2 µm pore-size Nitrocellulose membranes or methanol-activated 0.45 µm pore-size PVDF membranes (BioRad), using either the Trans-Blot Turbo Transfer System (BioRad) or tank (wet) transfer depending on protein molecular mass. Large proteins were transferred by wet transfer in wet transfer buffer (3.05 g/l Tris-base, 14.4 g/l glycine) either overnight at 25V (4°C) or for 60 minutes with extensive cooling in wet transfer buffer containing 20% (v/v) methanol at 300 mA. Membranes were blocked in 5% (w/v) BSA dissolved in Tris-buffered saline (TBS) containing 0.1% (v/v) Tween 20 (TBS-T) (=blocking buffer) for 1 hour at RT and primary antibodies (Supplementary Table 4), diluted in blocking buffer, were incubated with the membranes overnight at 4°C with agitation. Membranes were washed three times for 10 minutes with TBS-T and then incubated with secondary horseradish peroxidase (HRP)-conjugated antibodies. Clarity western ECL substrate reagent (BioRad) was used to image immunoblots using a Fusion Solo chemiluminescence imager (Vilber Lourmat, UK). Where possible, over-saturated images were avoided and exposures within the dynamic range were selected. Band intensities were quantified based on pixel volume of the bands using Fusion software (Vilber Lourmat, UK) or Image Lab software (Standard edition, Bio Rad).

***Sample preparation for DNAscent***

SCC13 cells (growing in the presence of mitotically inactivated 3T3-J2 feeder cells) were incubated with 50 µM EdU (Fisher Scientific) for 5 minutes, 50 µM BrdU (Abcam) for 10 minutes, followed by thymidine (Fisher Scientific) for 20 minutes, with washes performed between each incubation using warm (37°C) FAD medium. After feeder removal, SCC13 cells were collected by scraping into ice-cold PBS, pelleted, washed at 4°C and frozen at -80 degrees.

High molecular weight (HMW) genomic DNA was extracted using Puregene Cell Kit (158043, Qiagen) according to the manufacturer's protocol. The concentration, quality, and length of genomic DNA were measured via Qubit and genomic DNA Tapestation analysis (5067-5365, Agilent). Sequencing libraries were prepared using either Ligation Sequencing Kit V14 (SQK-LSK114, Oxford Nanopore Technologies) or Ultra-Long DNA Sequencing Kit V14 (SQK-ULK114, Oxford Nanopore Technologies) following the manufacturer's protocol. Sequencing was performed by Genomics Birmingham (University of Birmingham) by loading prepared libraries onto PromethION flow cells R10.4.1 (FLO-PRO114M, ONT). Sequencing runs lasted 72 hours, with fresh library loading performed every 24 hours (for SQK-ULK114 libraries) by pausing the runs and washing flow cells with Flow Cell Wash Kit (EXP-WSH004, ONT) according to the manufacturer's guidelines.

***RT-qPCR***

mRNA was converted to cDNA (including the removal of genomic DNA) using the QuantiTect Reverse Transcription Kit (Qiagen) as per the manufacturers' instructions. For the Comparative $C_T$ method, qPCR reactions were set up using 50 ng cDNA, 10 µl 2x TaqMan™ Fast Advanced Master Mix (Applied Biosystems™, Fisher Scientific), 1 µl 20x TaqMan™ Gene Expression Assay (FAM) (Fisher Scientific) (Table S3D) and nuclease-free water. Reactions were performed in technical triplicates in MicroAmp® Fast 96-Well reaction plates (Applied Biosystems™, Fisher Scientific) and run on a StepOnePlus™ Real-Time PCR System (Fisher Scientific) for 40 cycles; 20 seconds at 95°C, 1 second at 95°C, 20 seconds at 60°C. Relative gene expression was analysed by the $2^{(-\Delta\Delta C_T)}$ method. GAPDH was used as housekeeping gene for normalization.

***RNA preparation and Illumina sequencing***

RNA was extracted from cells using the RNeasy Plus Mini kit (Qiagen) and the QIAshredder shredding system (Qiagen) as per the manufacturer's instructions. RNA yield and purity were analysed using the Quant-iT™ Qubit RNA BR Assay Kit (Thermo Fisher) in Qubit™ Assay Tubes (Thermo Fisher) on a Qubit 4 Fluorometer (Thermo Fisher). RNA sequencing was outsourced to Novogene Ltd. (UK). Messenger RNA was purified from total RNA by poly-A enrichment method using poly-T oligo-attached magnetic beads. RNA library preparation was performed using the Novogene in-house kit NGS RNA Library Prep Set (PT042) for the preparation of a 250~300 bp insert cDNA library (unstranded). The library was checked with Qubit and real-time PCR for quantification and bioanalyzer for size distribution detection. Paired end PE150 sequencing was performed on Novaseq6000 short-read sequencing machine (6Gb raw data per sample) platform. Base calling was performed by Novogene using RTA (Real Time Analysis) software. All samples had a Phred score >30.

***RNA sequencing data analysis***

Paired reads were aligned to Ensembl GRCh38.105 2.7.4a [17] using bowtie2 (v2.4.4)[18] default settings, sorted and indexed with Samtools (v1.2)[19], and unstranded exonic read pairs were counted using FeatureCounts (v2.0.7)[20] using a slightly modified Ensembl gtf with 'chr' prefix nomenclature. DESeq2 (v1.44.0)[21] was used in R (v4.4.2) to assess features with >10 reads across conditions and differential expression of transcripts with a fold change >1.5 and FDR <0.05. All other results were given the value 0 and genes with no differential expression were removed from the final data frame for visualisation purposes only.

***Pathway analysis***

Gene Set Enrichment Analysis [22] was performed in R, using clusterProfiler (v4.12.6)[23] on all genes annotated with Entrez IDs using AnnotationDbi (v1.66.0) and org.Hs.eg.db.Hs (v3.19.1) packages. GO Biological Processes [24, 25], Hallmark [26] and Reactome [27] pathway gene sets were downloaded from the Molecular Signature Database (https://www.gsea-msigdb.org/gsea/msigdb) using msigdbr (v7.5.1). Specific sets of genes defining progenitor cell populations, early differentiation and late (terminal) differentiation have been experimentally validated using organotypic models [28]. ‘set.seed’ in R (v4.4.2) was set to ‘1234’ and Benjamini-Hochberg correction (adjusted p-value <0.05) was used for multiple comparison corrections.

***Gene set variation analysis***

A more stringent p-value threshold ($<1\times10^{-8}$) was applied to the SCC13 siYAP/TAZ RNA-seq dataset to reduce the gene set size to <500, as recommended for robust gene set variation analysis (GSVA) [29]. This yielded 486 upregulated and 250 downregulated DEGs in siYAP/TAZ compared to siNTC SCC13 cells. GSVA was performed using the GSVA function (GSVA R package v1.52.3) with default settings. The input comprised the selected SCC13 RNA-seq gene sets and an expression matrix from 237 cSCC patients with known clinical outcome [30], covering primary cSCC (metastatic and non-metastatic), perilesional normal skin and metastasis where available. GSVA enrichment scores were calculated for each sample, and boxplots were used to visualize differential pathway activity across tissue types. T-tests were applied to assess significant differences in enrichment scores between groups.

***Replication fork dynamics analysis by DNAscent***

Base calling of raw sequencing files (pod5 format) and alignment (--mm2-preset map-ont option) were performed using Dorado (version 0.7.3) and the configuration dna_r10.4.1_e8.2_400bps_fast@v5.0.0. Both reads that passed and failed the base calling quality metrics were included in downstream processing. Reads were aligned to the human telomere-to-telomere (T2T) reference genome (chm13v2.0.fa). Next, the output was converted from sam format to bam format using samtools (version 522 1.14). The DNAscent pipeline (version 4.1.1, available on https://github.com/MBoemo/DNAscent) was run with subprogrammes index, detect and forkSense, as previously described [31, 32]. The DNAscent forkSense output (bed file format) was processed with custom Python (version 3.11.5) scripts to obtain fork speeds and stall scores.

***Reproducibility of experiments***

RIME was performed on two independent cell cultures for each cell type, starting from freshly thawed cell stocks. For each fixed cell pellet prepared from the independent cell cultures, YAP RIME was then performed in technical replicates. Control RIME was performed using isotype-matched IgG, and for each independent experiment, the two replicate IgG control immunoprecipitations were pooled for mass spectrometry analysis. For RNA-seq analysis, two independent cell cultures, starting from freshly thawed cell stocks were separately transfected with siRNA SMARTpools. For DNAscent analysis, two independent cell cultures, starting from freshly thawed cell stocks were used. For co-immunoprecipitations two to three independent experiments were performed with similar results. For immunostainings of cells, representative images from one out of three experiments are shown. Western blots are representative of at least two independent experiments. Cell cycle analysis was performed on two to four independent experiments. For clonal growth assays at least three independent experiments were performed with three technical replicates per condition.

***Statistical analysis and data presentation***

Statistical analyses were performed with GraphPad Prism software (version 10.3.1) or R (v7.5.1). No statistical method was used to predetermine sample size. Statistical tests used to determine p-values are specified in Figure Legends. Sample size is indicated in Figure

Legends, Data sets with sufficient *n* numbers were first analysed for normal distribution. In cases where the data were not found to be normally distributed, they were subsequently analysed using appropriate non-parametric tests (specified in Figure Legends), provided that the variance was comparable between the groups analysed. In some figures, superplots [33] were used to communicate both the cell-level variability and the experimental reproducibility. The investigators analysing the data were not blind to the identity of the samples. Further R (v7.5.1) data manipulation packages included genefilter (v1.86.0), dplyr (v1.1.4), tibble (v3.2.1), reshape2 (v1.4.4), plyr (v1.8.9), gage (v2.54.0) [34], gageData (v2.42.0), and rrvgo (v1.16.0), and data were visualised in R using gplots (v3.2.0), ggplot2 (v3.5.1), viridis (v0.6.5), DOSE (v3.30.5) [35], pheatmap (v1.0.12), RColorBrewer (v1.1-3), eulerr (7.0.2), and ggrepel (v0.9.6) packages.

**Supplementary references**


1 Todaro GJ, Lazar GK, Green H. The initiation of cell division in a contact-inhibited mammalian cell line. *J Cell Physiol* 1965; 66: 325-333.

2 Rheinwald JG, Green H. Serial cultivation of strains of human epidermal keratinocytes: the formation of keratinizing colonies from single cells. *Cell* 1975; 6: 331-343.

3 Rheinwald JG. Serial cultivation of normal human epidermal keratinocytes. *Methods Cell Biol* 1980; 21A: 229-254.

4 Walko G, Woodhouse S, Pisco AO, Rognoni E, Liakath-Ali K, Lichtenberger BM *et al*. A genome-wide screen identifies YAP/WBP2 interplay conferring growth advantage on human epidermal stem cells. *Nat Commun* 2017; 8: 14744.

5 Rheinwald JG, Beckett MA. Tumorigenic keratinocyte lines requiring anchorage and fibroblast support cultured from human squamous cell carcinomas. *Cancer Res* 1981; 41: 1657-1663.

6 Proby CM, Purdie KJ, Sexton CJ, Purkis P, Navsaria HA, Stables JN *et al*. Spontaneous keratinocyte cell lines representing early and advanced stages of malignant transformation of the epidermis. *Exp Dermatol* 2000; 9: 104-117.

7 Ichinose Y, Iguchi H, Ohta M, Katakami H. Establishment of lung cancer cell line producing parathyroid hormone-related protein. *Cancer Lett* 1993; 74: 119-124.

8 Totaro A, Castellan M, Battilana G, Zanconato F, Azzolin L, Giulitti S *et al*. YAP/TAZ link cell mechanics to Notch signalling to control epidermal stem cell fate. *Nat Commun* 2017; 8: 15206.

9 De Rosa L, Secone Seconetti A, De Santis G, Pellacani G, Hirsch T, Rothoeft T *et al*. Laminin 332-Dependent YAP Dysregulation Depletes Epidermal Stem Cells in Junctional Epidermolysis Bullosa. *Cell Rep* 2019; 27: 2036-2049 e2036.

10 Mellacheruvu D, Wright Z, Couzens AL, Lambert JP, St-Denis NA, Li T *et al*. The CRAPome: a contaminant repository for affinity purification-mass spectrometry data. *Nat Methods* 2013; 10: 730-736.

11 Schindelin J, Arganda-Carreras I, Frise E, Kaynig V, Longair M, Pietzsch T *et al*. Fiji: an open-source platform for biological-image analysis. *Nat Methods* 2012; 9: 676-682.

12 Stirling DR, Swain-Bowden MJ, Lucas AM, Carpenter AE, Cimini BA, Goodman A. CellProfiler 4: improvements in speed, utility and usability. *BMC Bioinformatics* 2021; 22: 433.

13 Uhlen M, Fagerberg L, Hallstrom BM, Lindskog C, Oksvold P, Mardinoglu A *et al*. Proteomics. Tissue-based map of the human proteome. *Science* 2015; 347: 1260419.

14 Ponten F, Jirstrom K, Uhlen M. The Human Protein Atlas--a tool for pathology. *J Pathol* 2008; 216: 387-393.

15 Melendez Garcia R, Haccard O, Chesneau A, Narassimprakash H, Roger J, Perron M *et al*. A non-transcriptional function of Yap regulates the DNA replication program in Xenopus laevis. *Elife* 2022; 11.

16 Alver RC, Chadha GS, Gillespie PJ, Blow JJ. Reversal of DDK-Mediated MCM Phosphorylation by Rif1-PP1 Regulates Replication Initiation and Replisome Stability Independently of ATR/Chk1. *Cell Rep* 2017; 18: 2508-2520.

17 Harrison PW, Amode MR, Austine-Orimoloye O, Azov AG, Barba M, Barnes I *et al*. Ensembl 2024. *Nucleic Acids Res* 2024; 52: D891-D899.

18 Langmead B, Salzberg SL. Fast gapped-read alignment with Bowtie 2. *Nat Methods* 2012; 9: 357-359.

19 Li H, Handsaker B, Wysoker A, Fennell T, Ruan J, Homer N *et al*. The Sequence Alignment/Map format and SAMtools. *Bioinformatics* 2009; 25: 2078-2079.

20 Liao Y, Smyth GK, Shi W. featureCounts: an efficient general purpose program for assigning sequence reads to genomic features. *Bioinformatics* 2014; 30: 923-930.

21 Love MI, Huber W, Anders S. Moderated estimation of fold change and dispersion for RNA-seq data with DESeq2. *Genome Biol* 2014; 15: 550.

22 Subramanian A, Tamayo P, Mootha VK, Mukherjee S, Ebert BL, Gillette MA *et al*. Gene set enrichment analysis: a knowledge-based approach for interpreting genome-wide expression profiles. *Proc Natl Acad Sci U S A* 2005; 102: 15545-15550.

23 Xu S, Hu E, Cai Y, Xie Z, Luo X, Zhan L *et al*. Using clusterProfiler to characterize multiomics data. *Nat Protoc* 2024; 19: 3292-3320.

24 Gene Ontology C, Aleksander SA, Balhoff J, Carbon S, Cherry JM, Drabkin HJ *et al*. The Gene Ontology knowledgebase in 2023. *Genetics* 2023; 224.

25 Ashburner M, Ball CA, Blake JA, Botstein D, Butler H, Cherry JM *et al*. Gene ontology: tool for the unification of biology. The Gene Ontology Consortium. *Nat Genet* 2000; 25: 25-29.

26 Liberzon A, Birger C, Thorvaldsdottir H, Ghandi M, Mesirov JP, Tamayo P. The Molecular Signatures Database (MSigDB) hallmark gene set collection. *Cell Syst* 2015; 1: 417-425.

27 Milacic M, Beavers D, Conley P, Gong C, Gillespie M, Griss J *et al*. The Reactome Pathway Knowledgebase 2024. *Nucleic Acids Res* 2024; 52: D672-D678.

28 Lopez-Pajares V, Qu K, Zhang J, Webster DE, Barajas BC, Siprashvili Z *et al*. A LncRNA-MAF:MAFB transcription factor network regulates epidermal differentiation. *Dev Cell* 2015; 32: 693-706.

29 Hanzelmann S, Castelo R, Guinney J. GSVA: gene set variation analysis for microarray and RNA-seq data. *BMC Bioinformatics* 2013; 14: 7.

30 Wang J, Harwood CA, Bailey E, Bewicke-Copley F, Anene CA, Thomson J *et al*. Transcriptomic analysis of cutaneous squamous cell carcinoma reveals a multigene prognostic signature associated with metastasis. *J Am Acad Dermatol* 2023; 89: 1159-1166.

31 Jones MJK, Rai SK, Pfuderer PL, Bonfim-Melo A, Pagan JK, Clarke PR *et al*. A high-resolution, nanopore-based artificial intelligence assay for DNA replication stress in human cancer cells. *Nat Commun* 2025; 16: 7732.

32 Lumeau A, Pfuderer P, De Angelis S, Scarth J, Guscott M, Shaikh N *et al*. High levels of DNA replication initiation factors indicate ATRi sensitivity via excessive origin firing. *bioRxiv* 2025: 2025.2002.2027.640046.

33 Lord SJ, Velle KB, Mullins RD, Fritz-Laylin LK. SuperPlots: Communicating reproducibility and variability in cell biology. *J Cell Biol* 2020; 219.

34 Luo W, Friedman MS, Shedden K, Hankenson KD, Woolf PJ. GAGE: generally applicable gene set enrichment for pathway analysis. *BMC Bioinformatics* 2009; 10: 161.

35 Yu G, Wang LG, Yan GR, He QY. DOSE: an R/Bioconductor package for disease ontology semantic and enrichment analysis. *Bioinformatics* 2015; 31: 608-609.
