## Supplementary Table 3 for "YAP engages RIF1 to dampen replication stress-induced DNA damage in human squamous cell carcinoma"

**Supplementary Table 3A – Constitutive shRNAs (Merck Mission®)**

| **Target gene ID** | **Alias** | **Vector** | **Target sequence** | **Clone ID** |
| --- | --- | --- | --- | --- |
| / | shNTC | pLKO.1  (puromycin  -resistant) | / | TCRN0000204151515MN |
| RIF1 | shRIF1-017 |  | TCTTATGAGACGTATAGTATT | TCRN0000220017 |
| RIF1 | shRIF1-022 |  | GCTATCTGGAAGGAGCTAATT | TCRN0000155022 |
| RIF1 | shRIF1-431 |  | CGCATTCTGCTGTTGTTGATT | TCRN0000155431 |
| CDKN1A | shP21-021 |  | GACAGATTTCTACCACTCCAA | TCRN0000287021 |
| CDKN1A | shP21-091 |  | CGCTCTACATCTTCTGCCTTA | TCRN0000287091 |

**Supplementary Table 3B – Inducible shRNAs (Dharmacon™ SMARTvector)**

| **Target gene ID** | **Alias** | **Vector** | **Target sequence** | **Clone ID** | **Lot #** | **Viral titre** |
| --- | --- | --- | --- | --- | --- | --- |
| / | shNTC | piSMART mCMV/  TurboRFP  (puromycin  -resistant) | non-targeting | / | V20012406 | 4.73x107 TU/mL |
| YAP1 | shYAP-1 |  | GCATGAGACAATTTCCATA | V3IHSMCR 6221595 | V20022702 | 9.39x107 TU/mL |
| YAP1 | shYAP-2 |  | CCATATTAGTGAATCTGTT | V3IHSMCR 6830676 | V20022702 | 2.70x107 TU/mL |
| YAP1 | shYAP-3 |  | AAGTGAGCCTGTTTGGATG | V3IHSMCR 10065402 | V20022702 | 9.75x107 TU/mL |
| WWTR1 | shTAZ-1 |  | CAGTCCTATTGTAGCTTAT | V3IHSMCR 6575751 | V20022702 | 2.16x107 TU/mL |
| WWTR1 | shTAZ-2 |  | GGTTTTGATTGAGAGTAAC | V3IHSMCR 8835294 | V20022702 | 9.68x107 TU/mL |
| WWTR1 | shTAZ-3 |  | CGACCTGATTTACAGTTTC | V3IHSMCR 10169649 | V20022702 | 7.66x107 TU/mL |

**Supplementary Table 3C – siRNAs (Dharmacon™ ON-TARGET plus SMARTpool)**

| **Target gene ID** | **Alias** | **Target sequences** |
| --- | --- | --- |
| / | siNTC | UGGUUUACAUGUCGACUAA, UGGUUUACAUGUUGUGUGA, UGGUUUACAUGUUUUCUGA, UGGUUUACAUGUUUUCCUA |
| YAP1 | siYAP | GCACCUAUCACUCUCGAGA, UGAGAACAAUGACGACCA, GGUCAGAGAUACUUCUUAA, CCACCAAGCUAGAUAAAGA |
| WWTR1 | siTAZ | CCGCAGGGCUCAUGAGUAU, GGACAAACACCCAUGAACA, AGGAACAAACGUUGACUUA, CCAAAUCUCGUGAUGAAUC |
| RIF1 | siRIF1 | CCUCAAAUGAAAUGCGAAA, UCACGUAGCCCUAAAUUUA, GAAUCAAAUCUAAGGACUA, GCAAGUUCCUGAUGAUUUA |
| TEAD1 | siTEAD1 | CGAUUUGUAUACCGAAUAA, CACAAGACGUCAAGCCUUU, AAACAGGGAUACACAAGAA, GAAAGGUGGCUUAAAGGAA |
| TEAD3 | siTEAD3 | GGAAGAAGGUGCGGGAGUA, CGCCGACGCUCAGCAGUUA, UUGAUUGCACGCUAUAUUA, GAUCGUCUCUGCCAGUGUC |
| TEAD4 | siTEAD4 | GACAGAGUAUGCUCGCUAU, GGACACUACUCUUACCGCA, UCAAGCACCUCCCUGAGAA, CCCAUGAUGUGAAGCCUUU |

**Supplementary Table 3D – RT-qPCR TaqMan™ assays**

| **Target gene ID** | **Assay ID** | **Dye** |
| --- | --- | --- |
| RIF1 | Hs00871714_m1, cat#4331182 | FAM |
| IVL | Hs00846307_s1, cat#4331182 | FAM |
| GAPDH | Hs02758991_g1, cat#4331182 | FAM |
| GAPDH | Hs02758991_g1, cat#4331182 | VIC |
