## Supplementary Table 4 for "YAP engages RIF1 to dampen replication stress-induced DNA damage in human squamous cell carcinoma"

| **Supplementary Table 4** |  |  |  |  |  |  |  |  |  |
| --- | --- | --- | --- | --- | --- | --- | --- | --- | --- |
| **Primary antibodies** |  |  |  |  |  |  |  |  |  |
| **Antibody information** | | | | | **Dilution** | | | | |
| **Antibody** | **Company/Source** | **Catalog Number/Reference** | **Clone number** | **Description** | **Western Blot** | **IFM-cells** | **IFM-tissues** | **PLA** | **IP** |
| anti-YAP | Santa Cruz Biotechnology | sc-101199 | 63.7 | Mouse monoclonal | 1:500 | 1:50 |  | 1:50 |  |
| anti-YAP | Cell Signaling Technology | 14074 | D8H1X | Rabbit monoclonal | 1:1000 | 1:100 |  | 1:100 | 0.07 µg/IP |
| anti-YAP/TAZ | Cell Signaling Technology | 28287 | D24E4 | Rabbit monoclonal | 1:1000 |  |  |  |  |
| anti-phospho YAP (Ser127) | Cell Signaling Technology | 13008 |  | Rabbit polyclonal | 1:1000 |  |  |  |  |
| anti-YAP | US Biologicals | Y1200-01D |  | Rabbit polyclonal | 1:1000 |  |  |  | 15 µg/IP |
| anti-TAZ | Fisher Scientific | 18668227 | CL0371 | Mouse monoclonal | 1:500 |  |  |  |  |
| anti-RIF1 | Cell Signaling Technology | 95558 | D2F2M | Rabbit monoclonal | 1:1000 | 1:200 | 1:50 | 1:200 |  |
| anti-pan-TEAD | Cell Signaling Technology | 13295 | D3F7L | Rabbit polyclonal | 1:1000 |  |  |  |  |
| anti-TEF1 (TEAD1) | BD Biosciences |  | 31/TEF1 | Mouse monoclonal |  |  |  | 1:100 |  |
| anri-ARID1A | Merck | HPA005456 |  | Rabbit polyclonal | 1:1000 |  |  |  |  |
| anti-CHD4 | Abcam | ab70469 | 3F2/4 | Mouse monoclonal | 1:1000 |  |  |  |  |
| anti-phospho Histone H2A.X (Ser139) | Merck-Millipore | 05-636 | JBW301 | Mouse monoclonal | 1:500 | 1:100 |  | 1:100 |  |
| anti-p21 Waf1/Cip1 | Cell Signaling Technology | 2947 | 12D1 | Rabbit monoclonal | 1:1000 |  |  |  |  |
| anti-CHK1 | Santa Cruz Biotechnology | sc-8408 | G-4 | Mouse monoclonal | 1:500 |  |  |  |  |
| anti-phospho CHK1 (Ser317) | Cell Signaling Technology | 12302 | D12H3 | Rabbit monoclonal | 1:1000 |  |  |  |  |
| anti-phospho CHK1 (Ser345) | Cell Signaling Technology | 2348 | 133D3 | Rabbit monoclonal | 1:1000 |  |  |  |  |
| anti-RAD51 | Abcam | ab133534 | EPR4030(3) | Rabbit monoclonal |  | 1:100 |  |  |  |
| anti-GAPDH | Protein Tech | HRP-60004 |  | Mouse monoclonal | 1:5000 |  |  |  |  |
| anti-CYR61 | Cell Signaling Technology | 14479 | D4H5D | Rabbit monoclonal | 1:1000 |  |  |  |  |
| anti-RPA32 | Santa Cruz Biotechnology | sc-56770 | 9H8 | Mouse monoclonal | 1:500 |  |  |  |  |
| anti-phospho-RPA32/RPA2 (Ser33) | Cell Signaling Technology | 3099645 | E9N1T | Rabbit monoclonal | 1:1000 |  |  |  |  |
| anti-phospho RPA32/RPA2 (Ser8) | Cell Signaling Technology | 54762 | E5A2F | Rabbit monoclonal | 1:1000 |  |  |  |  |
| anti-phospho RPA32/RPA2 (Ser4 & Ser8) | Abcam | ab243866 | BL-165-5F1 | Rabbit monoclonal | 1:1000 |  |  |  |  |
| anti-phospho MCM2 (Ser53) | Abcam | ab70367 |  | Rabbit polyclonal | 1:1000 |  |  |  |  |
| anti-MCM4 | Santa Cruz Biotechnology | sc-28317 | G-7 | Mouse monoclonal | 1:1000 |  |  |  |  |
| anti-Histone H3 | Abcam | ab1791 |  | Rabbit polyclonal | 1:1000 |  |  |  |  |
| anti-PCNA | Cell Signaling Technology | 2586 | PC10 | Mouse monoclonal | 1:1000 |  |  |  |  |
| anti-Keratin 14 | Biolegend | 906004 | Poly9060 | Chicken polyclonal |  |  | 1:250 |  |  |
| Cell Cycle (pCdk/pHH3/Actin) cocktail | Abcam | ab136810 |  | Rabbit monoclonal | 1:1000 |  |  |  |  |
| Cyclophilin B | Cell Signaling Technology | 43603 | D1V5J | Rabbit monoclonal | 1:1000 |  |  |  |  |
| anti-α-Tubulin | Merck | T6199 | DM1A | Mouse monoclonal | 1:2000 |  |  |  |  |
| anti-α-Tubulin | Cell Signaling Technology | 3873 | DM1A | Mouse monoclonal | 1:1000 |  |  |  |  |
| anti-beta-Actin | Merck |  | AC-40 | Mouse monoclonal | 1:500 |  |  |  |  |
| anti-CDK2 | Cell Signaling Technology | 2546 | 78B2 | Rabbit monoclonal | 1:1000 |  |  |  |  |
| anti-CDK4 | Cell Signaling Technology | 12790 | D9G3E | Rabbit monoclonal | 1:1000 |  |  |  |  |
| anti-CDK6 | Cell Signaling Technology | 3136 | DCS83 | Rabbit monoclonal | 1:1000 |  |  |  |  |
| Rabbit mAb IgG XP® Isotype Control | Cell Signaling Technology | 3900 | DA1E | Rabbit monoclonal |  |  |  |  | 0.07 µg/IP or 15 µg/IP |

| **Secondary antibodies** |  |  |  |  |  |  |  |  |  |
| --- | --- | --- | --- | --- | --- | --- | --- | --- | --- |
| **Antibody information** | | | | | **Dilution** | | | | |
| **Antibody** | **Company/Source** | **Catalog Number/Reference** | **Clone number** | **Description** | **Western Blot** | **IFM-cells** | **IFM-tissues** | **PLA** | **IP** |
| Anti-rabbit IgG, HRP-linked Antibody | Cell Signaling Technology | 7074 |  | Goat anti-rabbit IgG | 1:2000 |  |  |  |  |
| Anti-mouse IgG, HRP-linked Antibody | Cell Signaling Technology | 7076 |  | Goat anti-mouse IgG | 1:2000 |  |  |  |  |
| Mouse anti-rabbit IgG (conformation specific) | Cell Signaling Technology | 3678 | L27A9 | Mouse anti-rabbit IgG | 1:2000 |  |  |  |  |
| Rabbit anti-mouse IgG (light chain specific) | Cell Signaling Technology | 58802 | D3V2A | Rabbit anti-mouse IgG | 1:1000 |  |  |  |  |
| Goat anti-rabbit IgG (H+L), highly cross-adsorbed, Alexa Fluor™ Plus 555 | Thermo Fisher Scientific | A32732 |  | Goat anti-rabbit IgG |  | 1:1000 |  |  |  |
| Goat anti-mouse IgG (H+L), highly cross-adsorbed, Alexa Fluor™ Plus 488 | Thermo Fisher Scientific | A32723 |  | Goat anti-mouse IgG |  | 1:1000 |  |  |  |
| Donkey anti-chicken IgY (H+L), highly cross adsorbed, Alexa Fluor™ 594 | Thermo Fisher Scientific | A78951 |  | Donkey anti-chicken IgY |  |  | 1:500 |  |  |
| Donkey anti-rabbit IgG (H+L), highly cross-adsorbed, Alexa Fluor™ 488 | Thermo Fisher Scientific | A21206 |  | Donkey anti-rabbit IgG |  |  | 1:500 |  |  |
